# MRI-compatible 3D Soft Carbon Neural Interface Enabled by Versatile Depth-tunable Laser Drawing

**DOI:** 10.64898/2026.10.19.760266

**Authors:** Xuanqi Wang, Honglong Chang, Feng Yue, Bowen Ji

## Abstract

Soft neural interfaces are designed to minimize mechanical mismatch with the brain. However, planar and recessed electrode sites remain locally decoupled from the curved and pulsating cortical surface, degrading the fidelity of neural sensing. Here, we report a 3D soft carbon electrode array, enabled by a depth-tunable laser drawing strategy. This laser-driven workflow achieves versatile fabrication of customizable 3D soft carbon neural interfaces, eliminating the reliance on conventional thin-film microfabrication processes. The fully silicone-based electrode structure realize device-scale conformality, and the 3D soft carbon sites promote a non-penetrating, intimate neural-electrode interface. The CNT-3D array combined superior in-vitro electrochemical and in-vivo recording performance compared with Au-2D and CNT-2D controls. Furthermore, in bilateral rabbit cortex implantation, CNT-3D array retained 80.4% of electrocorticography power after 4 months, versus 11.7% for CNT-2D control. Moreover, the 3D sensing sites significantly suppressed line-frequency harmonics and background noise interference. Subsequently, during magnetic resonance imaging, the CNT-3D array exhibits negligible implant-induced artifacts compared to the conventional Pt-Ir metal array. Overall, this work establishes a laser drawing strategy that extends neural interfaces from macro device-scale conformality to micro site-scale coupling, providing a technological pathway for practical and translational soft neurotechnology.

## Introduction

Minimally invasive neural interfaces require stable cortical coupling^1^, tissue-matched mechanics^2,3^, and compatible postoperative magnetic resonance imaging (MRI)^4,5^ to support reliable clinical translation^6^. Flexible polymer films and soft elastomers have been widely adopted to improve conformal integration with complex and dynamic neural surfaces^7–10^. However, the variable cortex curvature, constant physiological micromotion, and pulsatile cerebrospinal fluid can compromise local electrode-tissue apposition and introduce variability in interfacial coupling^2,11,12^. Device-scale conformality does not necessarily ensure intimate and mechanically stable contact at every individual recording site, particularly when the contacts are recessed relative to the encapsulation surface^13,14^. Planar sensing sites lead to locally weakened coupling from the cortex, substantially degrading interfacial sensing fidelity and increasing noise interference susceptibility^15,16^. Therefore, a significant requirement is to translate macro device-scale conformality into micro site-scale coupling at the neural-electrode interface.

Advances in soft and mechanically compliant implantable neural interfaces have substantially supported the practicality for motor function recovery^8,17^, hearing loss rehabilitation^10^, speech decoding^18,19^, and vision restoration^20,21^. These studies establish intrinsically compliant materials and architectures as effective strategies for advancing neural interfaces toward reliable clinical translation. Building on these advances, artifact-minimized postoperative MRI is increasingly needed to assess implant positioning and monitor surrounding neural tissue during long-term implantation^22,23^, motivating recent soft implant researches to replace conventional metallic conductors with low-susceptibility materials^4,24^. However, current soft neural interfaces remain largely constrained to planar sensing architectures, maintaining intimate site-scale coupling while integrating compliant mechanics and compatible MRI remains a design challenge.

This challenge motivates the geometry redesign of the local sensing sites exposed above the soft silicone encapsulation surface^4,13,25^. Beyond material softness, three-dimensional (3D) spatial organization can introduce sensing capabilities that are difficult to realize with planar layouts^26^. Protruding 3D electrode sites offers a promising non-penetrating approach for superior neural sensing fidelity^16,27^. In this electrode architecture, the 3D feature establishes an intimate and active neural-electrode interface that eliminates passive micro gaps. Such site-scale coupling is expected to reduce the impedance, minimize interference, and enhance cortical signal fidelity from the complex in vivo environment.

However, the fabrication of 3D neural interfaces based on deformable soft elastomers remains technically endeavor. Conventional micro-electro-mechanical system (MEMS) microfabrication is optimized for planar layouts, whereas creating independently tunable out-of-plane microgeometries typically requires additional molding, printing, forming or selective-opening steps, complicating integration with soft substrates^13,25,28^. Moreover, metallic conductors remain heavily reliant in conventionally microfabricated electrodes, creating an inherent material-level conflict with MRI^29–31^. The pronounced magnetic susceptibility mismatch between metallic layers and surrounding tissues induces local magnetic-field inhomogeneity, leading to image distortion^5,32^. Meanwhile, the metal electrodes has to face fatigue, corrosion, and even delamination during chronic implantation^6,33,34^. Consequently, the integration of 3D soft neural interfaces and fully metal-free electrical functionality demands a fundamentally reengineered fabrication strategy beyond the conventional MEMS processes.

Here, we report a 3D soft carbon electrode array, bridging device-scale conformality and site-scale cortical coupling, while producing negligible MRI artifacts. This array is fabricated by a depth-tunable laser drawing strategy with sacrificial PVA/PMMA dual templates. The spatially selective laser penetration enables simultaneous patterning of the interconnects, protruding 3D sites and pads in soft silicone matrix, followed by the backfill of the CNTs-embedded silicone conductive network. This method enables efficient and customizable electrode fabrication compared to MEMS processes. We next systematically evaluated the electrochemical and mechanical performance of the resulting electrodes. Furthermore, bilateral implantation in rabbit cortex enabled a direct comparison between the fabricated CNT-3D electrodes and CNT-2D controls, isolating the advantage of 3D sites through in vivo impedance measurements and chronic electrocorticography (ECoG) recordings. Finally, 5.0 T MRI was used to compare postoperative imaging compatibility between the CNT-3D array and Pt-Ir metal array. Collectively, the laser drawing strategy establishes scalable soft 3D carbon neural implants toward MRI-compatible minimally invasive brain-computer interface.

## Results

### 3D soft carbon neural interface

Figure 1a illustrates the micro site-scale coupling interface between the protruding 3D carbon micro sites and the cortex. The soft conical geometry was designed to promote non-penetrating and intimate cortical contact, as well as extend the active surface beyond the silicone substrate. Additionally, The ECoG waveforms reveal a pronounced improvement from lower-amplitude, noise-contaminated recordings at the planar interface to higher-fidelity, noise-suppressed signals at the 3D soft CNT interface (Fig. 1a).

**Fig. 1.**
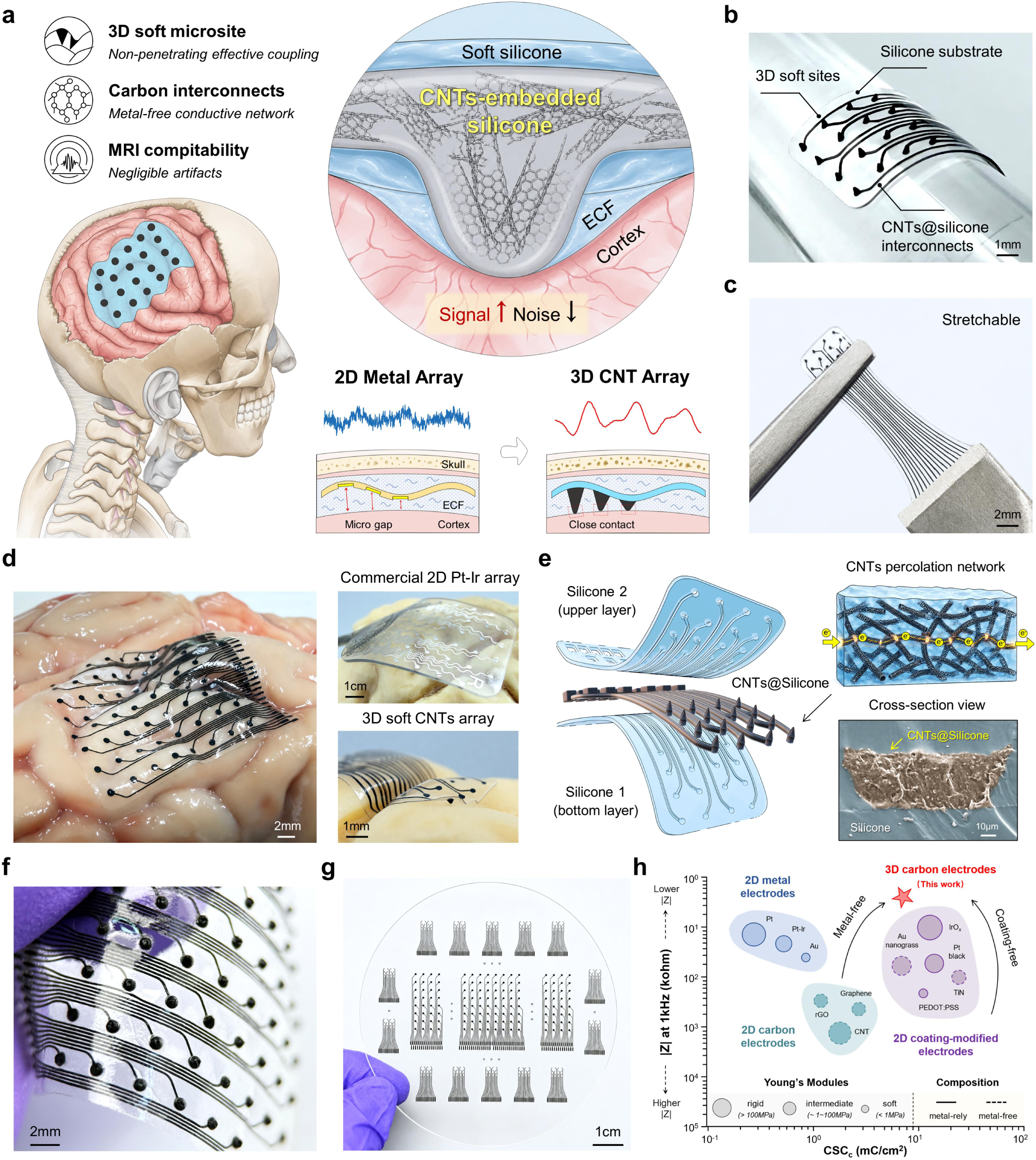
Design of the 3D soft carbon electrode array. **a,** Schematic of the site-scale coupling at the electrode-tissue interface. Representative ECoG recordings illustrate the transition from noisy, low-amplitude waveforms to high-fidelity neural signals. **b,** Photograph of the 16-channel array conformally wrapped around a glass rod, showing the silicone substrate, embedded CNTs@silicone interconnects and protruding 3D micro sites. **c,** Photograph of this array under stretching deformation. **d,** The 64-channel array conforming to the sulci and gyri on porcine cortex. Right images provide a microscopic comparison of the commercial 2D Pt-Ir array and the 3D soft CNT array. **e,** Exploded model and cross-sectional SEM image of the CNT array, illustrating the interconnected CNTs network within the elastomer matrix. **f,** Magnified photograph of the 64-channel array. **g,** Scalable fabrication of the 3D soft CNT arrays. **h,** Performance benchmark comparing 2D metal, 2D carbon, 2D coating-modified and 3D carbon electrodes, showing simultaneous low impedance and high CSCc achieved by this work.

As shown in Figure 1 b, we integrated the 3D sensing sites and conductive pathways within a fully soft elastomeric architecture. This electrode array comprising the silicone elastomer body, and the carbon nanotubes (CNTs)-embedded silicone traces with 3D microsites (Fig. 1e). It has been established that the carbon traces had been inserted between the layers of silicone encapsulation as confirmed by cross-sectional scanning electron microscopy (SEM) and elemental mapping (Fig. 1e and Supplementary Fig. 3). Within the CNTs@Silicone composite, crosslinked CNTs formed interconnected percolation pathways through physical contacts and tunnelling junctions, providing electrical conduction throughout the elastomeric matrix without metallic materials (Supplementary Fig. 2). The 3D soft carbon array retained mechanical stability, allowing it to undergo cyclic stretch, twist, press and rub deformation without structural damage and functional failure (Fig. 1c and Supplementary Fig. 4).

Subsequently, we examined the device-scale conformality at the physical brain surface. The 64-channel array followed the gyri, sulci, and local curvature of the porcine cortex (Fig. 1d, left). At the local site-scale interface, the magnified comparison with a commercial planar platinum-iridium (Pt-Ir) array revealed a marked difference in cortical apposition (Fig. 1d, right, Supplementary Fig. 1). Whereas the commercial array decoupled from the exposed cortical surface, while the 3D carbon array closely followed the local tissue contour, without an obvious interfacial gap. A subsequent magnified view revealed the 64-channel 3D sites and the embedded CNTs@Silicone interconnects integrated within the transparent silicone substrate (Fig. 1f and Supplementary Fig. 5). Collectively, these observations establish a dual-scale cortical interface in which the conformal array accommodates macroscopic cortical topology while the protruding 3D carbon sites promote intimate physical apposition at individual sensing locations.

This 3D soft carbon electrode was extended across multiple channel counts and array footprints. On the 4-inch PMMA substrate, one 64-channel array, two 32-channel arrays and fourteen 16-channel arrays were fabricated in parallel, demonstrating the scalability across device dimensions and channel numbers (Fig. 1g). Benchmarking against representative ECoG electrodes positioned the present device within a favorable electrochemical-mechanical performance space, combining low impedance with high cathodic charge storage capacity (CSC_c_) (Fig. 1h and Supplementary Table 1).

### Versatile depth-tunable laser drawing fabrication

To fabricate the soft multilayer carbon array with out-of-plane microtopography, we developed a depth-tunable laser drawing strategy based on the PVA/silicone/PMMA stack (Fig. 2a and Supplementary Fig. 7). Instead of functioning as simple through-cutting, this laser drawing process combines CAD-defined lateral patterning with region-specific ablation depths to encode different functional structures within the stack. For the interconnect regions, the laser is employed to specifically penetrate through the PVA mask, while preserving the underlying silicone substrate. While at the electrode site and the contact pad regions, penetration extends through the PVA and silicone layers into the PMMA mold to generate 3D micro cavities. (Fig. 2a and Supplementary Fig. 8a-c). By controlling laser parameters such as power, frequency, and path, we achieve depth-tunable formation of the 3D micro cavities.

**Fig. 2.**
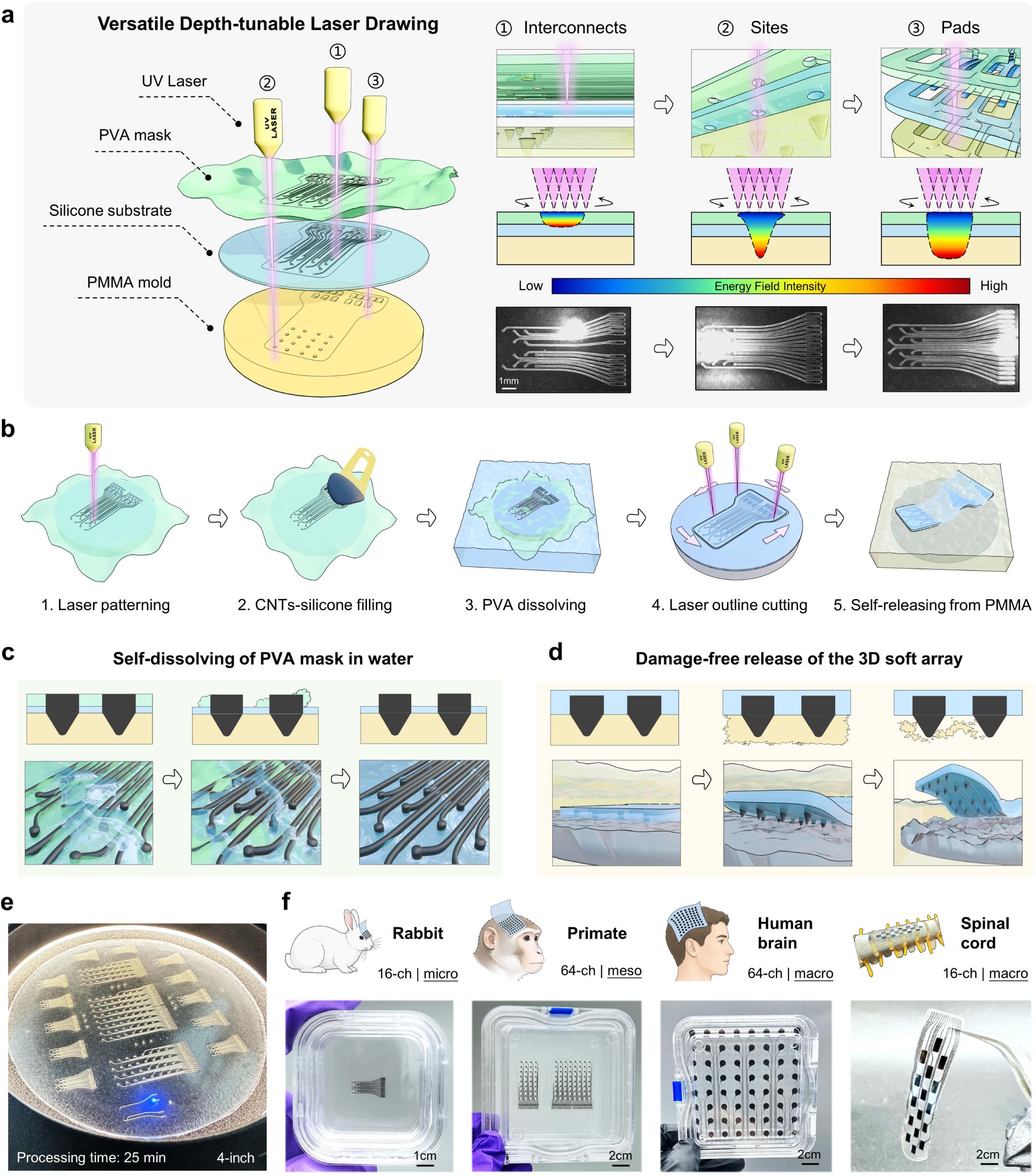
Laser drawing strategy for versatile neural interfaces fabrication. **a,** Schematic of the depth-tunable laser drawing process. Ultrafast UV laser simultaneously patterns the PVA mask, silicone substrate and PMMA mold with region-specific tunable penetration depths. **b,** Key processes of the laser-driven fabrication workflow. **c,** Automatic removal of the water-soluble PVA mask without mechanical peeling. **d,** Automatic release of the formed array in acetone without mechanical demolding. **e,** Photograph of the laser drawing process, corresponding to the electrode arrays shown in Fig. 1g. **f,** Representative electrodes including microscale 16-channel rabbit array, mesoscale 32/64-channel primate array, macroscale 64-channel human brain array, and 16-channel spinal cord array.

As Fig. 2b shows, the laser-patterned channels and micro cavities were subsequently backfilled with the CNTs-embedded silicone conductive composite, which was prepared by uniformly dispersing multi-walled carbon nanotubes of varying lengths in silicone (see the Method section). The PVA layer functioned as a water-soluble patterning mask, confining the conductive CNTs@Silicone to the laser-drawn interconnect, site, and pad regions (Fig. 2c). After the curing of the carbon composite, the stack is to be immersed in deionized water. This will cause the PVA mask to self-dissolve and the formed carbon structures to become exposed without mechanical peeling, thereby preserving the dimensional integrity of the soft embedded carbon layer (Supplementary Fig. 9). Conversely, the underlying PMMA functioned as both the supporting carrier and the 3D mold that defined the out-of-plane micro cavities (Fig. 2d). Following the silicone encapsulation and laser outline cutting processes, the dissolution of PMMA in acetone resulted in the release of the formed electrode array without mechanical demolding, thereby minimizing the damage to the protruding 3D microstructures (Supplementary Fig. 10). Consequently, the two sacrificial layers exhibited complementary functions: PVA facilitated the delineation of conductive patterns and the removal of non-mechanical masks, while PMMA enabled 3D shaping and dissolution-assisted device release.

The laser-drawing strategy could be replicated across multiple electrode arrays on a common 4-inch PMMA substrate, enabling rapid wafer-scale batch fabrication (Fig. 2e). A 3 × 3 layout comprising nine 9-channel arrays was completed in just 15 minutes (Supplementary Fig. 8). Notably, this fabrication workflow is suitable across 2-inch to 8-inch wafer dimensions. And the scaling of the process did not necessitate alterations to the underlying material stack or the laser-drawing principle, thereby establishing a shared manufacturing foundation for versatile array layouts.

As Figure. 2f shows, the laser-drawing strategy enabled the reconfiguration of the electrode geometry, channel count, and overall device format for different dimensions of neural interface. Representative cortical devices included a 16-channel microscale array for rabbit and 32/64-channel mesoscale arrays for primate. The array was further extended to a macroscale electrode format for human brain, with larger sensing sites and compliant serpentine interconnects (Fig. 2f and Supplementary Fig. 11). Moreover, a 16-channel spinal cord electrode array further demonstrated the versatility of the laser-drawing principle, showing its capacity to adapt to geometries tailored to anatomically distinct central nervous system interfaces (Fig. 2f).

### Superior electrochemical performance

Having established the fabrication strategy, we next examined the influence of material and 3D geometry on the electrode-electrolyte interface under controlled electrochemical conditions (Fig. 3a). Three electrode architectures were evaluated: a planar Au array (Au-2D), a planar CNTs@silicone array (CNT-2D) and a protruding CNTs@silicone array (CNT-3D). These electrodes provide complementary controls for distinguishing between material and microstructure contributions (Fig. 3a and Supplementary Fig. 13). Comparison between the Au-2D and CNT-2D arrays primarily assessed the influence of replacing a conventional metallic interface with the CNTs@silicone composite while retaining a planar site geometry. In contrast, CNT-2D and CNT-3D arrays shared the same carbon conductive composite and differed principally in sensing-site geometry, providing the key same-material control for evaluating the contribution of 3D conical structure. All three arrays were characterized under the same measurement conditions, enabling a comparison of cyclic voltammetry (CV), electrochemical impedance spectroscopy (EIS), and voltage transient responses on a common basis (Supplementary Fig. 14).

**Fig. 3.**
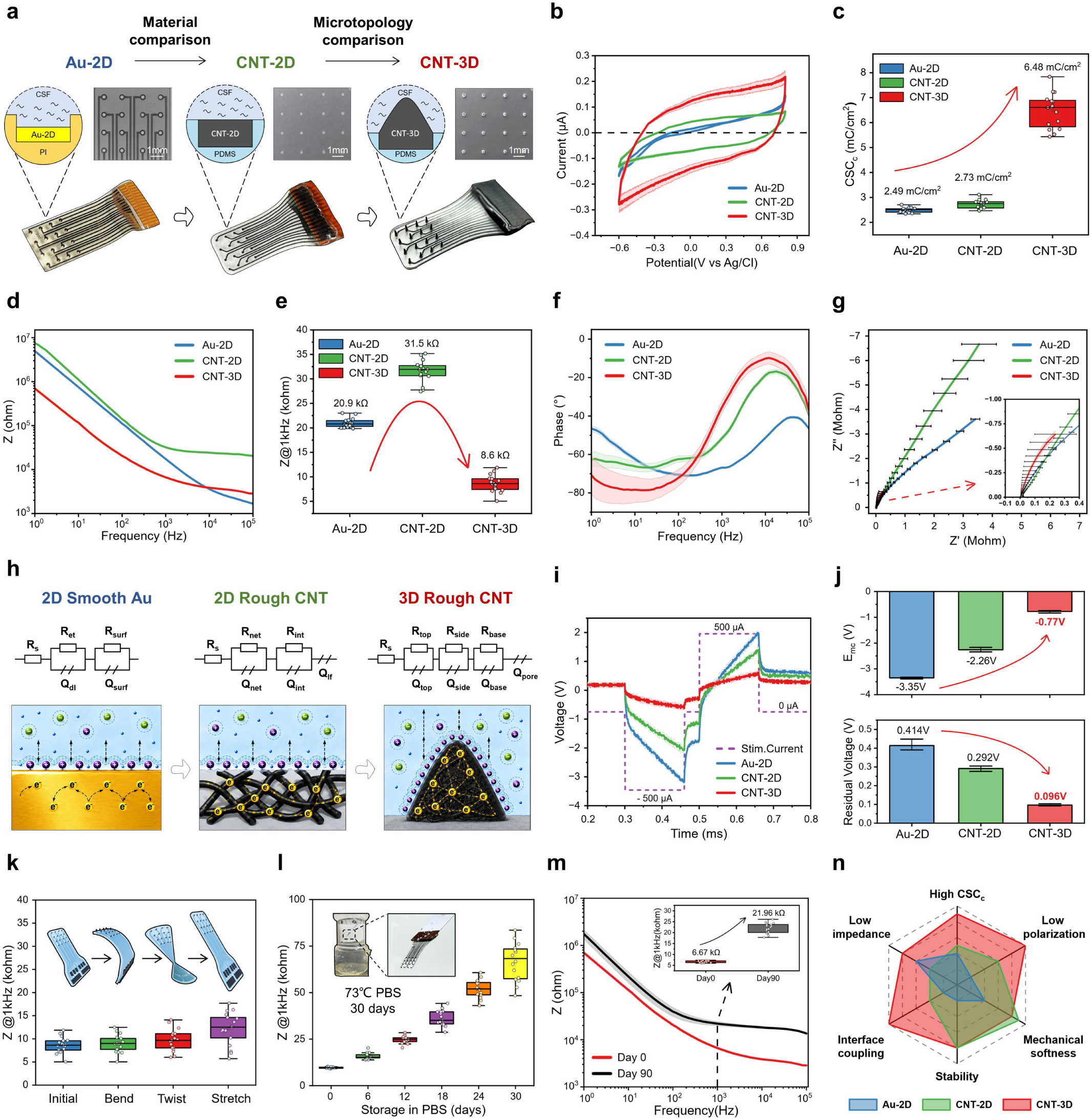
Comparison of the electrochemical performance. **a,** Photographs, schematics, and SEM images of the 16-channel Au-2D, CNT-2D and CNT-3D arrays. **b,c,** Cyclic voltammetry curves and quantified CSCc. **d-g,** Electrochemical impedance spectra curves, quantified 1 kHz impedance, phase spectra curves and Nyquist plots. **h,** Equivalent circuit models of the above three electrode-electrolyte interfaces. **i,j,** Voltage transient response curves and quantified stimulation metrics under identical charge-balanced biphasic current pulses. **k-m,** Impedance stability after bending, twisting and stretching deformation (1,000 cycles), accelerated aging, and 3 months in vivo implantation. **n,** Radar plot summarizing the superiority of CNT-3D array, relative to Au-2D and CNT-2D controls.

The results of the CV test demonstrated that the CNT-3D electrodes exhibited a substantially enlarged cathodic voltametric response in comparison with both Au-2D and CNT-2D planar controls (Fig. 3b and Supplementary Fig. 15a). Quantification of the CSC_c_ yielded 2.49 mC cm⁻² for Au-2D, 2.73 mC cm⁻² for CNT-2D, and 6.48 mC cm⁻² for CNT-3D, corresponding to 2.6-fold and 2.4-fold increases, respectively (Fig. 3c and Supplementary Fig. 15b,c). Notably, the modest difference between Au-2D and CNT-3D is contrasted by the pronounced enhancement from CNT-2D to CNT-3D, supporting a substantial contribution from the protruding 3D geometry. The higher CSC_c_ is consistent with an increased electrochemically accessible electrode-electrolyte interface^27^. This, in turn, improve the amount of charge that can be reversibly accommodated.

Meanwhile, the EIS revealed a significant decrease in electrode-electrolyte interface for CNT-3D array (Fig. 3d and Supplementary Fig. 16). At 1 kHz frequency, the impedance magnitude decreased to approximately 8.6 kΩ for CNT-3D, in comparison with 31.5 kΩ for CNT-2D and 20.9 kΩ for Au-2D (Fig. 3e). The phase spectra demonstrated that the observed reduction was concomitant with a distinct frequency-dependent polarization response, rather than a uniform decrease in impedance magnitude alone (Fig. 3f). The Nyquist plots demonstrated the complex-impedance trajectories, indicating disparities in the interfacial dynamics of the three arrays (Fig. 3g). This interpretation is further substantiated by equivalent-circuit analysis, which revealed that Au-2D is characterized by effective polarization at the planar Au-electrolyte interface, while CNT-2D incorporates contributions from planar CNT-electrolyte interface and low-frequency polarization. In contrast, CNT-3D captures spatially distributed responses associated with the cone top, sidewall, and base/pore regions (Fig. 3h and Supplementary Fig. 17). Collectively, these results suggest that the 3D micro geometry facilitates more than merely increasing the electrochemically accessible area. It redistributes the electrode-electrolyte response across multiple interfacial regions, thereby providing a mechanistic basis for the reduced impedance of 3D sensing sites.

Therefore, under the same identical charge-balanced biphasic current pulses, the three electrodes exhibited markedly different voltage-transient responses, with CNT-3D showing the smallest voltage excursion (Fig. 3i). Moreover, the peak-to-peak voltage decreased from 5.15 V for Au-2D, and 3.44 V for CNT-2D to 1.67 V for CNT-3D, indicating substantially reduced voltage demand and polarization during the imposed current pulse (Fig. 3j and Supplementary Fig. 18a). Furthermore, CNT-3D exhibited a more rapid return to the baseline following pulse termination, with a recovery time of only 0.02 ms, compared with 0.31 ms for CNT-2D and 0.41 ms for Au-2D (Supplementary Fig. 18b). The smallest voltage excursion and fastest post-pulse recovery are consistent with a more efficient interfacial response of the CNT-3D electrode during current delivery.

Subsequently, the stability of these electrochemical advantages under repeated mechanical deformation and chronic implantation-relevant exposure was assessed. The CNT-3D array demonstrated electrical integrity over 1,000 cycles of large repeated bending, twisting, and stretch, consistent with the deformation-tolerant CNTs@silicone structure (Fig. 3k and Supplementary Fig. 20). Long-term stability was further evaluated by subjecting the samples to accelerated aging in 73°C Phosphate Buffered Saline (PBS) for 30 days, corresponding to 364 days of implantation at physiological temperature. During this period, the mean 1 kHz impedance remained below 100 kΩ, with no occurrence of open-circuit failure or abrupt impedance increase across the 16-channel array (Fig. 3l and Supplementary Fig. 19). Furthermore, over the course of 120 days cortical implantation in rabbit, the electrical interfacing of CNT-3D arrays also maintained impedance stability (Fig. 3m). Collectively, as illustrated in Figure. 3n, a comprehensive summary of the three electrodes is provided, including CSC_c_, impedance, degree of polarization induced by electrical stimulation, mechanical softness, chronic stability, and interface coupling. Notably, CNT-3D demonstrates a more expansive multidimensional performance envelope in comparison to Au-2D and CNT-2D.

### Improved cortical recordings and in vivo impedance

After demonstrating the electrochemical advantages of CNT-3D array, we evaluated the cortical signal acquisition capability of the Au-2D, CNT-2D, and CNT-3D electrode arrays (Fig. 4a). Square-wave electrical stimulation was applied to the forelimbs of rats, and cortical electrical signals were recorded using the three arrays (Fig. 4a and Supplementary Fig. 21). CNT-3D array exhibited more pronounced cortical signals together with stronger stimulus-associated time-frequency activity, providing initial functional evidence of the improved acquisition performance enabled by CNT-3D array (Fig. 4b,c).

**Fig. 4.**
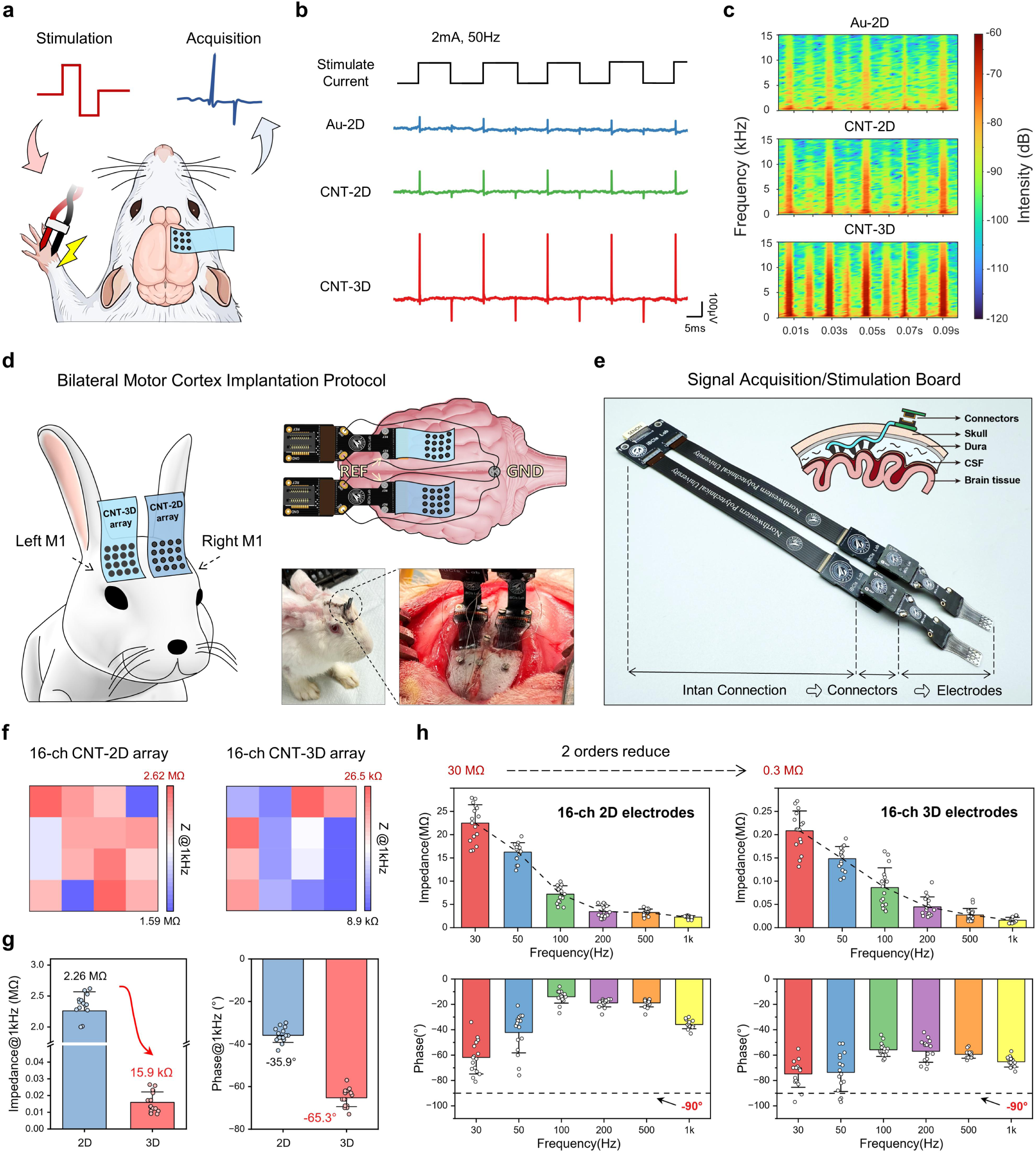
Comparison of cortical recordings and in vivo impedance. **a,** Acute forepaw stimulation-acquisition paradigm, evaluating cortical recordings with Au-2D, CNT-2D, and CNT-3D arrays. **b,c,** Representative recordings and corresponding time-frequency spectrograms. **d,** Schematics and photographs of the bilateral motor cortex region implantation protocol based on a New Zealand white rabbit. **e,** Image of the designed signal acquisition/stimulation board for simultaneous bilateral characterization. **f,** Spatial maps of 1 kHz in vivo impedance across 16 channels of the implanted arrays. **g,h,** Quantified impedance and phase characteristics. CNT-3D electrodes exhibited markedly reduced in vivo impedance and a more capacitive electrode-electrolyte interface.

To further validate the contribution of the protruding 3D sites under physiological in vivo cortical conditions, we next designed a bilateral motor cortex (M1) implantation comparison experiment in New Zealand white rabbit. As Figure 4d and supplementary Figure 23 illustrates, the CNT-2D and CNT-3D arrays were implanted over bilateral M1 regions of the rabbit. This experiment design aims to minimize material, inter-animal, and acquisition-system-related confounds, facilitating a controlled comparison of planar and protruding sensing-site geometries. Both arrays were connected to the customized bilateral signal acquisition/stimulation system, enabling matched characterization within the same experimental session (Fig. 4e and Supplementary Figs. 22 and 23).

Subsequently, the in vivo interface impedance magnitude and phase were measured from the applied current and the corresponding voltage response (Supplementary Fig. 24). Spatial maps of the 1 kHz in vivo impedance revealed a pronounced array-wide separation between the two implanted interfaces across the full 4 × 4 layouts (Fig. 4f). Quantitatively, the mean 1 kHz in vivo impedance decreased from 2.26 MΩ for CNT-2D to 15.9 kΩ for CNT-3D, corresponding to an order of magnitude reduction (Fig. 4g). Notably, this separation was substantially greater than the 3.7-fold difference measured between CNT-2D and CNT-3D in PBS. This finding indicates that the geometry of the sensing site exerts a substantial influence on the effective cortical interface under physiological conditions, particularly in the presence of blood or biological tissue (Figs. 3e and 4g). The marked and array-wide reduction of in vivo impedance supports substantially improved effective electrode-brain interface enabled by the protruding 3D sensing sites. Furthermore, we ascertained the impedance magnitude and phase responses of the array at multiple frequency points. Similarly, CNT-3D exhibited significantly lower in vivo impedance magnitude compared to CNT-2D (Fig. 4g,h). The corresponding phase spectra revealed distinct frequency-dependent interfacial behavior between the two electrodes, with the overall phase of CNT-3D being closer to −90°, exhibiting a stronger capacitive response than the planar CNT-2D interface (Fig. 4h).

### Advanced chronic ECoG fidelity

The subsequent investigation focused on assessing whether the CNT-3D interface led to enhanced ECoG recording fidelity during chronic implantation. As demonstrated in Figure 5a, the improved cortical coupling would facilitate much better acquisition of physiologically significant cortical field potentials. This is achieved by decreasing electrode-tissue separation and limiting signal attenuation or shunting through intervening fluid layer. The enhancement of the interface would also serve to mitigate the vulnerability to interference arising from unstable or spatially diffuse coupling pathways. Furthermore, the 3D soft sensing sites would maintain more effective cortical coupling over time compared to 2D controls. Consequently, the recording advantage of CNT-3D array would be better preserved during chronic implantation rather than being confined to the early postoperative period.

**Fig. 5.**
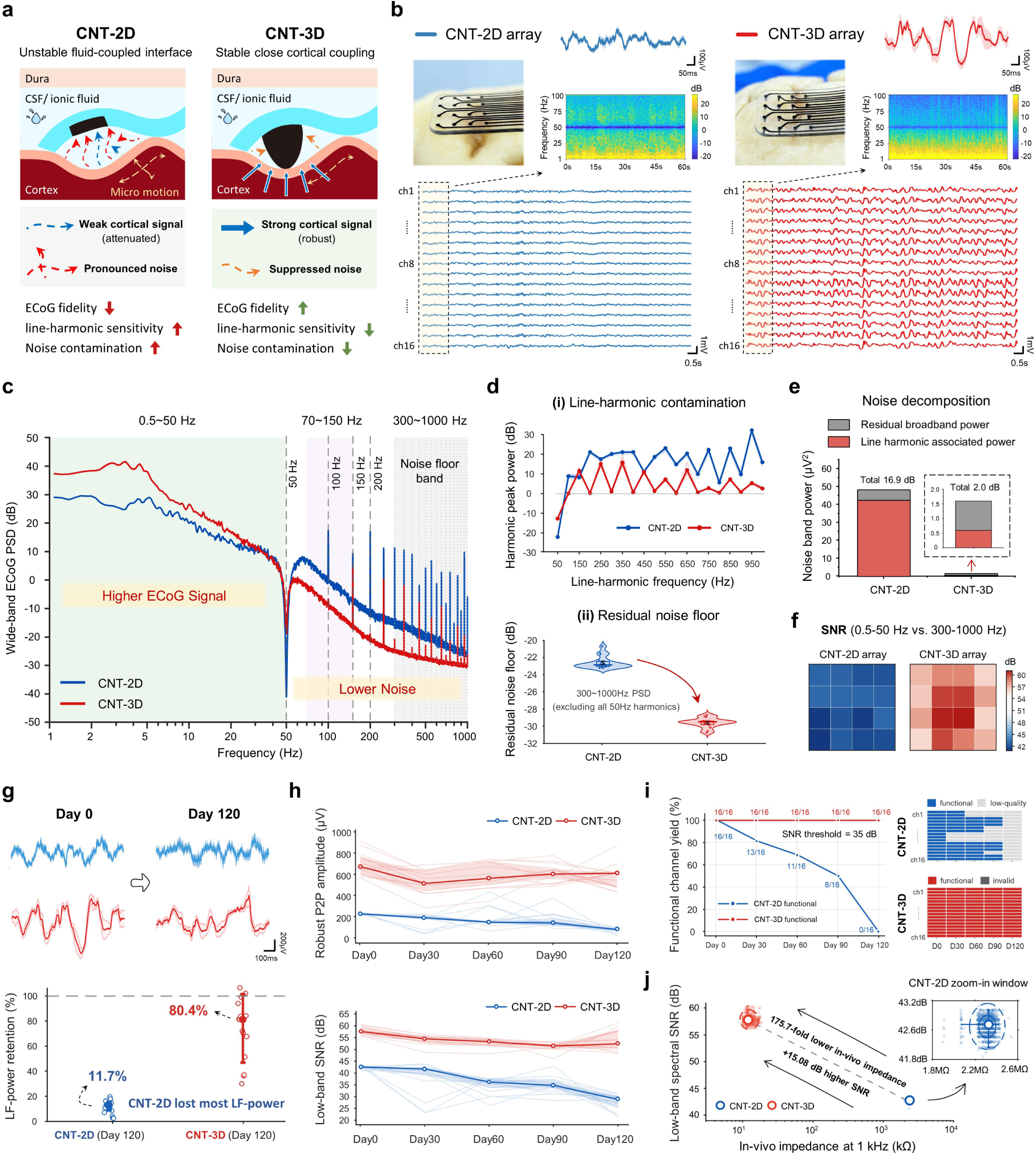
ECoG recordings of CNT-2D and CNT-3D arrays. **a,** Conceptual comparison of the electrode-cortex interfaces, illustrating the 3D micro site reduces cerebrospinal-fluid separation, microscale interfacial gaps and motion-induced instability. **b,** Photos and representative ECoG waveforms. CNT-3D produced larger and cleaner ECoG signals with more consistent multichannel responses. **c,** Mean power spectral density (PSD) curves. CNT-3D preserved stronger low-frequency cortical activity while suppressing high-frequency interference. **d,** Quantification of line-harmonic interference and residual broadband noise. **e,** Decomposition of high-frequency power into harmonic-associated and residual broadband components, showing lower contributions from both sources in CNT-3D recordings. **f,** Spatial maps of low-band spectral SNR across the 4 × 4 arrays, demonstrating improved recording uniformity for CNT-3D array. **g,** Representative array-averaged recordings on Days 0 and 120 and the corresponding low-frequency power retention. **h,** Chronic robust peak-to-peak amplitude and low-band spectral SNR. **i,** Functional channel yield over 120 days (SNR threshold: 35 dB). **j,** Joint analysis of impedance and SNR, showing a distinct low-impedance, high-signal and low-noise operating regime for CNT-3D arrays.

Postoperative simultaneous recordings from the bilaterally implanted arrays demonstrated visibly stronger cortical fluctuations in CNT-3D across 16-channel ECoG recordings (Fig. 5b). Spectral analysis further localized this difference predominantly to the physiologically relevant low-frequency range, with CNT-3D exhibiting greater spectral power than CNT-2D over 0.5-50 Hz (Fig. 5c). Consistent with this observation, independent quantification of the low-frequency recordings showed higher RMS amplitude, robust peak-to-peak amplitude and mean spectral power for CNT-3D, supporting a consistent enhancement across complementary signal-amplitude metrics (Supplementary Fig. 30).

However, the elevated high-frequency power observed in CNT-2D recordings, raised the question of whether this component represented genuine physiological high-gamma activity or interface-associated contamination. To resolve this ambiguity, the high-frequency recordings were decomposed into line-harmonic-associated and residual broadband components (Fig. 5d,e). In comparison to CNT-3D, CNT-2D demonstrated 8.9-fold elevated total high-frequency power, encompassing a 35.9-fold escalation in line-harmonic-associated power and a 6.0-fold augmentation in the residual broadband component. This observation signifies that the discernible high-frequency enhancement observed in CNT-2D recordings could not be ascribed to a solitary narrowband interference source (Fig. 5d,e and Supplementary Fig. 26). The apparent enhancement in CNT-2D at 70-150 Hz exhibited a heightened sensitivity to the progressive exclusion of line-frequency harmonics, substantiating a substantial contribution from harmonic-associated contamination within this nominal high-gamma band (Supplementary Fig. 27).

Further analysis, including additional inter-channel correlation, coherence, common-mode, and principal-component analyses, demonstrated that the elevated CNT-2D high-frequency component exhibited stronger shared characteristics across spatially separated channels (Supplementary Figs. 28 and 29). This finding aligns with the hypothesis of a spatially diffuse rather than localized cortical origin. Temporal-envelope analysis yielded complementary evidence, with CNT-2D exhibiting a more persistent and less burst-like 70-150 Hz component than CNT-3D (Supplementary Fig. 30). Collectively, these analyses indicate that the observed 70-150 Hz enhancement in CNT-2D was consistent with line-harmonic-sensitive and spatially diffuse non-physiological contamination rather than localized cortical high-gamma activity.

As illustrated in Figure 5f, the channel-resolved maps of low-band spectral SNR were examined. CNT-3D demonstrated a higher recording-quality regime across the array than CNT-2D, indicating a spatially consistent shift toward higher-fidelity recording across the CNT-3D interface. This pattern was corroborated by complementary 4 × 4 maps of robust peak-to-peak amplitude, 0.5-50 Hz ECoG power, and 300-1000 Hz noise floor, which showed broadly higher signal amplitude and low-frequency power together with a lower high-frequency noise floor for CNT-3D across recording sites (Supplementary Fig. 31).

As for the chronic implantation, ECoG recordings demonstrated that the early recording-quality advantage of CNT-3D persisted throughout the 120-day implantation period (Fig. 5g and Supplementary Figs. 32 and 33). Conversely, the physiologically relevant low-frequency activity recorded by CNT-2D gradually diminished over time. The quantification of low-frequency power retention revealed a clear distinction between the two interfaces. At Day 120, CNT-3D retained approximately 80.4% of its Day 0 power, whereas CNT-2D retained only 11.7% (Fig. 5g and Supplementary Fig. 34). Consistent with this divergent temporal evolution, the CNT-3D/CNT-2D robust peak-to-peak amplitude ratio increased from approximately 2.97 at Day 0 to 7.79 at Day 120, while the corresponding low-band SNR difference increased from 15.08 to 23.53 dB (Fig. 5h and Supplementary Fig. 35). In contrast, the 300-1000 Hz noise floor exhibited a comparable increase in both arrays during the specified period (+3.79 dB for CNT-2D and +3.65 dB for CNT-3D; Supplementary Fig. 36). Therefore, the chronic recording advantage of CNT-3D arose primarily from preservation of physiologically relevant low-frequency signal, supporting sustained recording fidelity of the protruding 3D interface during chronic implantation.

Then, we tested the functional channel yield of CNT-2D and CNT-3D arrays during 120-day implantation, using a SNR threshold of 35 dB (Figure 5i). CNT-3D maintained a higher proportion of high-fidelity recording channels over this experiment and preserved substantially greater functional yield at Day 120 than CNT-2D. Figure 5j and Supplementary Figure 37 jointly examined the recording quality and the 1 kHz in vivo interface impedance established in Figure 4. This analysis yielded a discernible operating regime for CNT-3D, which is distinguished by reduced impedance, augmented low-frequency ECoG power, and diminished residual high-frequency noise.

### Negligible artifacts in postoperative MRI

Beyond sustained ECoG recording, compatibility with postoperative MRI is valuable for visualizing surrounding anatomy and supporting longitudinal assessment after implantation^35,36^. Therefore, we evaluated the MRI performance of the CNT-3D array and a size-matched Pt-Ir array using the bilateral implantation protocol (Fig. 6a,c and Supplementary Figs. 38 and 39). A posterior titanium cranial screw was retained as an additional metallic reference, and all implanted components were examined under the identical MRI acquisition conditions (Fig. 6c and Supplementary Fig. 40).

**Fig. 6.**
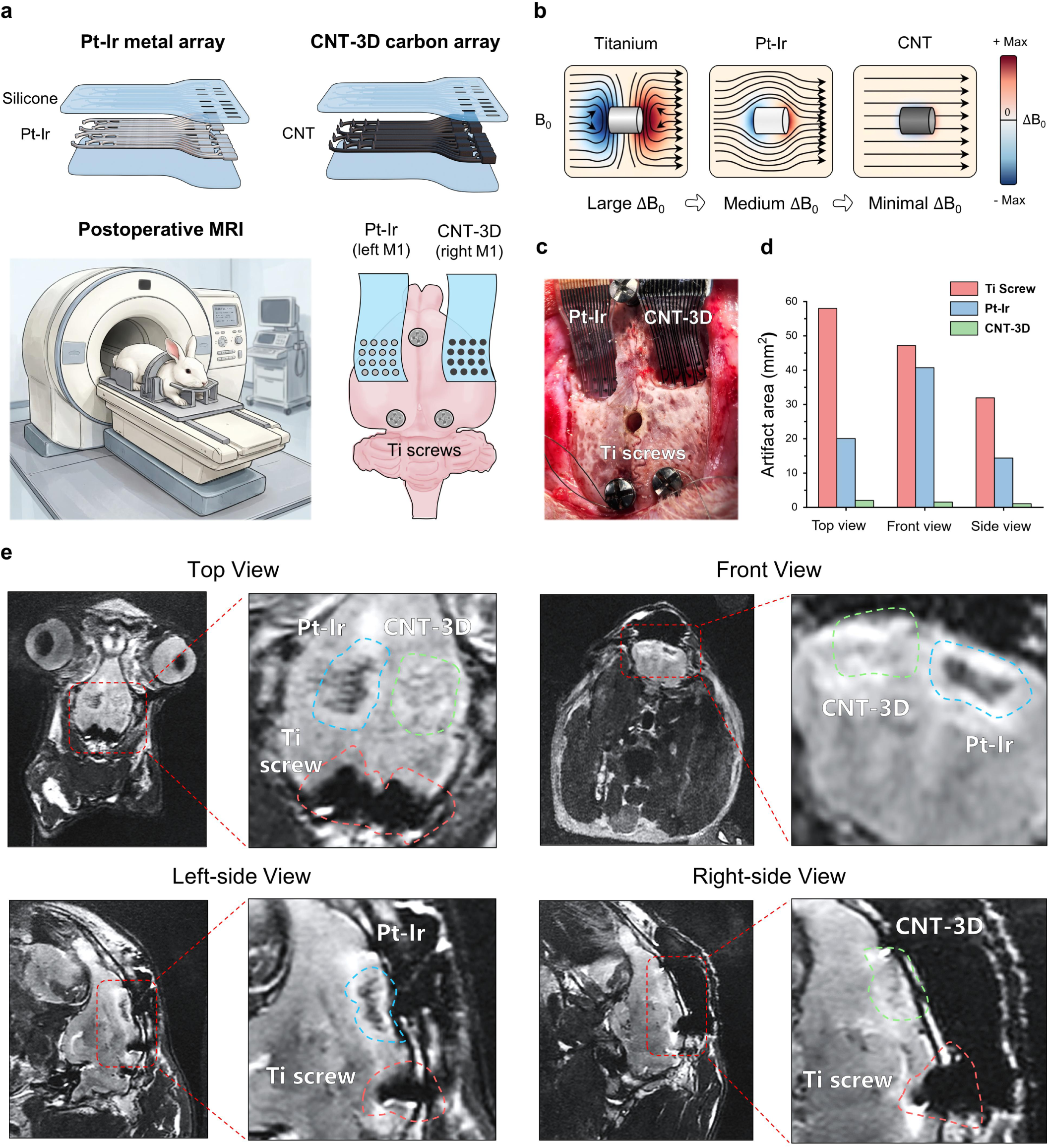
Postoperative MRI results of Pt-Ir array, CNT-3D array and Ti screws. **a,** Experimental design for postoperative 5.0 T MRI evaluation in a New Zealand white rabbit implanted with bilateral size-matched Pt-Ir and CNT-3D arrays, together with a posterior titanium cranial screw. **b,** Schematic illustrating MRI artifact formation arising from magnetic susceptibility mismatch. **c,** Photograph of the implanted rabbit brain showing the locations of the Pt-Ir array, CNT-3D array, and titanium screw. **d,** Quantification of MRI artifact areas measured from the top, front, left- and right-side views. **e,** Representative MRI images acquired from the top, front, left- and right-side views.

The magnetic-susceptibility mismatch between the implants and surrounding tissue results in the perturbation of the local B_0_ field, leading to signal loss and spatial distortion in MRI images (Fig. 6b). The three implanted materials exhibited a clear artifact hierarchy: the titanium screw generated the strongest local distortion, Pt-Ir produced an intermediate but clearly visible artifact, whereas CNT-3D caused only minimal image disturbance. Postoperative MRI acquired in top, front, left- and right-side views, showed that the cortical anatomy adjacent to CNT-3D remained clearly interpretable with minimal visible local distortion. Conversely, Pt-Ir produced localized image disruption, and the titanium screw generated substantially more extensive distortion (Fig. 6e). Quantitative analysis yielded mean artifact areas of 1.53 mm^2^, 25.05 mm^2^ and 45.71 mm^2^ for CNT-3D, Pt-Ir, and titanium, respectively, corresponding to a 16.4-fold reduction relative to Pt-Ir and a 29.88-fold reduction relative to titanium (Fig. 6d).

The MRI of an unimplanted control rabbit provided the anatomical baseline, while bilateral Pt-Ir controls reproduced visible artifacts at both cortical implant regions, confirming the reproducibility of the metal-associated image distortion (Supplementary Figs. 41-43). Collectively, these results demonstrate that the CNT-3D architecture establishes intimate and durable electrophysiological interfacing, while the metal-free CNTs network preserves high-field postoperative neuroimaging access.

## Discussion

The neural interfaces has been primarily driven by the pursuit of device-scale conformality, with the objective of reducing mechanical mismatch with the brain tissue. However, device conformality does not guarantee intimate interface at individual sensing sites. This work shift soft cortical interfacing from device-scale softness toward site-scale coupling. In this instance, the 3D soft carbon sites extend the conductive interface beyond the confines of the substrate plane without penetrating the cortex. The utilization of 3D sensing sites simultaneously engineers the physical electrode-cortex geometry and the electrochemical interface, which has been demonstrated to result in consistent superiority over the planar counterpart. More importantly, the protruding sites relocate the active surface toward the cortex, yielding enhanced cortical signal acquisition and diminished interface-associated interference.

Furthermore, the 3D soft carbon array integrates interconnects, electrode sites, and pads within a shared CNTs@silicone conductive phase, which is embedded within the soft silicone structure. This approach eliminates the requirement for serpentine metal traces and unstable electrode-coating interface, while retaining the mechanical and electrochemical advantages associated with carbon-based neural electrodes. Enabled by this metal-free architecture, the 3D soft carbon array generates negligible implant-induced MRI artifacts, providing a promising platform for artifact-minimized postoperative neuroimaging in chronically implanted neural interfaces.

The laser drawing fabrication strategy is also of significance because it is challenging to realize 3D silicone interfaces with conventional manufacturing routes. The depth-tunable laser process utilizes a distinct approach, whereby the vertical microstructure is programmed directly through laser penetration depth. Consequently, modifications can be made to electrode height, diameter, routing and array geometry without changing the fundamental materials. The key advance is not just the simplification of processes, but the digitally programmable dimension and versatile fabricated neural electrodes. This provides a direct route from planar flexible metal electronics to customizable 3D soft carbon interfaces.

Ultimately, the fidelity of neural signals from medium to long-term implants not only depends on the electrode materials, but is also influenced by the structure of the sensing site. Broader translation of 3D soft cortical interfaces will require validation in larger animal models and ultimately, clinical evaluation. Concurrently, scaling toward higher-density microelectrode arrays will demand finer spatial resolution, tighter depth control, and greater fabrication uniformity from the laser drawing strategy. Overall, this work establishes a laser drawing strategy for versatile soft 3D carbon neural interfaces, laying the foundational work toward high-performance and clinically deployable platforms for next-generation brain-computer interfaces.

## Methods

### Design of the 3D soft carbon neural interface

The 3D soft carbon electrode arrays were designed with a silicone/CNTs@silicone/silicone sandwich structure. The 16-channel cortical array comprised a 4 × 4 arrangement over an active area of 5 × 5 mm², with an electrode site diameter of 200 μm and an interelectrode pitch of 1.2 mm. Each protruding CNT-3D microelectrode site possesses a height of 180 μm and a basal diameter of 200 μm. The interconnects had a width of 150 μm and were terminated in 1.75 × 0.3 mm² contact pads for electrical interfacing with the acquisition/stimulation system through the connection method described in Supplementary Fig. 12.

Three electrodes (Au-2D, CNT-2D, and CNT-3D) were designed for comparative characterization. CNT-2D arrays used the same CNTs@silicone conductive composite, silicone architecture, channel configuration and site diameter as the CNT-3D arrays, but incorporated planar electrode sites. Au-2D control arrays were fabricated using conventional MEMS processes, consisting of a PI/Cr/Au/PI thin-film stack with planar Au electrode sites measuring 200 μm in diameter. The Au-2D array functioned as conventional metallic thin-film controls, whereas the comparison between the composition-matched CNT-2D and CNT-3D arrays was used to distinguish the contribution of 3D site microstructure.

The 3D soft carbon neural interface was further designed in multiple channel counts and form factors to accommodate different anatomical targets and spatial scales. The channel number, site dimensions, interelectrode spacing, substrate outline and overall array footprint were adapted to the corresponding target anatomy, while retaining the same CNTs@silicone conductive architecture and laser-defined fabrication principle. Device specific dimensions are provided in Supplementary Table 4.

### Preparation of the CNTs-embedded silicone composite

The conductive composite was consisted of a silicone elastomer matrix (SYLGARD™ 184, Dow Inc., USA) and two types of multi-walled carbon nanotubes (MWCNTs) with nominal lengths of approximately 10-30 μm and 0.5-2 μm (XFM13 and XFM70, respectively; XFNANO, Jiangsu, China). The silicone precursor was initially prepared by combining the base and curing agent at a mass ratio of 10:1. This mixture was then subjected to vacuum-degassing for a duration of 5 min. The 10-30 μm and 0.5-2 μm MWCNTs were premixed at a mass ratio of 10:3, and subsequently incorporated into the silicone precursor at a silicone-to-MWCNT mass ratio of 5:1. The mixture was subsequently subjected to a homogenization process under vacuum conditions in a planetary mixer (BHZ-150, BHLAB, Henan, China) at a speed of 500 rpm for 120 s, followed by an additional 200 s of operation at 1,500 rpm. The resultant CNTs@silicone composite was then utilized in the fabrication of interconnects, 3D sites and pads.

### Laser drawing fabrication workflow

The core fabrication strategy of 3D soft carbon neural interfaces was accomplished through an depth-tunable laser drawing method, which is based on the water-soluble PVA and acetone-soluble PMMA sacrificial templates. The silicone material (NuSil MED-4244, Avantor^®^, USA) was prepared by mixing the base and curing agent at a mass ratio of 10:1, followed by planetary mixing and vacuum degassing. Then, it was spin-coated onto a 4 inch, 10 mm thick PMMA substrate at 2,000 rpm for 180 s, vacuum-treated to remove trapped air and thermally cured at 90 °C for 2 h. After that, the cured silicone thin film was treated with O₂ plasma (PT-5S, Sanhe Boda Electromechanical Technology, China) at 50% power and 40% oxygen for 120 s. A 76 μm thick commercial PVA film (L0515, Bowei POLYVA, China) was then laminated onto the surface of silicone film and hot pressed at 80 °C for 1 h to establish uniform contact and eliminate interfacial air pockets and wrinkles. The resulting PVA/silicone/PMMA stack functioned as the laser processing platform.

The depth-tunable laser drawing was performed using a 355 nm nanosecond UV laser (Polar2-355-5AZ1, Wuhan Huaray Precision Laser, China). The CAD-defined patterns corresponding to the interconnects, sites and pads were sequentially processed using region-specific laser parameters, including laser power/pulse energy, pulse duration, repetition rate, scanning speed and cycles, as summarized in Supplementary Table 2. For the interconnect regions, shallow laser ablation selectively penetrated the PVA film while retaining the underlying silicone layer. For the site regions, deeper ablation extended through the PVA and silicone layers and partially into the PMMA substrate, generating conical micro cavities that defined the final 3D carbon micro sensing sites. The contact pad regions were similarly penetrated through the PVA and silicone layers but to a shallower depth within the PMMA, forming recessed cavities for thickened 3D carbon pads. The CNTs@silicone composite was then manually spread across the laser-drawn surface using a scraper to fill the interconnect channels, electrode site and contact pad cavities. Excess composite material adhering to the surface of the PVA is then removed using anhydrous ethanol. The patterned CNTs@silicone layer was thermally cured at 90 °C for 1 h (LC-DZK-6055B, Lichen Scientific Instrument, China). Subsequently, the stack was immersed in 80 °C deionized water for 1 h to dissolve the PVA sacrificial mask.

After rinsing with deionized water and drying by oven, a second silicone layer (NuSil MED-4244, Avantor^®^, USA) was spin-coated at 2,000 rpm for 180 s and cured at 90 °C for 2 h. In this step, the 3D sites and pads remained embedded within the underlying PMMA cavities, whereas the interconnects became embedded between the two silicone layers. Then, the device outline was defined by UV laser cutting using the parameters listed in Supplementary Table 2. Finally, the entire stack was immersed in acetone for 10 min to dissolve the PMMA sacrificial mold and release the formed electrode array. The released arrays were rinsed with anhydrous ethanol and stored under constant temperature and humidity conditions for subsequent pad-to-pad connections.

### Morphological characterization

The surface morphology features of the 3D soft carbon neural interfaces were characterized by scanning electron microscopy (SEM; MIRA, TESCAN, Czech Republic). Before the imaging process, the samples were sputter-coated with a thin Au layer using an ion sputter coater (MC1000, Hitachi High-Tech, Japan). SEM imaging was performed using a secondary electron detector at an accelerating voltage of 10 kV and a beam current of 100 pA, with a typical working distance of 12 mm.

For cross-sectional characterization, representative electrode arrays were cryogenically cooled and sectioned with a sharp blade to expose the multilayer device structure. For the cross-sectional image presented in Fig. 1e, false coloration was implemented in ImageJ (National Institutes of Health, USA) to visually distinguish the silicone and CNTs@silicone regions. For compositional analysis, energy-dispersive X-ray spectroscopy (EDS; Bruker Nano GmbH, Berlin, Germany) integrated with the SEM was used to acquire elemental maps of CNT (C), oxygen (O), and silicon (Si) from representative device cross sections.

### In vitro assessment of cortical conformality

The cortical conformality of the 3D soft carbon neural interfaces was assessed by freshly harvested porcine brain tissue. The fresh porcine brains were obtained from a local slaughterhouse and used shortly after collection. The dura mater was carefully removed to expose the cortical and pial surfaces, which were maintained hydration with 0.1 M PBS (Biosharp, China) during the testing process. A 64-channel 3D soft carbon array was placed over the freshly harvested porcine cortex to assess device-scale adaptation to gyral-sulcal topography. Prior to this step, the PBS was applied to facilitate wet interfacial contact. Additionally, a 16-channel 3D soft carbon array were positioned over the swine porcine cortical regions containing pronounced gyri and sulci to assess site-scale coupling interface over fixed, highly curved cortical surface features.. For comparison, a commercial planner Pt-Ir electrode array “AirRay grids” (REF 1031.2094.90, CorTec GmbH, Germany) was placed on the same cortical regions.

### Electrochemical characterization

Electrochemical measurements were performed using the electrochemical workstation (CHI660E, CH Instruments, China). The conventional three-electrode configuration was employed, with the electrode array under test serving as the working electrode, an Ag/AgCl electrode functioning as the reference electrode and a Pt electrode acting as the counter electrode. The individual channels of the Au-2D, CNT-2D and CNT-3D arrays were characterized independently under identical conditions. Cyclic voltammetry (CV) was performed between −0.6 and 0.8 V versus Ag/AgCl, with a scan rate of 0.1 V s⁻¹ for three consecutive cycles. The second cycle was used for quantitative analysis. The calculation of the CSC_c_ was performed by integrating the cathodic current of the CV curve and normalizing by the potential scan rate and the projected geometric area of the electrode site,

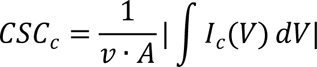

where () is the cathodic current, is the potential scan rate and is the projected geometric area of the electrode site. For all three electrodes, was defined by the nominal circular footprint of the 200 μm diameter electrode site, including the CNT-3D micro electrodes.

Electrochemical impedance spectroscopy (EIS) was performed over a frequency range of 1 Hz to 100 kHz at the open-circuit potential. The equivalent circuit fitting was performed in ZView (v.40h, Scribner Associates, USA).

### Voltage-transient stimulation characterization

Voltage-transient measurements were performed in 0.1 M PBS. Charge-balanced, cathodic-first biphasic current pulses were delivered between the working and counter electrodes using the Intan acquisition/stimulation system (RHS 2000, Intan Technologies, USA). Each pulse comprised a 150 μs cathodic phase at −500 μA (0.30-0.45 ms), a 50 μs interphase interval (0.45-0.50 ms) and a 150 μs anodic phase at +500 μA (0.50-0.65 ms), with zero current before and after stimulation. The working-electrode potential relative to the Ag/AgCl reference was recorded using a oscilloscope (DS2202A, RIGOL Technologies, China). Three channels were tested for each electrode type, with multiple pulses recorded from each channel under identical stimulation conditions.

Voltage traces were baseline-corrected using the pre-stimulation interval from 0.20-0.30 ms (Fig.3 i). The maximum cathodic polarization potential (*E_mc_*) was defined as the minimum baseline-corrected potential within the cathodic-phase analysis window (0.301-0.460 ms). Peak-to-peak voltage (*V_pp_*) was calculated as the difference between the maximum and minimum potentials within the stimulation window from 0.301 to 0.660 ms. The residual voltage was defined as the mean baseline-corrected potential over the final 20 sampled points of the recording window (0.781-0.800 ms). The recovery time was calculated from the end of the biphasic stimulus to the first post-stimulation time point at which the absolute voltage deviation from the baseline fell below _ℎ_ = max (3*_aeie_*, 0.05), where *σ_baseline_* is the standard deviation of the pre-stimulation baseline.

### Electrochemical stability under deformation and accelerated aging

Deformation stability of the CNT-3D arrays was evaluated using an mechanical testing system (IBTC-100SL, CARE Measurement and Control Test, China). Devices were subjected to 1,000 cycles of 180° bending, 180° twisting and stretching between 0 and 50% strain. Cyclic stretch loading was performed at 0.5 mm/s. Electrode impedance was measured in 0.1 M PBS using the impedance testing function of the Intan acquisition/stimulation system.

Accelerated aging test was performed by immersing CNT-3D arrays in 0.1 M PBS maintained at 73 °C based on the constant-temperature circulator (HMHX-08, Jiangsu Hengmin Instrument Manufacturing, China). The arrays were aged for 30 days, with impedance measurements performed on days 0, 6, 12, 18, 24 and 30. At each time point, the arrays were removed from the aging bath, equilibrated in 0.1 M PBS at room temperature and characterized using the Intan acquisition/stimulation system.

### Animal subjects, surgical procedures and electrode implantation

All animal procedures were approved by the Animal Ethics Committee of Hainan University (Approval Number: HNUAUCC-2025-00336). The experimental model comprised one male Sprague-Dawley rat (297 g) and four male New Zealand white rabbits (2.86, 2.94, 3.09 and 3.15 kg). General anesthesia was induced and maintained with inhaled isoflurane throughout the surgical procedure.

In the rat experiment, the animal was secured within a stereotaxic frame, and a craniotomy approximately 5 × 5 mm² was created over the cerebral hemisphere. The dura mater was removed, and 9-channel electrode arrays were gently placed on the cortical surface for acute signal recording. The animal was euthanized using deep anesthesia after the forepaw stimulation experiment.

For the rabbit experiments, bilateral cranial windows approximately 5 × 5 mm² were created over the primary motor cortex (M1). This was followed by the removal of the dura mater to expose the cortical surface. The electrode arrays were positioned over bilateral M1 regions and connected to the corresponding reference and ground skull electrodes and back-end interface. The connector assemblies were secured with dental cement before scalp closure. After the surgical procedure, erythromycin ointment was applied topically to the incision, followed by subcutaneous enrofloxacin (5.0 mg kg⁻¹) and meloxicam (0.3 mg kg⁻¹) as antibiotics and analgesics, respectively.

### In vivo impedance measurement

The in vivo impedance was measured using the Intan acquisition/stimulation system. The implanted 16-channel arrays were connected to the Intan system through the customized board. A titanium screw positioned adjacent to the electrode array served as the reference (REF), whereas another titanium screw positioned posteriorly and farther from the array served as the ground (GND).

At each measurement frequency, the Intan system applied a sinusoidal test signal and recorded the corresponding voltage response across the in vivo electrodes. Impedance magnitude and phase were obtained as 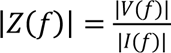 and θ_z_(f) = θ_v_(f) - θ_i_(f) respectively. The same measurement protocol used for chronic in vivo impedance tracking from day 0 to day 120.

### ECoG recordings

For the forepaw stimulation experiments on rat, stimulation needles were inserted into the left forepaw. The needle was connected to the Intan system for electrical currents delivery. Cathodic-first, charge-balanced biphasic square-current pulses with an amplitude of 5 mA were delivered at 50 Hz for 10 s stimulation trains, with a phase duration of 20ms and no interphase interval. A 9-channel electrode array was placed epidurally over the right hemisphere and connected to the neural data acquisition system (Apollo II, Bio-Signal technologies, China). Cortical signals were acquired at 30 kHz. Au-2D, CNT-2D and CNT-3D arrays were systematically evaluated in the same animal under consistent conditions.

For chronic bilateral ECoG recordings, 16-channel CNT-2D and CNT-3D arrays implanted over bilateral M1 regions of New Zealand white rabbit were simultaneously connected to the Intan system. The titanium reference and ground screws used for in vivo impedance measurements were retained for ECoG acquisition. Continuous ECoG signals were recorded at 30 kHz for 5 min under consistent awake, resting conditions on postoperative day 0, 30, 60, 90, and 120.

### ECoG analysis

ECoG signals were processed using customized scripts in MATLAB R2022b (The MathWorks, USA), and all script code is available as open access. The raw recordings were acquired at 30 kHz and processed using analysis-specific pipelines. The quantitative results were exported from MATLAB as numerical tables and subsequently plotted in Origin (2024, OriginLab, USA).

For chronic ECoG recordings, the CNT-2D and CNT-3D datasets were processed using identical analysis procedures. Low-frequency cortical activity was defined as 0.5-50 Hz, the 70-150 Hz range was designed as a high-frequency diagnostic band, and 300-1,000 Hz was employed to characterize broadband noise. Power spectral density (PSD) was estimated using Welch’s method with 2 s windows, 50% overlap and an Non-Uniform Fast Fourier Transform (NFFT) of 65,536. For the primary spectral analyses, ±1 Hz windows around 50-Hz power-line harmonics were excluded. The robust peak-to-peak amplitude was calculated as the difference between the 99th and 1st percentiles of the 0.5-50-Hz band-pass-filtered signal. The broadband noise floor was defined as the median PSD within 300-1,000 Hz after harmonic exclusion. The low-band spectral SNR was calculated as *10*·*log_10_(P_LF_/P_noise_)*, where *P_LF_* is the mean PSD within 0.5-50 Hz and *P_noise_* is the median PSD within 300-1,000 Hz. Channel-wise metrics were mapped onto the physical 4 × 4 electrode layout for spatial comparison.

### MRI compatibility assessment

The in vivo MRI compatibility was evaluated using the magnetic resonance imaging system (Jupiter 5.0 T, United Imaging Healthcare, China). Rabbits were anesthetized by intramuscular injection of Zoletil 50 (10 mg kg⁻¹) into the anterior thigh before MRI. For the primary comparison, a New Zealand white rabbit implanted with bilateral 16-channel CNT-3D and size-matched Pt-Ir arrays, together with a posterior titanium cranial screw, was imaged to evaluate implant-induced MRI artifacts. To ensure the reliability of the findings, additional scans were acquired from a non-implanted control rabbit and a rabbit implanted bilaterally with Pt-Ir arrays. The MRI images were reviewed using U_VIEWER software (United Imaging Healthcare, China). The artifacts were evaluated in top, front, left- and right-side views. Artifact regions were manually delineated and artifact areas were calculated in ImageJ (v1.54u8, National Institutes of Health, USA) according to regions showing evident signal loss or geometric distortion.

## Data Availability Statement

The main data supporting the results in this study are available within the paper and its Supplementary Information. Source data are provided with this paper. Any further data of this study are available from the corresponding author upon reasonable request.

## Acknowledgements

The authors gratefully acknowledge financial support from the National Natural Science Foundation of China (62674196), the Science and Technology Innovation 2030-Major Project (2022ZD0208601, 2022ZD0208600), the Innovation Capability Support Program of Shaanxi (No. 2024RS-CXTD-17).

## Author contributions

**Conceptualization and methodology:** Xuanqi Wang and Bowen Ji; **Experiments and visualization:** Xuanqi Wang; **Funding acquisition, project administration, and supervision:** Bowen Ji; **Writing-original draft:** Xuanqi Wang. **Writing-review and editing:** Honglong Chang, Feng Yue, and Bowen Ji.

## References

1. Lee, J. M. et al. The ultra-thin, minimally invasive surface electrode array NeuroWeb for probing neural activity. Nat. Commun. 14, 7088 (2023).

2. Lacour, S. P., Courtine, G. & Guck, J. Materials and technologies for soft implantable neuroprostheses. Nat. Rev. Mater. 1, 16063 (2016).

3. Tringides, C. M. et al. Viscoelastic surface electrode arrays to interface with viscoelastic tissues. Nat. Nanotechnol. 16, 1019–1029 (2021).

4. Driscoll, N. et al. MXene-infused bioelectronic interfaces for multiscale electrophysiology and stimulation. Sci. Transl. Med. 13, eabf8629 (2021).

5. Zhao, S. et al. Full activation pattern mapping by simultaneous deep brain stimulation and fMRI with graphene fiber electrodes. Nat. Commun. 11, 1788 (2020).

6. Shen, K., Chen, O., Edmunds, J. L., Piech, D. K. & Maharbiz, M. M. Translational opportunities and challenges of invasive electrodes for neural interfaces. Nat. Biomed. Eng. 7, 424–442 (2023).

7. Ji, B. et al. Brainmask: an ultrasoft and moist micro-electrocorticography electrode for accurate positioning and long-lasting recordings. Microsyst. Nanoeng. 9, 126 (2023).

8. Minev, I. R. et al. Electronic dura mater for long-term multimodal neural interfaces. Science 347, 159–163 (2015).

9. Yi, J. et al. Water-responsive supercontractile polymer films for bioelectronic interfaces. Nature 624, 295–302 (2023).

10. Trouillet, A. et al. High-resolution prosthetic hearing with a soft auditory brainstem implant in macaques. Nat. Biomed. Eng. 9, 1403–1417 (2025).

11. Kern, M., Aertsen, A., Schulze-Bonhage, A. & Ball, T. Heart cycle-related effects on event-related potentials, spectral power changes, and connectivity patterns in the human ECoG. Neuroimage 81, 178–190 (2013).

12. Yin, J., Wang, S., Tat, T. & Chen, J. Motion artefact management for soft bioelectronics. Nat. Rev. Bioeng. 2, 541–558 (2024).

13. Lee, D. H. et al. Microelectrothermoforming (μETF): one-step versatile 3D shaping of flexible microelectronics for enhanced neural interfaces. npj Flex. Electron. 9, 5 (2025).

14. Wang, M. et al. Conformal bumped electrode web for chronic ECoG recordings in swine. Microsyst. Nanoeng. 12, 95 (2026).

15. Bai, R. et al. Ultrasoft pit-free electrodes: conformal low-noise electrochemical interface for ECoG recording. Sens. Actuators B Chem. 471, 141010 (2027).

16. Ji, B. et al. 3D soft microbump electrodes for elastic interaction with brain tissue. npj Flex. Electron. 9, 96 (2025).

17. Greiner, N. et al. Recruitment of upper-limb motoneurons with epidural electrical stimulation of the cervical spinal cord. Nat. Commun. 12, 435 (2021).

18. Metzger, S. L. et al. A high-performance neuroprosthesis for speech decoding and avatar control. Nature 620, 1037–1046 (2023).

19. Wyse-Sookoo, K. et al. Stability of ECoG high gamma signals during speech and implications for a speech BCI system in an individual with ALS: a year-long longitudinal study. J. Neural Eng. 21, 046016 (2024).

20. Chung, W. G. et al. Liquid-metal-based three-dimensional microelectrode arrays integrated with implantable ultrathin retinal prosthesis for vision restoration. Nat. Nanotechnol. 19, 688–697 (2024).

21. Chung, W. G. et al. An implantable epiretinal device for near-infrared light perception. Nat. Electron. 9, 569–580 (2026).

22. Yang, A. I. et al. Localization of dense intracranial electrode arrays using magnetic resonance imaging. Neuroimage 63, 157–165 (2012).

23. Succop, B. S. Jr, et al. Day one postoperative MRI findings following electrode placement for deep brain stimulation: analysis of a large case series. Front. Neurol. 14, 1253241 (2023).

24. Cho, Y. U. et al. MRI-compatible, transparent PEDOT:PSS neural implants for the alleviation of neuropathic pain with motor cortex stimulation. Adv. Funct. Mater. 34, 2310908 (2024).

25. Zhao, Q. et al. Highly stretchable and customizable microneedle electrode arrays for intramuscular electromyography. Sci. Adv. 10, eadn7202 (2024).

26. Liu, Z. et al. A three-dimensionally architected electronic skin mimicking human mechanosensation. Science 384, 987–994 (2024).

27. Wang, X. et al. 3D cone-shape carbon electrode array for high-fidelity ECoG acquisition. In 2026 IEEE 39th International Conference on Micro Electro Mechanical Systems (MEMS) 219–222 (IEEE, 2026).

28. Brown, M. A. et al. Direct laser writing of 3D electrodes on flexible substrates. Nat. Commun. 14, 3610 (2023).

29. Nimbalkar, S. et al. Glassy carbon microelectrodes minimize induced voltages, mechanical vibrations, and artifacts in magnetic resonance imaging. Microsyst. Nanoeng. 5, 61 (2019).

30. Ren, X. et al. Uniaxial extending neural probes for bleeding-absent implantation. npj Flex. Electron. 8, 36 (2024).

31. Yang, M. et al. Injectable hydrogel bioelectrostimulator for wireless deep brain neuromodulation. Nat. Commun. 17, 4526 (2026).

32. Bakhshaee Babaroud, N., et al. Multilayer CVD graphene electrodes using a transfer-free process for the next generation of optically transparent and MRI-compatible neural interfaces. Microsyst. Nanoeng. 8, 107 (2022).

33. Tintelott, M., Schander, A. & Lang, W. Understanding electrical failure of polyimide-based flexible neural implants: the role of thin film adhesion. Polymers 14, 3702 (2022).

34. Kim, H. J., Koo, J. H., Lee, S., Hyeon, T. & Kim, D.-H. Materials design and integration strategies for soft bioelectronics in digital healthcare. Nat. Rev. Mater. 10, 654–673 (2025).

35. Fallegger, F. et al. MRI-compatible and conformal electrocorticography grids for translational research. Adv. Sci. 8, 2003761 (2021).

36. Wang, X. et al. Subdural neural interfaces for long-term electrical recording, optical microscopy and magnetic resonance imaging. Biomaterials 281, 121352 (2022).

