## Supplementary Materials for "MRI-compatible 3D Soft Carbon Neural Interface Enabled by Versatile Depth-tunable Laser Drawing"

#### Abstract

Soft neural interfaces are designed to minimize mechanical mismatch with the brain. However, planar and recessed electrode sites remain locally decoupled from the curved and pulsating cortical surface, degrading the fidelity of neural sensing. Here, we report a 3D soft carbon electrode array, enabled by a depth-tunable laser drawing strategy. This laser-driven workflow achieves versatile fabrication of customizable 3D soft carbon neural interfaces, eliminating the reliance on conventional thin-film microfabrication processes. The fully silicone-based electrode structure realize device-scale conformality, and the 3D soft carbon sites promote a non-penetrating, intimate neural-electrode interface. The CNT-3D array combined superior in-vitro electrochemical and in-vivo recording performance compared with Au-2D and CNT-2D controls. Furthermore, in bilateral rabbit cortex implantation, CNT-3D array retained 80.4% of electrocorticography power after 4 months, versus 11.7% for CNT-2D control. Moreover, the 3D sensing sites significantly suppressed line-frequency harmonics and background noise interference. Subsequently, during magnetic resonance imaging, the CNT-3D array exhibits negligible implant-induced artifacts compared to the conventional Pt-Ir metal array. Overall, this work establishes a laser drawing strategy that extends neural interfaces from macro device-scale conformality to micro site-scale coupling, providing a technological pathway for practical and translational soft neurotechnology.

### List of Contents

#### Supplementary Figures

Supplementary Fig. 1.. 4
Supplementary Fig. 2. 5
Supplementary Fig. 3.. 6
Supplementary Fig. 4. 7
Supplementary Fig. 5. 8
Supplementary Fig. 6. 9
Supplementary Fig. 7. 10
Supplementary Fig. 8. 11
Supplementary Fig. 9. 12
Supplementary Fig. 10. 13
Supplementary Fig. 11. 14
Supplementary Fig. 12. 15
Supplementary Fig. 13. 16
Supplementary Fig. 14. 17
Supplementary Fig. 15. 18
Supplementary Fig. 16. 19
Supplementary Fig. 17. 20
Supplementary Fig. 18. 21
Supplementary Fig. 19. 22
Supplementary Fig. 20. 23
Supplementary Fig. 21. 24
Supplementary Fig. 22. 25
Supplementary Fig. 23. 26
Supplementary Fig. 24. 27
Supplementary Fig. 25. 28
Supplementary Fig. 26. 29
Supplementary Fig. 27. 30
Supplementary Fig. 28. 31
Supplementary Fig. 29. 32
Supplementary Fig. 30. 33
Supplementary Fig. 31. 34
Supplementary Fig. 32. 35

Supplementary Fig. 33. 36
Supplementary Fig. 34. 37
Supplementary Fig. 35. 38
Supplementary Fig. 36. 39
Supplementary Fig. 37. 40
Supplementary Fig. 38. 41
Supplementary Fig. 39. 42
Supplementary Fig. 40. 43
Supplementary Fig. 41. 44
Supplementary Fig. 42. 45
Supplementary Fig. 43. 46

#### Supplementary Tables

Supplementary Table 1. 47
Supplementary Table 2. 48
Supplementary Table 3. 49
Supplementary Table 4. 50

#### References

### Supplementary Figures

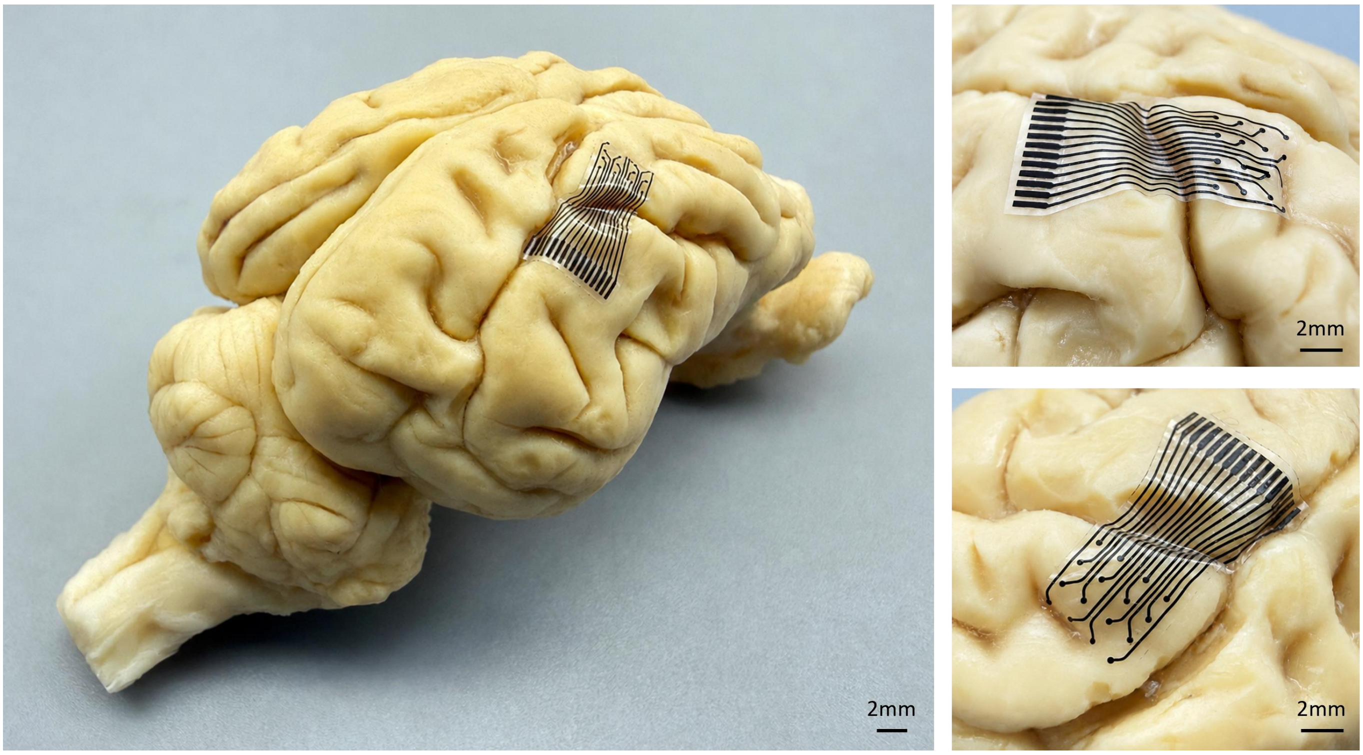


**Supplementary Fig. 1. Conformal interfacing of the 3D soft carbon electrode array with large curvature and complex gyral-sulcal topography of the porcine brain surface.**

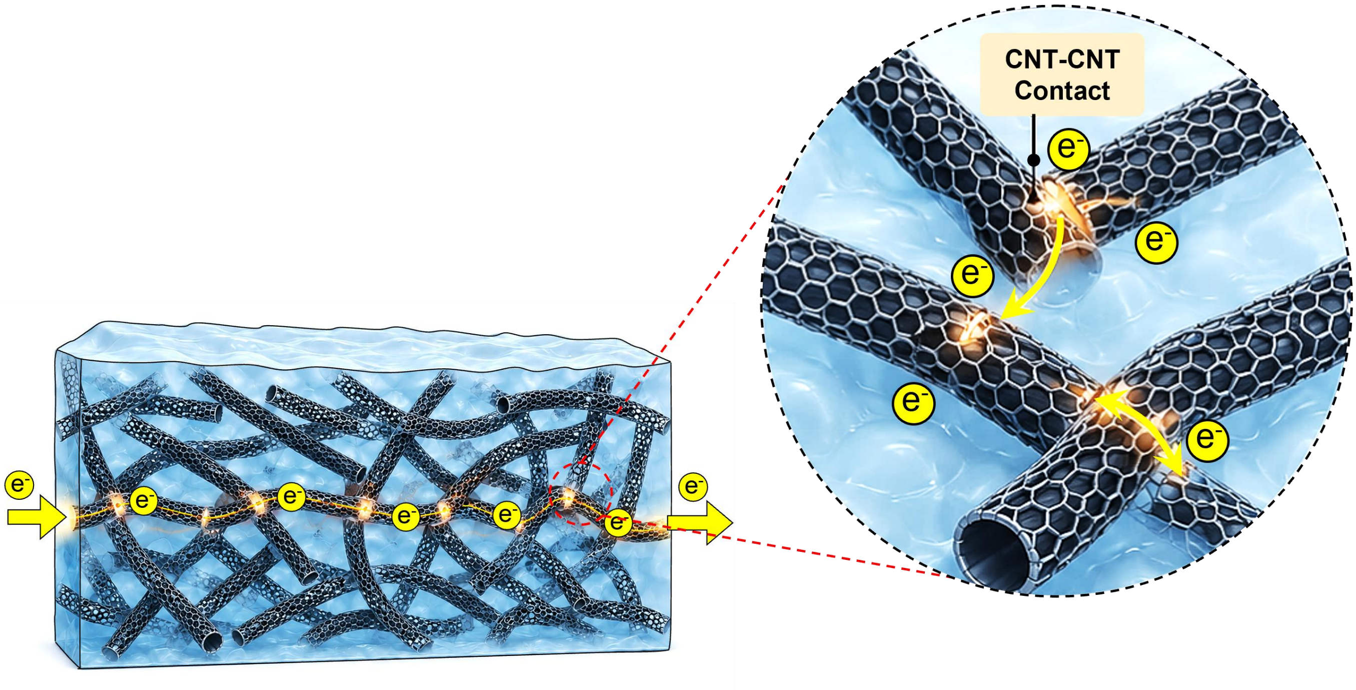


**Supplementary Fig. 2**. **Conductive mechanism of the CNTs percolation networks.** Dispersed carbon nanotubes form interconnected percolation pathways inside the silicone matrix, enabling charge transport via physical nanotube contacts and tunnelling junctions without metallic conductors.

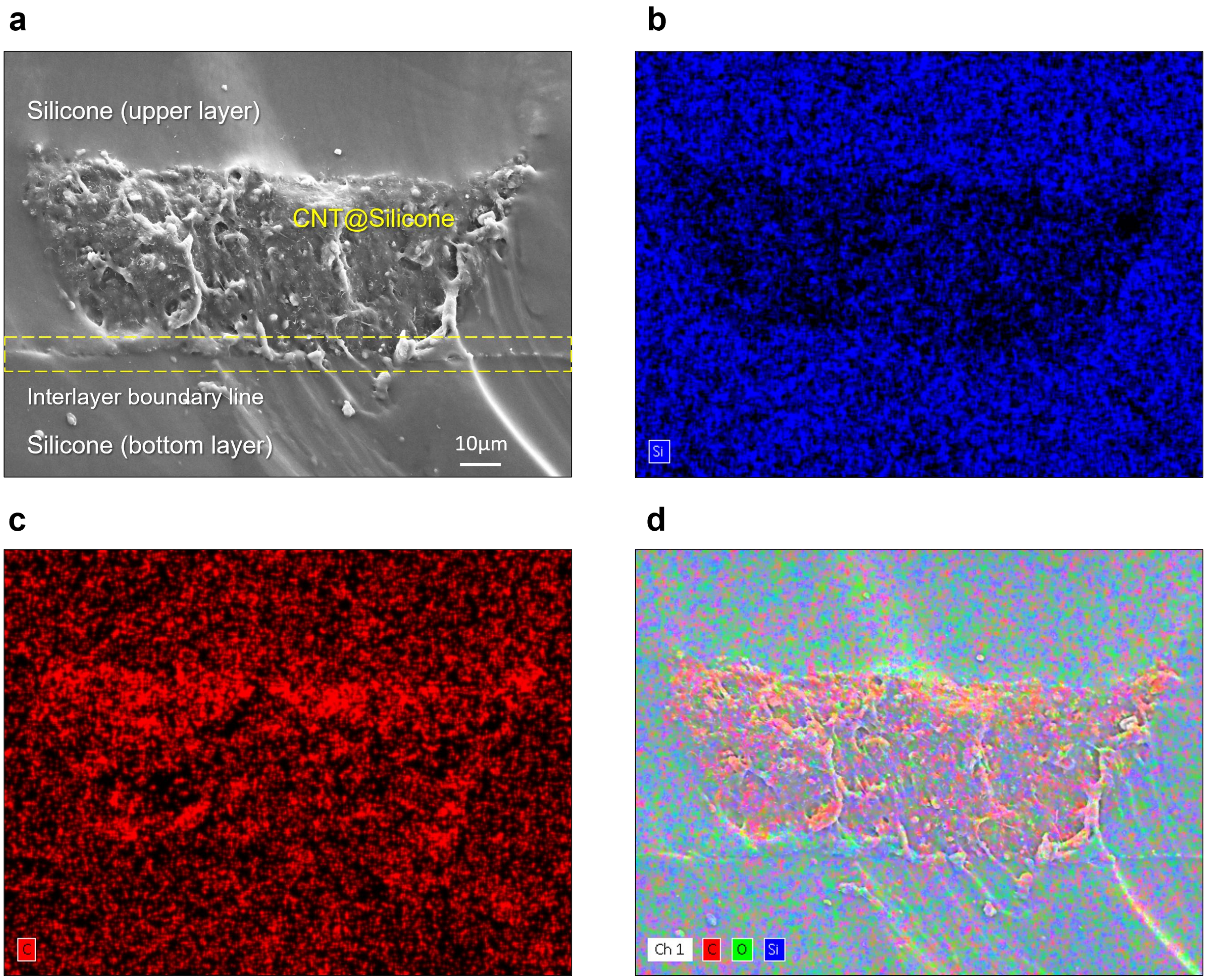


**Supplementary Fig. 3. Cross-sectional structure and elemental mapping. a,** Cross-sectional SEM image showing the upper silicone layer, the intermediate CNT@silicone interconnect, and the bottom silicone layer. Yellow parallel dashed lines indicate the interlayer boundaries. **b-d,** Corresponding EDS elemental maps of C, O and Si, respectively, showing the spatial elemental distributions across the multilayer cross-sections.

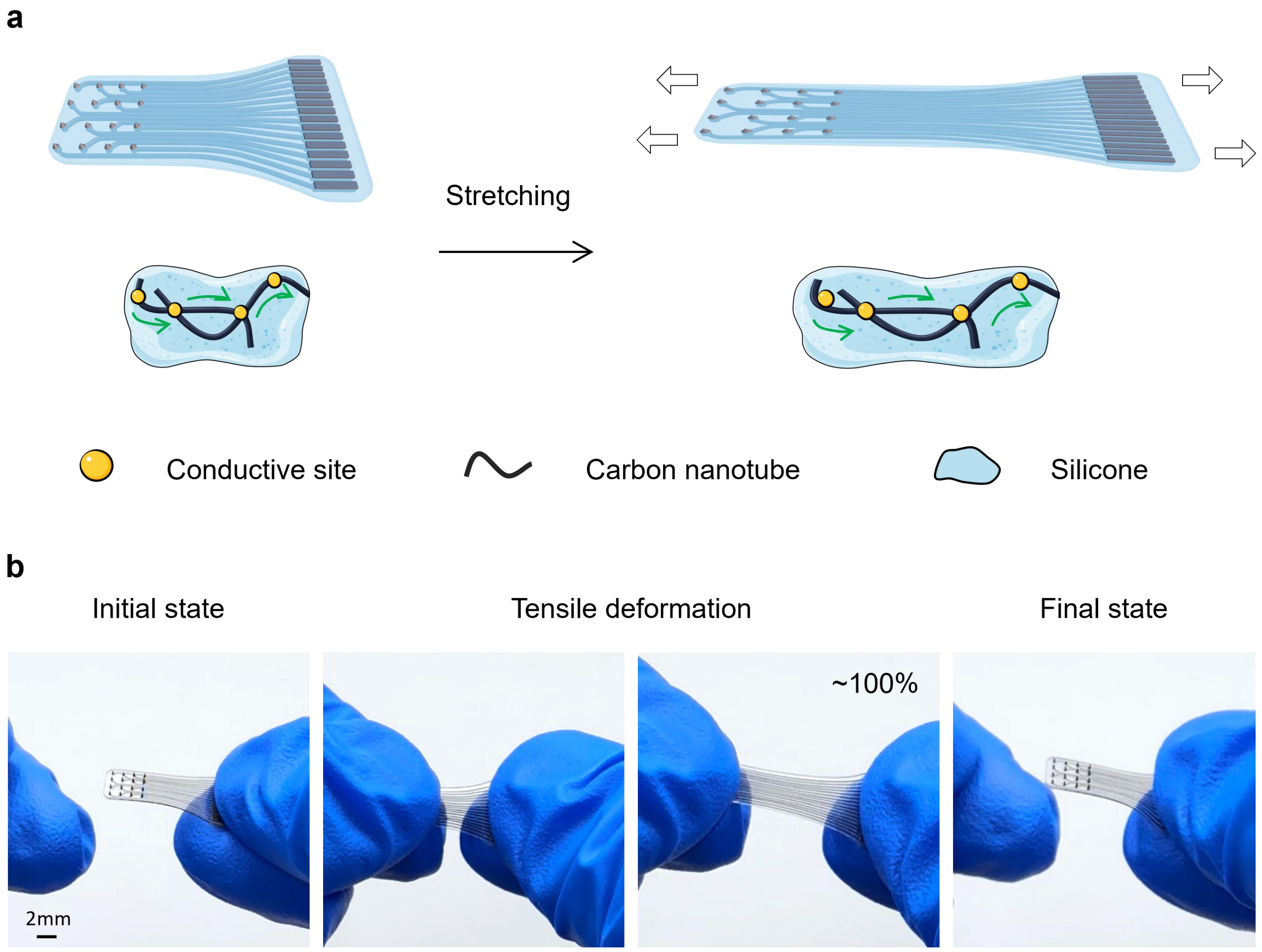


**Supplementary Fig. 4. Deformation stability of the 3D soft carbon electrode array. a,** Schematic illustration of the deformation-tolerant conductive network. The CNTs-embedded silicone conductive pathways preserve conductive percolation during curling, bending and stretching associated with surgical handling and post-implantation brain micromotion. **b,** Photographs of the array in the initial state, under tensile deformation and after release.

a b

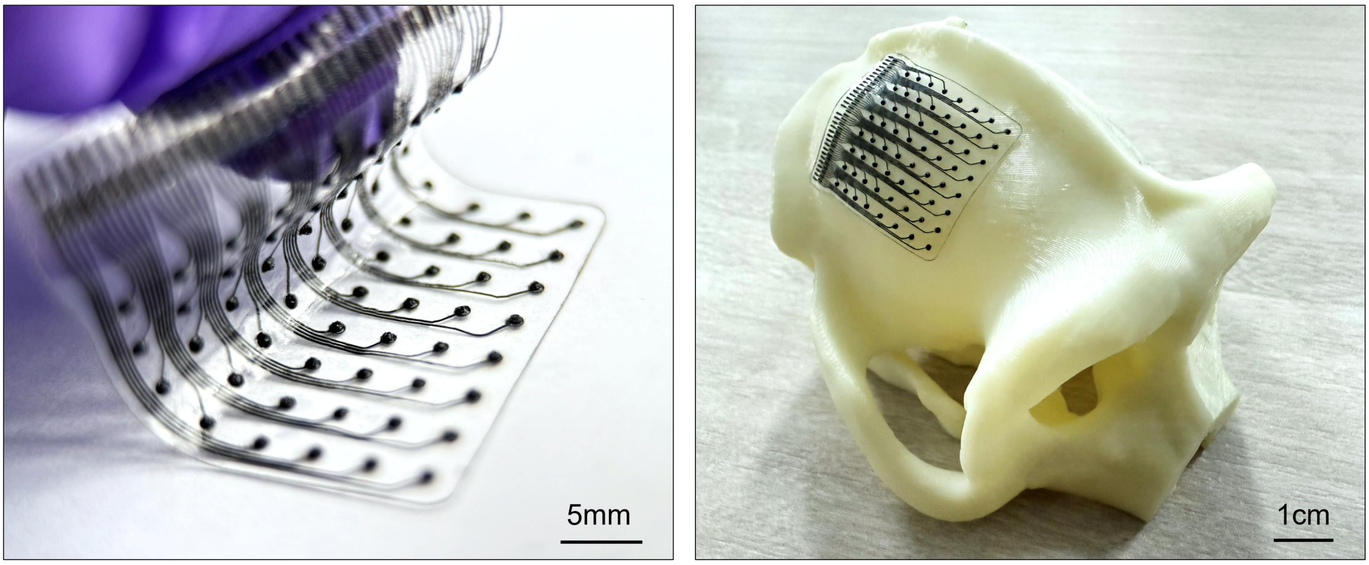


**Supplementary Fig. 5. Photograph of the 64-channel array with a primate cranial model. a,** Photograph of the fabricated 64-channel array. **b,** Conformal placement on a 3D-printed cynomolgus macaque skull model.

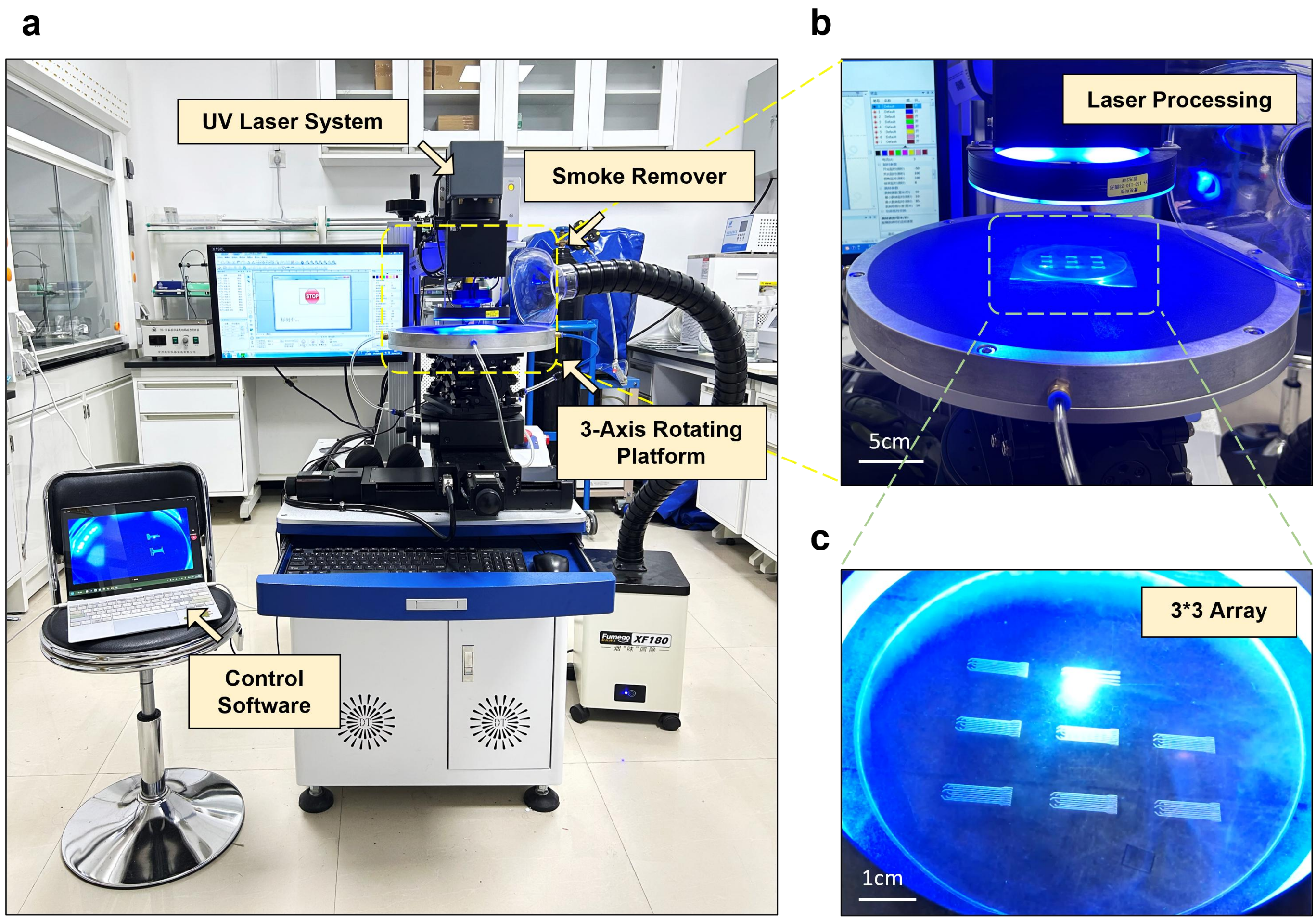


**Supplementary Fig. 6. UV Laser system for the fabrication of 3D soft carbon electrodes. a,** Photograph of the system, including the laser equipment, smoke extractor, three-axis rotary platform. **b,** Enlarged view of the three-axis rotary stage holding a 3-inch substrate during laser micromachining. **c,** Magnified view of the processed 3-inch substrate, nine arrays can be patterned within 10 min.

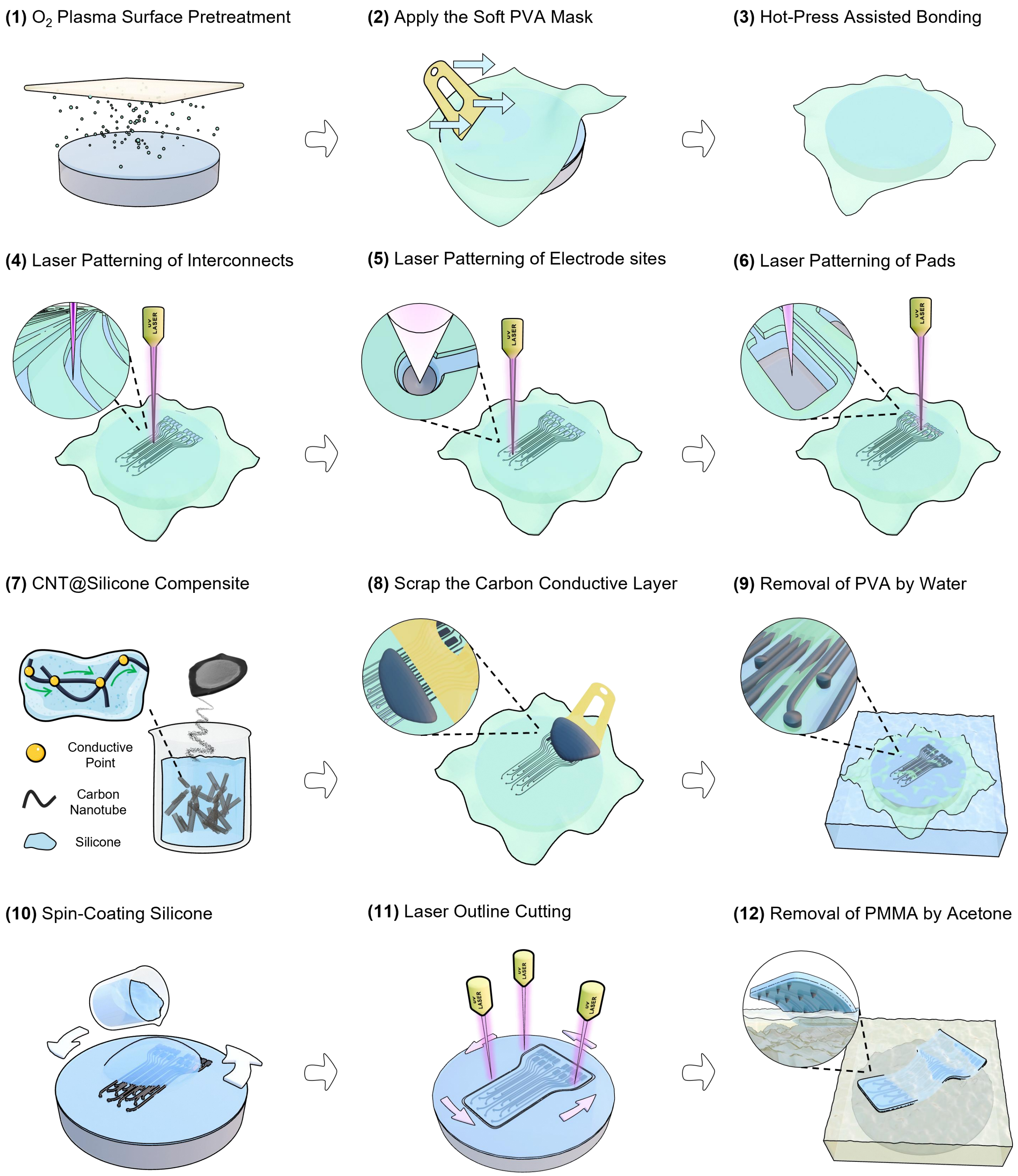


**Supplementary Fig. 7. Schematic illustration of the laser-driven fabrication.**

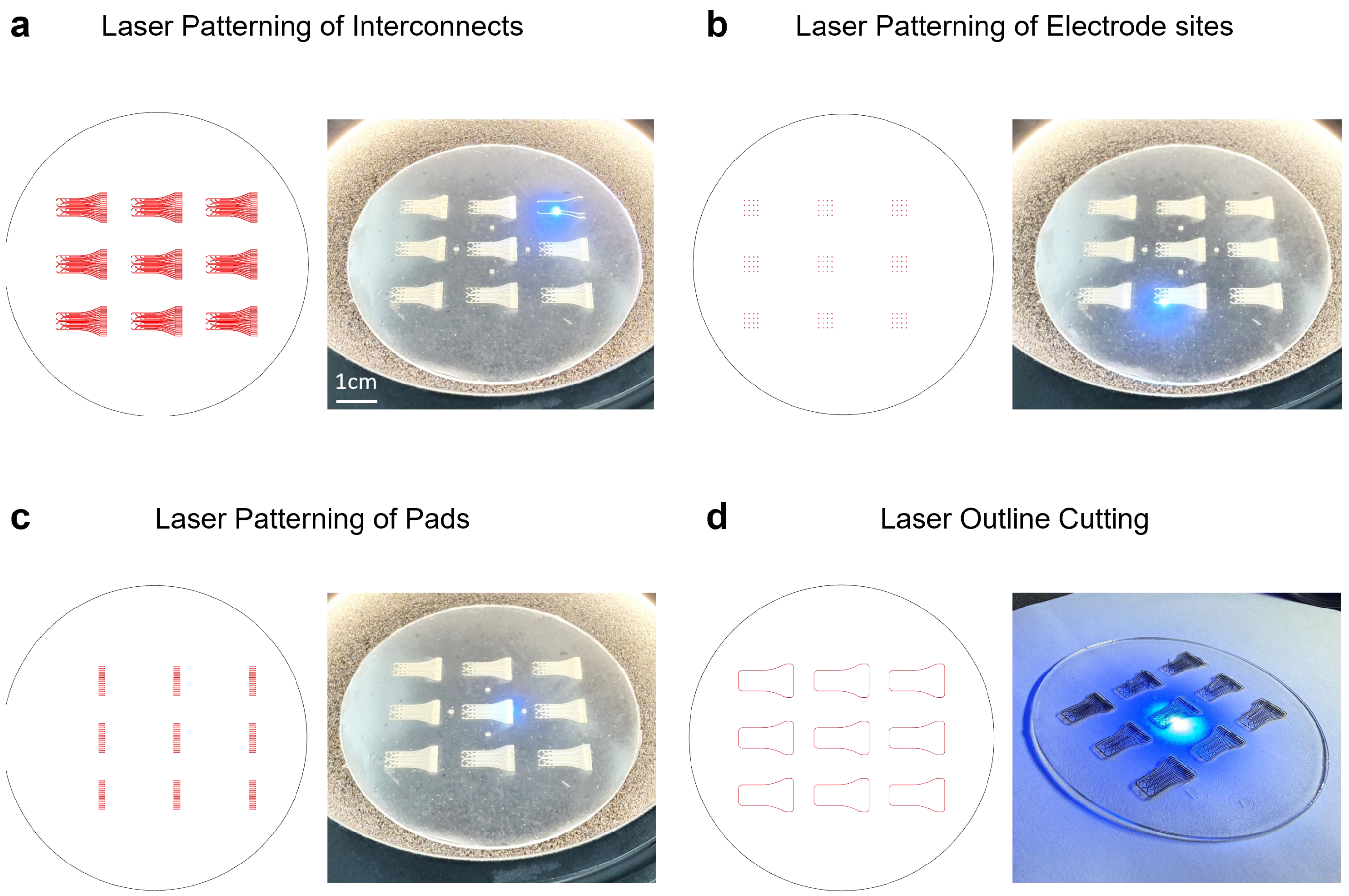


**Supplementary Fig. 8. Array-level visualization of laser darwing processes. a,** CAD layout and corresponding laser patterning photograph of the interconnects region pattern for 3 × 3 batch containing nine 16-channel arrays. **b,** the electrode sites. **c,** the contact pads pattern. **d,** the final outline cutting step.

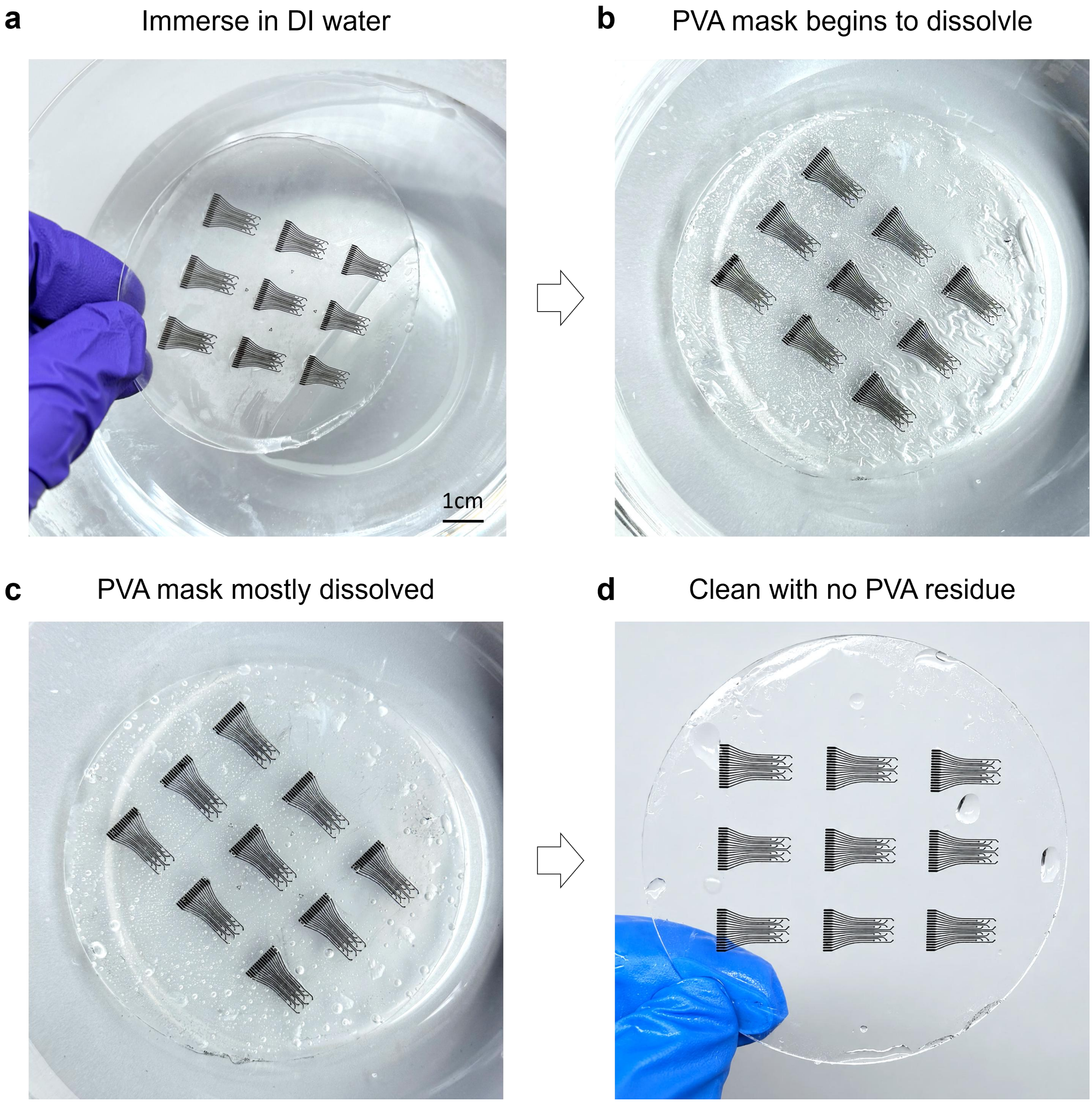


**Supplementary Fig. 9. Time-sequence photographs of the sacrificial PVA mask during dissolution in deionized (DI) water. a,** Initial immersion in DI water. **b,** Onset of PVA dissolution. **c,** The PVA mask mostly dissolved. **d,** Final clean structure after removal of the PVA mask.

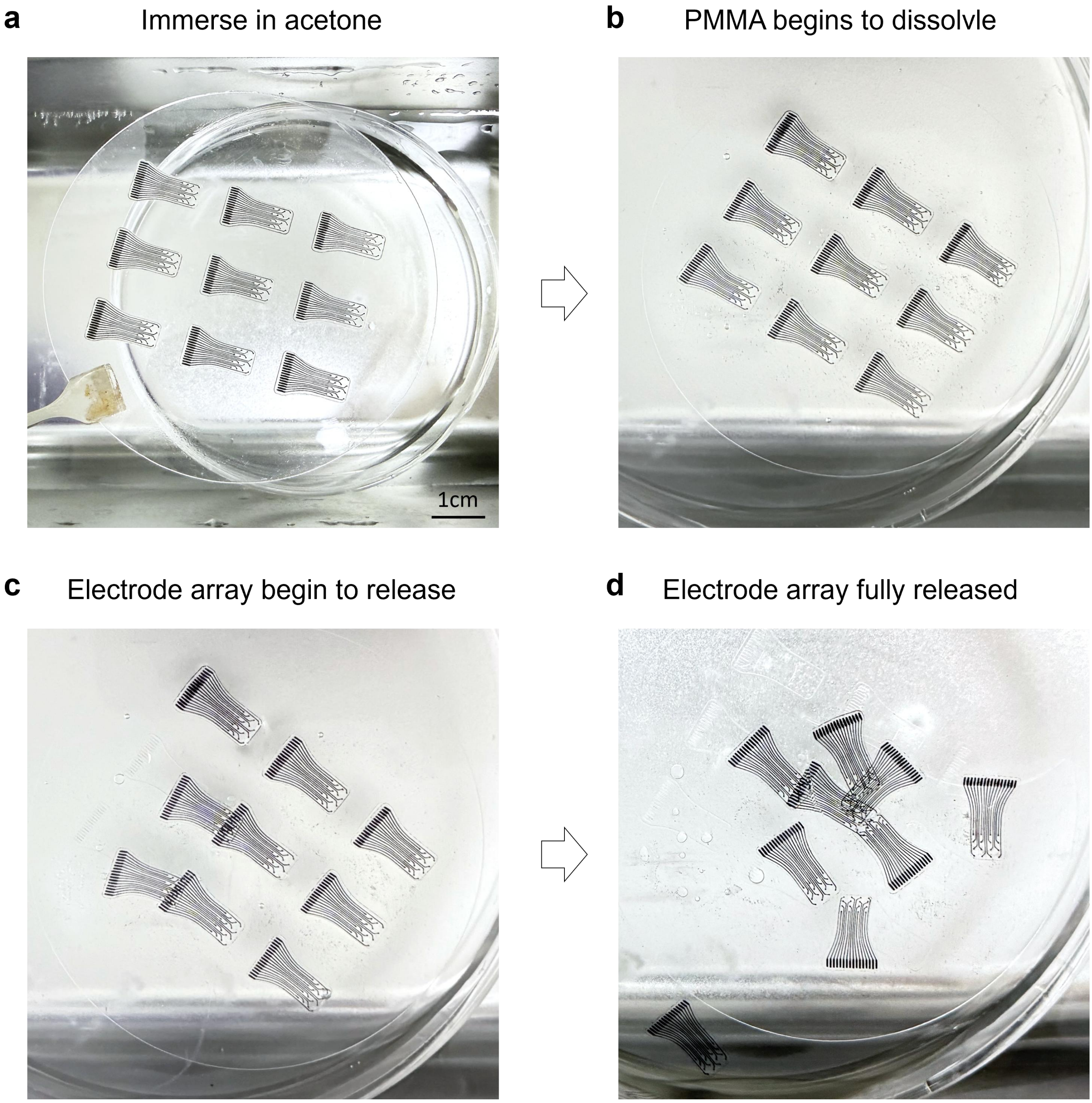


**Supplementary Fig. 10. Time-sequence photographs of the self-release process in acetone. a,** Initial immersion in acetone. **b,** Onset of PMMA dissolution. **c,** Automatic release of the electrode arrays without mechanical demolding. **d,** Fully released 3D soft electrode arrays.

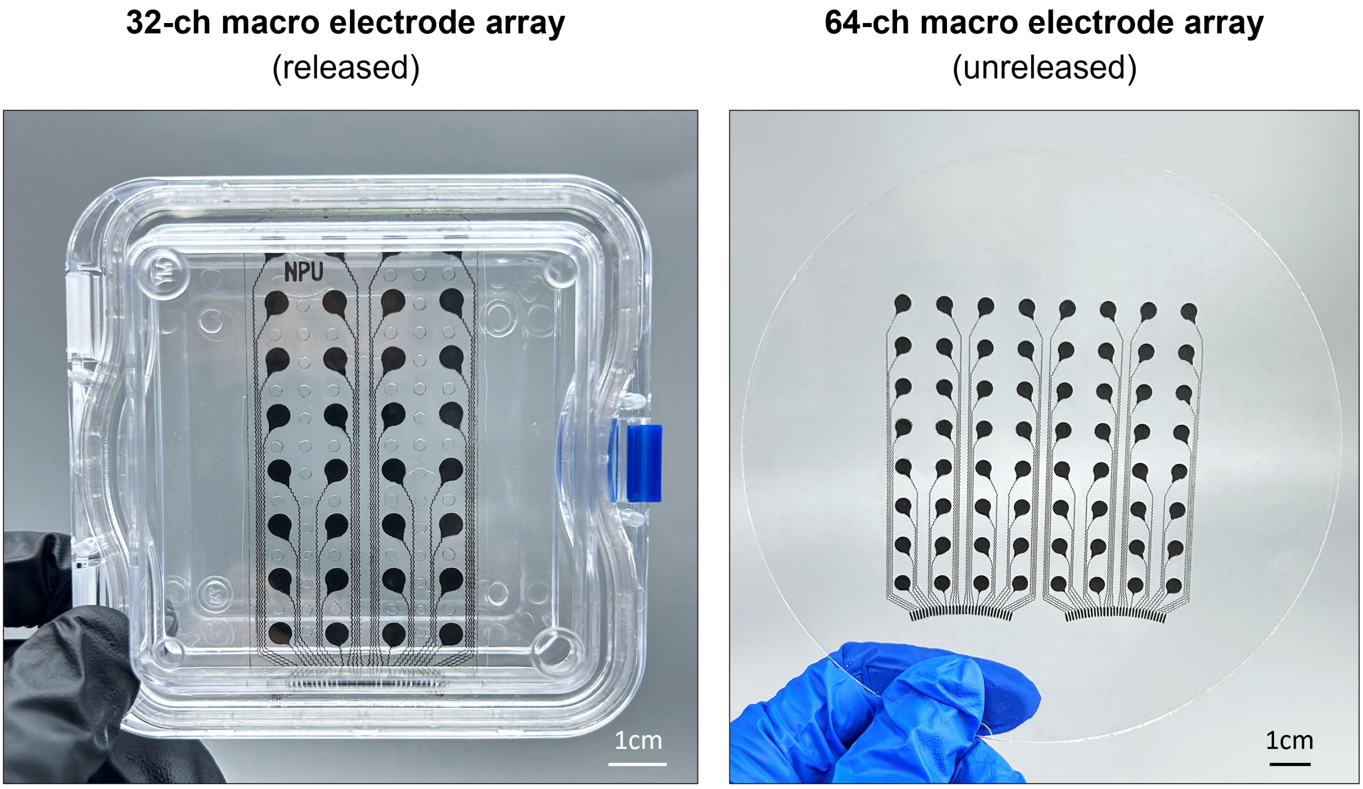


**Supplementary Fig. 11. Photograph of 32/64-channel macro array for human brain.** These array contain 2 mm diameter electrode sites and serpentine interconnects, demonstrating the scalability of the fabrication strategy from microscale electrode arrays to large-area macroelectrode interfaces with mechanically compliant wiring.

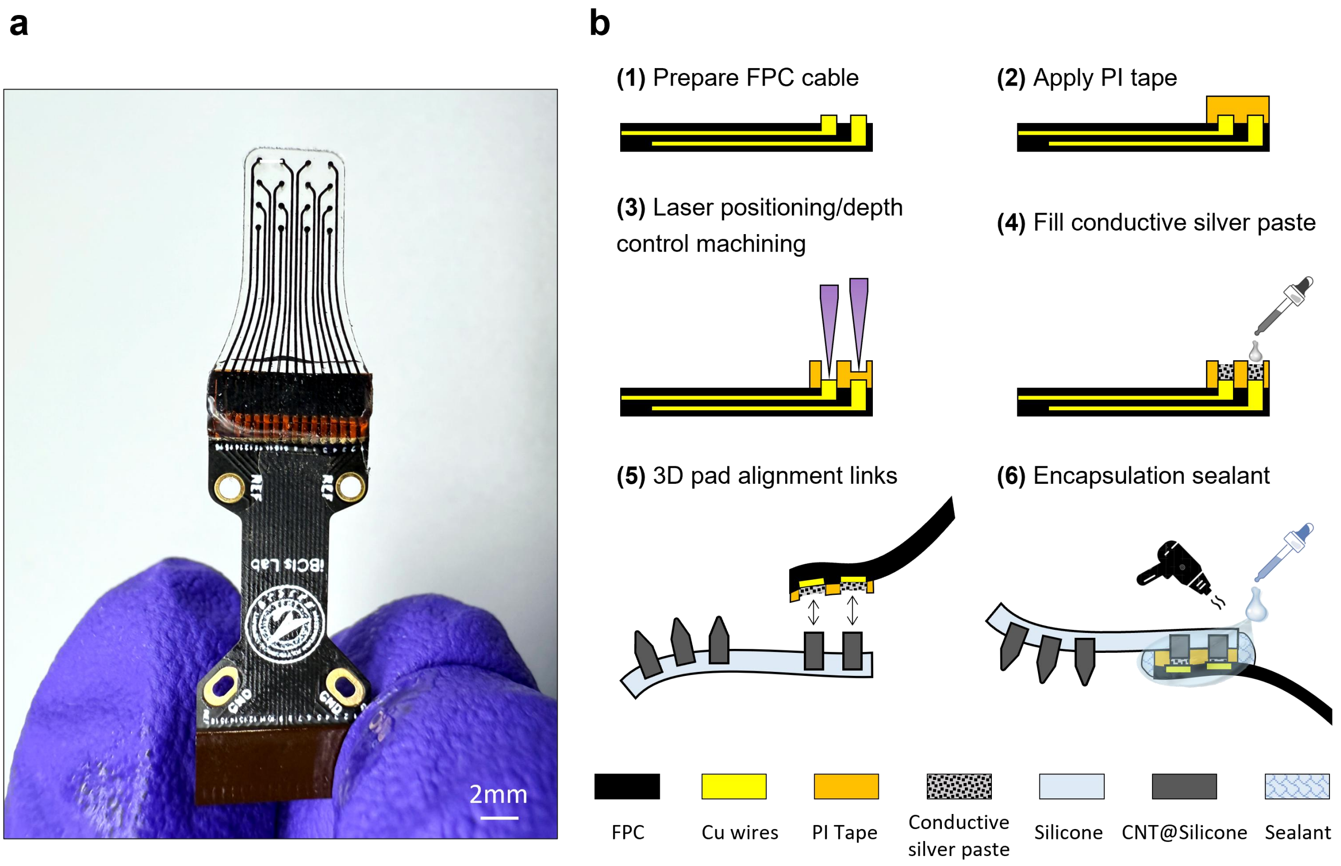


**Supplementary Fig. 12. Back-end interconnection strategy for 3D soft carbon electrode arrays. a,** Photograph of a 16-channel array connected to the flexible printed circuit (FPC) cable for external electrical interfacing. **b,** Schematic illustration of the 3D pad-based interconnection process.

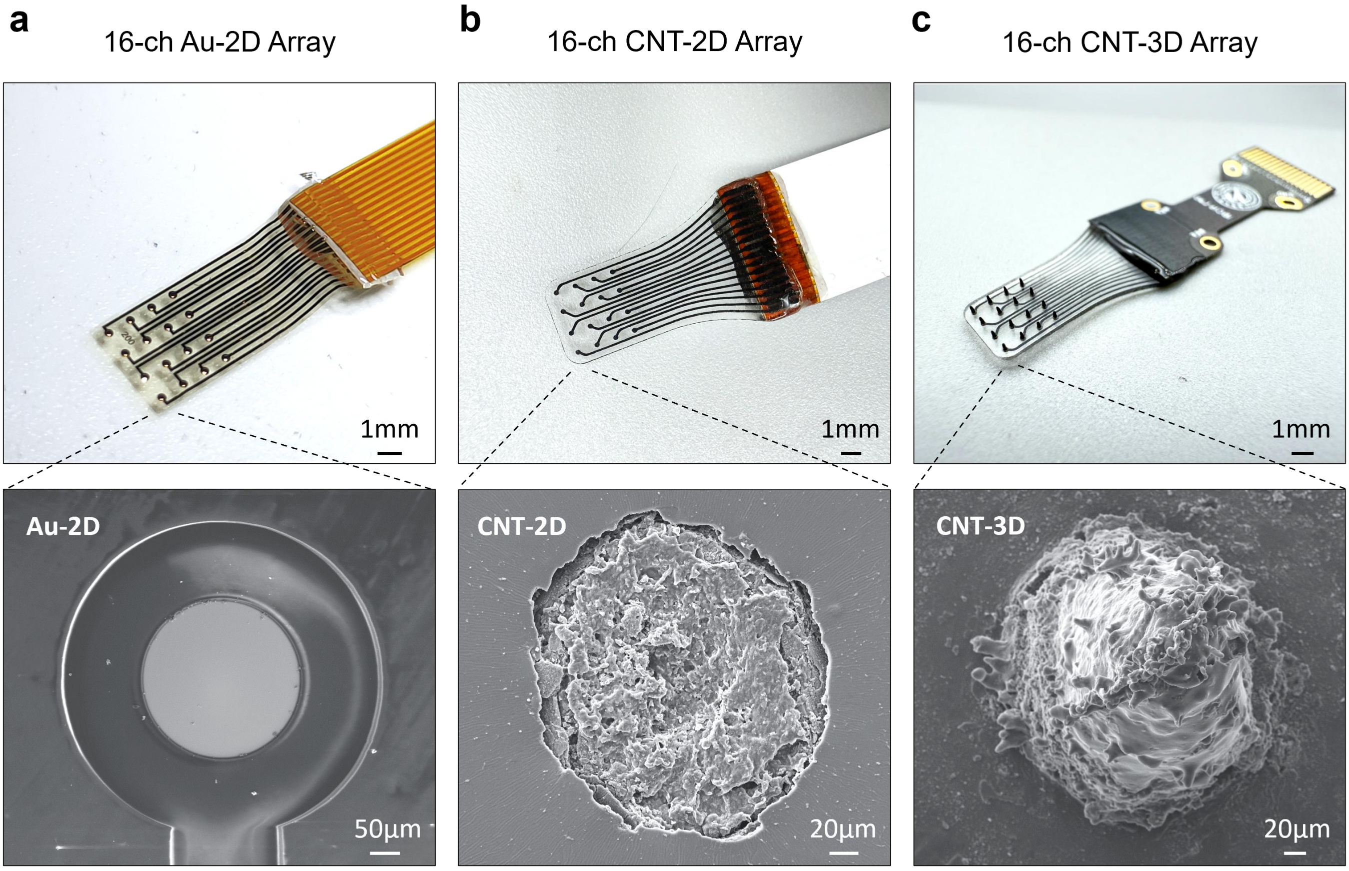


**Supplementary Fig. 13. Photographs of the three 16-channel electrode arrays used for comparative studies. a,** 16-channel Au-2D array with planar gold electrode sites. **b,** 16-channel CNT-2D array with planar CNTs@silicone electrode sites. **c,** 16-channel CNT-3D array with protruding 3D CNTs@silicone electrode sites. These devices were used for comparation platforms to evaluate the effects of electrode material and site geometry on electrochemical performance and in vivo ECoG recordings.

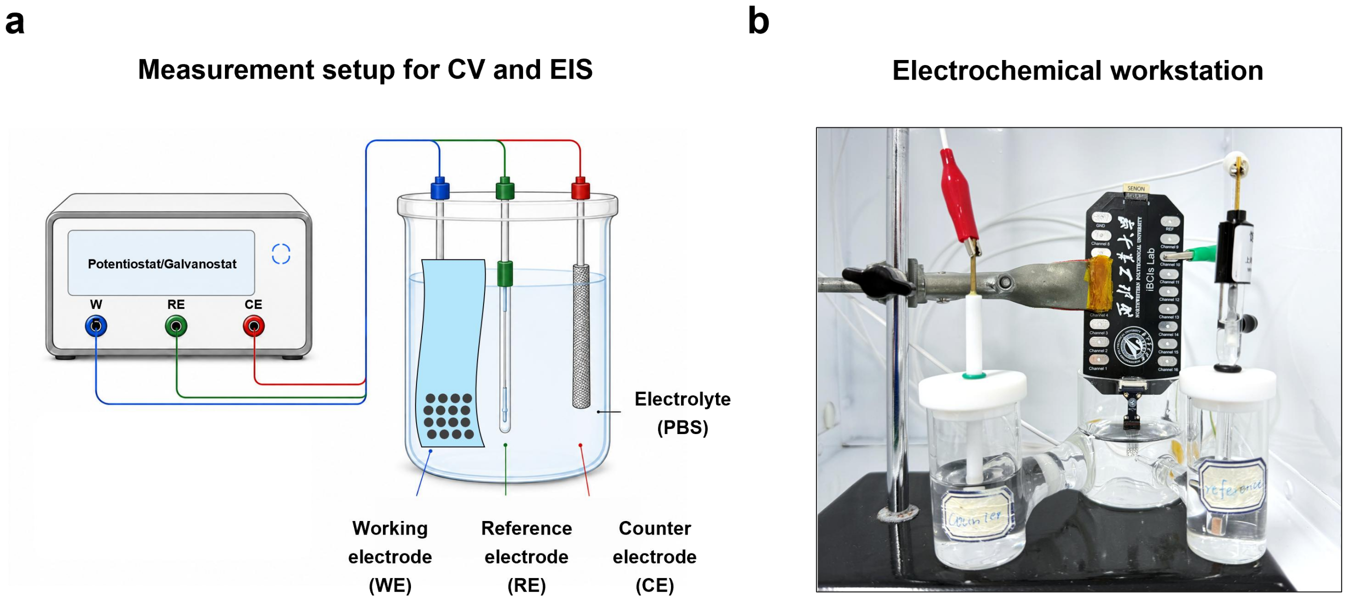


**Supplementary Fig. 14. Electrochemical characterization and measurement principle. a,** Wiring schematic of the three-electrode setup used for cyclic voltammetry (CV) and electrochemical impedance spectroscopy (EIS), in which the tested electrode served as the working electrode, with reference and counter electrodes completing the electrochemical circuit. **b,** Photograph of the electrochemical workstation.

**a**

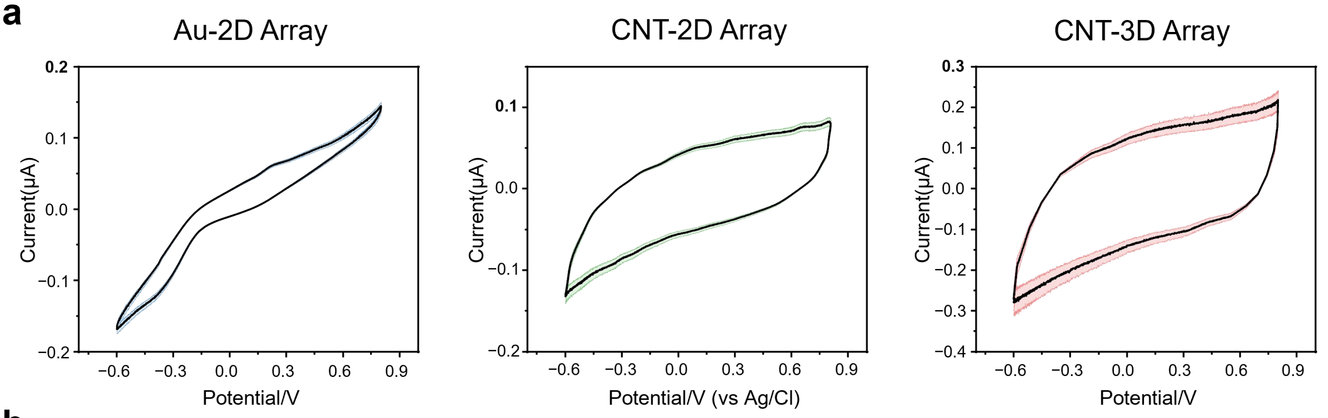


**b**

| Charge Storage Capacity - $CSC_c$ ($mC/cm^2$) | | | |
|---|---|---|---|
| **Channel No.** | **Au-2D** | **CNT-2D** | **CNT-3D** |
| Channel_1 | 2.48 | 2.71 | 6.42 |
| Channel_2 | 2.51 | 2.75 | 6.60 |
| Channel_3 | 2.51 | 2.76 | 6.61 |
| Channel_4 | 2.55 | 2.84 | 6.93 |
| Channel_5 | 2.71 | 3.11 | 7.84 |
| Channel_6 | 2.52 | 2.77 | 6.66 |
| Channel_7 | 2.42 | 2.59 | 5.93 |
| Channel_8 | 2.54 | 2.81 | 6.85 |
| Channel_9 | 2.59 | 2.91 | 7.22 |
| Channel_10 | 2.54 | 2.82 | 6.86 |
| Channel_11 | 2.40 | 2.56 | 5.74 |
| Channel_12 | 2.43 | 2.62 | 6.05 |
| Channel_13 | 2.59 | 2.91 | 7.23 |
| Channel_14 | 2.36 | 2.49 | 5.52 |
| Channel_15 | 2.34 | 2.47 | 5.43 |
| Channel_16 | 2.38 | 2.54 | 5.71 |

**c**

| | Mean ($mC/cm^2$) | Std Dev ($mC/cm^2$) | Min ($mC/cm^2$) | Max ($mC/cm^2$) | Coefficient of Variation (%) |
|---|---|---|---|---|---|
| **Au-2D** | 2.49 | 0.1 | 2.54 | 2.91 | ~3.6% |
| **CNT-2D** | 2.73 | 0.17 | 2.47 | 3.11 | ~6.2% |
| **CNT-3D** | 6.49 | 0.69 | 5.43 | 8.04 | ~10.6% |

**Supplementary Fig. 15**. **$CSC_c$ values of 16-channel Au-2D, CNT-2D and CNT-3D arrays. a,** Cyclic voltammetry curves measured from all 16 channels of the three arrays. **b,** Channel-wise $CSC_c$ values calculated from the corresponding CV curves. **c,** Summary comparison.

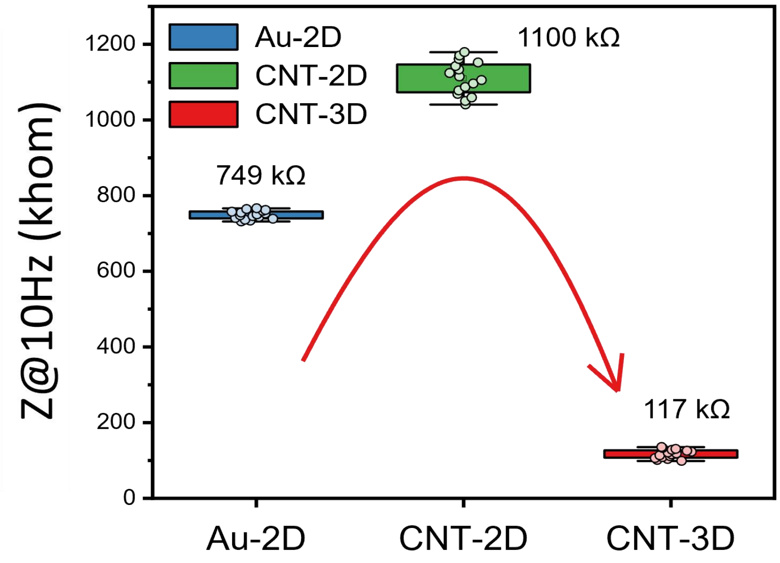


**Supplementary Fig. 16. Comparison of electrode impedance at 10 Hz for 16-channel Au-2D, CNT-2D and CNT-3D arrays.** Although 1 kHz impedance is commonly used as a standard benchmark for neural electrodes, low-frequency impedance is more representative of the electrode-electrolyte interface within the frequency range containing much of the ECoG activity analyzed in this study. Consistent with the trend observed at 1 kHz, the CNT-3D array exhibited substantially lower impedance than both planar electrode designs at 10 Hz.

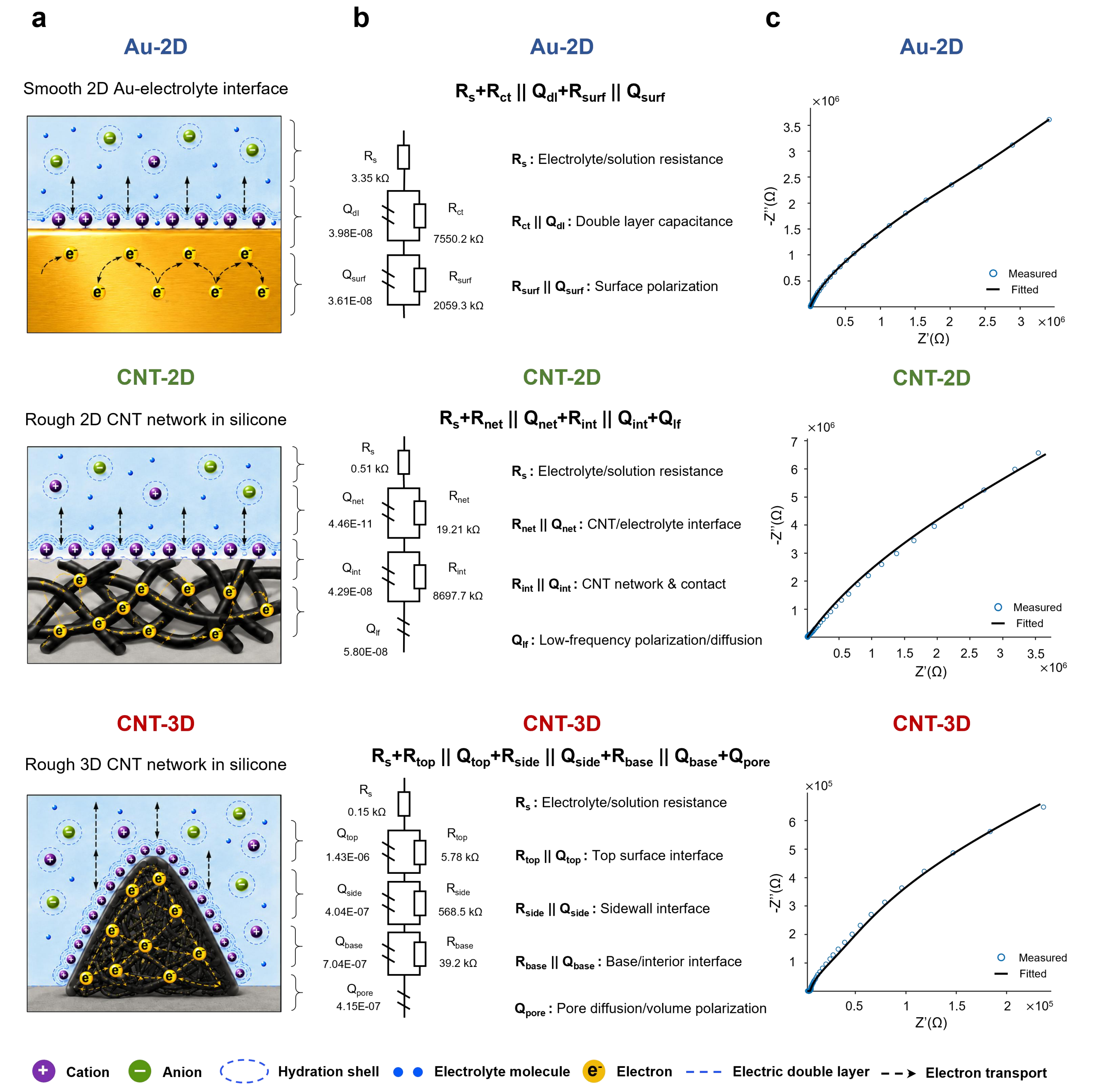


**Supplementary Fig. 17. Interfacial charge-transfer models and equivalent-circuit analysis of Au-2D, CNT-2D and CNT-3D electrodes. a,** Schematic illustration of the interfacial electron/ionic charge-conversion pathways for the three electrodes. **b,** Equivalent-circuit models used to describe the electrode-electrolyte interface together with the fitted circuit parameters. For Au-2D electrode, the model represents a planar Au/electrolyte interface with effective surface polarization. For CNT-2D electrode, the model reflects the two-dimensional CNT conductive network, the CNT/electrolyte interface and a low-frequency polarization contribution. For CNT-3D electrode, the model captures the spatially distributed interfacial responses arising from the cone top, sidewall and base/pore regions. **c,** Nyquist plots and corresponding equivalent-circuit fitting curves.

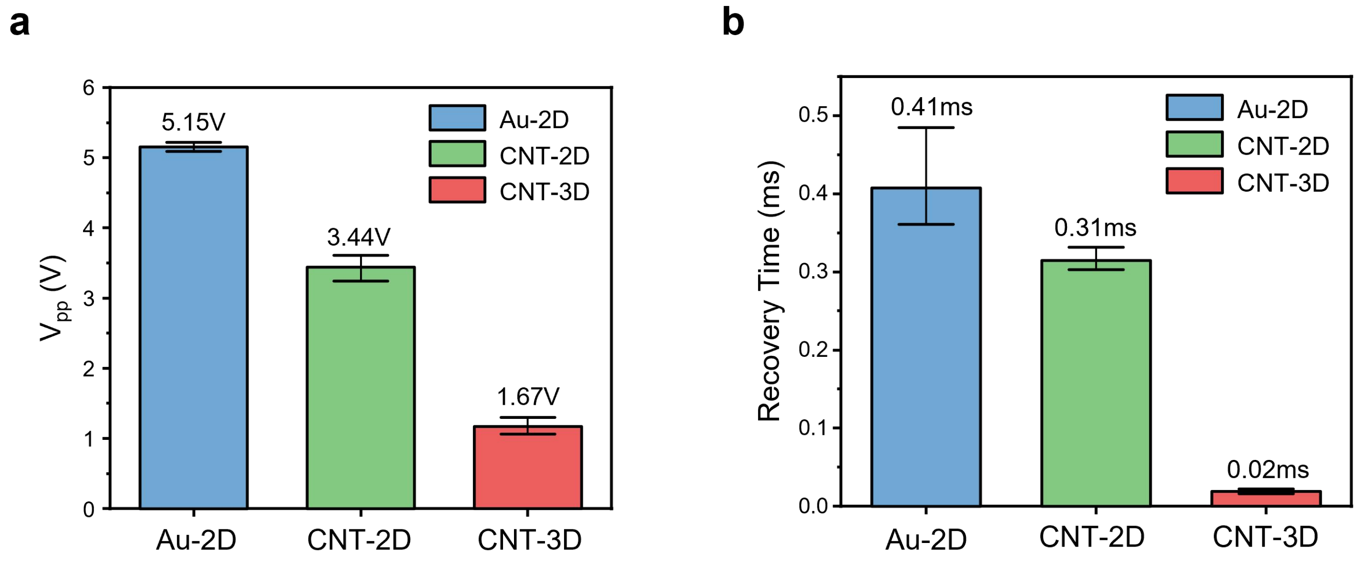


**Supplementary Fig. 18. Quantitative analysis of voltage-transient responses of Au-2D, CNT-2D and CNT-3D electrodes. a,** Comparison of the voltage peak-to-peak amplitude (Vpp) extracted from the voltage-transient responses under identical stimulation conditions. The Au-2D electrode exhibited the largest Vpp (5.15 V), followed by CNT-2D (3.44 V), whereas CNT-3D showed the smallest Vpp (1.67 V), indicating the lowest overall voltage demand during current pulsing. **b,** Comparison of recovery time, defined as the time required for the voltage to return toward baseline after stimulation. The Au-2D electrode showed the longest recovery time (0.41 ms), followed by CNT-2D (0.31 ms), whereas CNT-3D exhibited the shortest recovery time (0.02 ms). Together, these results indicate that the CNT-3D electrode provides the most efficient interfacial response, with reduced driving voltage and the fastest post-stimulation recovery.

**a**

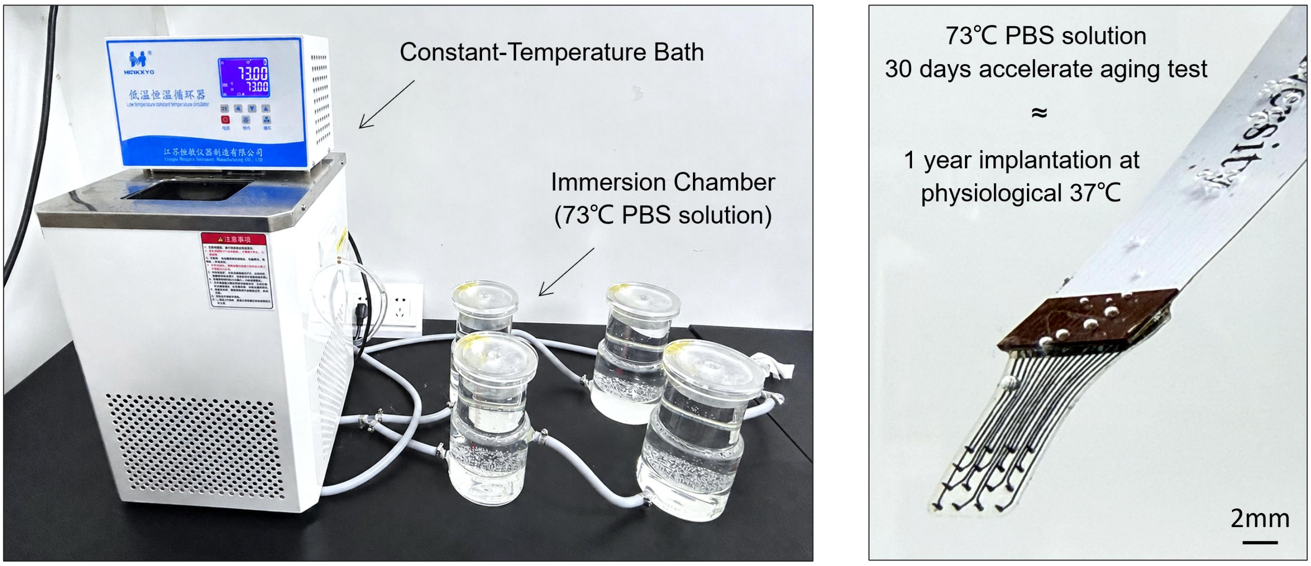


**b**

| Channel | \|Z\| @1kHz (kohm) | | | | | |
|---|---|---|---|---|---|---|
| | 0 day | 6 day | 12 day | 18 day | 24 day | 30 day |
| Channel 1 | 9.58 | 15.48 | 24.09 | 35.10 | 49.87 | 66.70 |
| Channel 2 | 9.34 | 15.16 | 20.38 | 39.96 | 60.73 | 73.08 |
| Channel 3 | 9.39 | 17.70 | 25.62 | 38.70 | 51.94 | 56.24 |
| Channel 4 | 9.54 | 16.62 | 22.69 | 44.18 | 48.50 | 73.82 |
| Channel 5 | 10.19 | 15.22 | 25.75 | 39.73 | 55.93 | 57.88 |
| Channel 6 | 9.89 | 13.90 | 27.10 | 33.08 | 45.79 | 48.36 |
| Channel 7 | 9.16 | 14.90 | 25.20 | 28.70 | 51.99 | 79.22 |
| Channel 8 | 9.74 | 15.50 | 23.77 | 39.58 | 52.59 | 52.79 |
| Channel 9 | 9.28 | 15.19 | 24.45 | 33.32 | 54.86 | 71.90 |
| Channel 10 | 9.88 | 13.84 | 23.61 | 37.02 | 43.03 | 61.96 |
| Channel 11 | 10.17 | 17.66 | 24.74 | 35.05 | 56.39 | 72.84 |
| Channel 12 | 9.40 | 14.24 | 22.41 | 31.92 | 58.39 | 76.10 |
| Channel 13 | 9.59 | 15.21 | 28.45 | 33.96 | 47.21 | 65.27 |
| Channel 14 | 9.90 | 16.67 | 25.62 | 32.71 | 54.82 | 83.56 |
| Channel 15 | 9.29 | 15.01 | 26.17 | 41.65 | 51.63 | 69.91 |
| Channel 16 | 10.02 | 20.21 | 24.98 | 34.28 | 49.09 | 57.04 |
| **Average** | **9.65 kohm** | **15.78 kohm** | **24.38 kohm** | **36.06 kohm** | **51.17 kohm** | **67.48 kohm** |

**Supplementary Fig. 19. Accelerated aging test of the 16-channel CNT-3D array. a,** Photograph of the immersed array in PBS at 73 °C. **b,** Summary of 1 kHz impedance values for all 16 channels, together with the channel-averaged impedance, measured at days 0, 6, 12, 18, 24 and 30. According to a $Q_{10} = 2$ accelerat ageing model, immersion in PBS at 73 °C for 30 days corresponds to approximately 364 days of implantation at physiological temperature ($\boldsymbol{Equivalent\ Implantation\ Time = Immerse\ Time \times Q10^{(T_{PBS}-T_{Body})/10}}$). The mean 1 kHz impedance increased from 9.65 kΩ to 67.5 kΩ after ageing, remaining below 100 kΩ. No open-circuit failure or abrupt impedance increase was observed across the 16 channels.

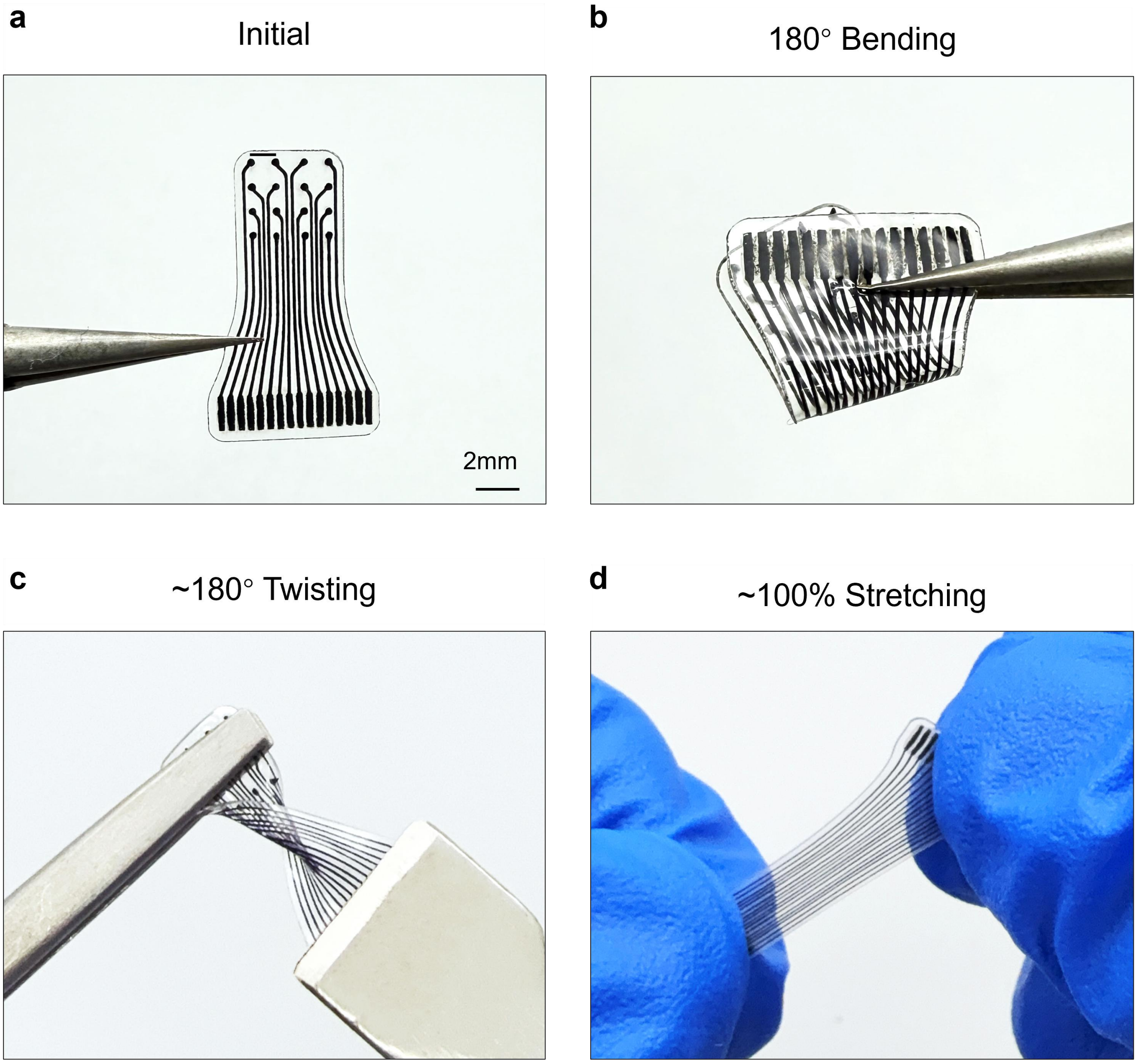


**Supplementary Fig. 20. Mechanical deformability of the 16-channel CNT-3D array.** Representative photographs of the CNT-3D array under different deformation states, including the initial state **(a),** 180° bending **(b),** 180° twisting **(c)**, and 100% stretching **(d).**

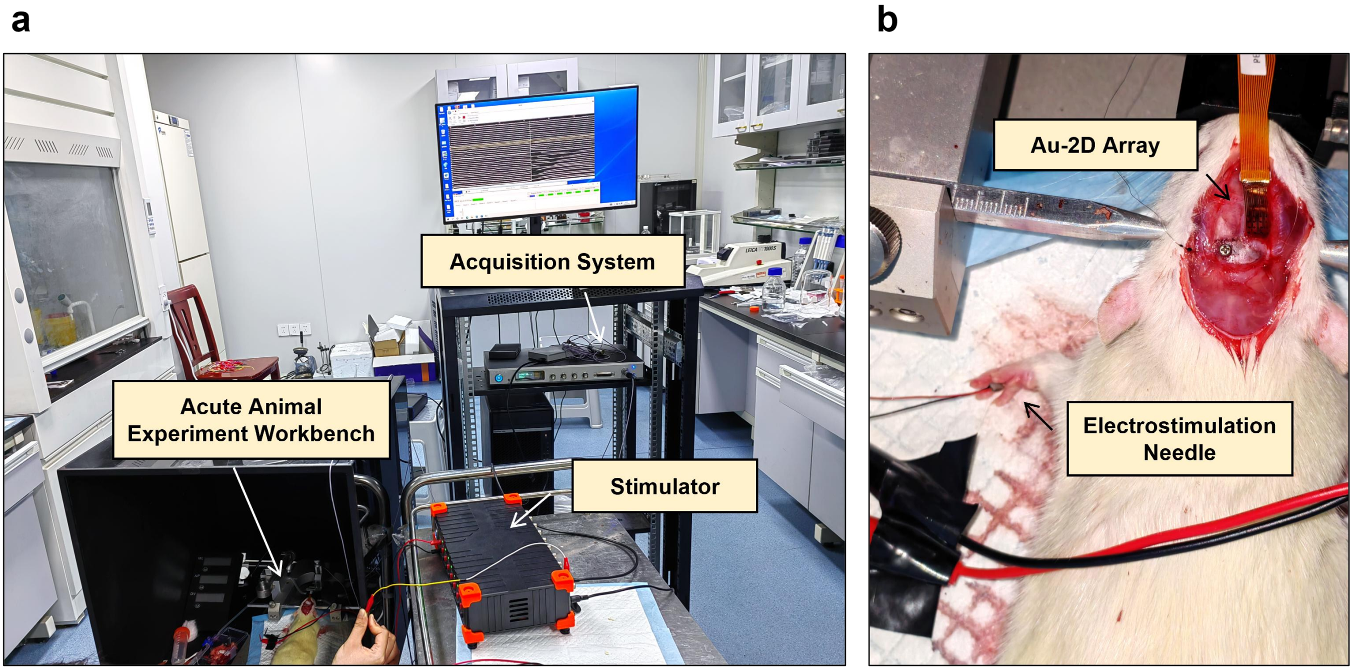


**Supplementary Fig. 21. The rat forepaw stimulation-acquisition experiment. a,** Photograph of the acute animal experimental setup, including the surgical workbench, signal acquisition system and electrical stimulator. **b,** Top-view photograph of the experimental rat, showing stimulation needles inserted into the left forepaw and the Au-2D array placed on the right cortical surface.

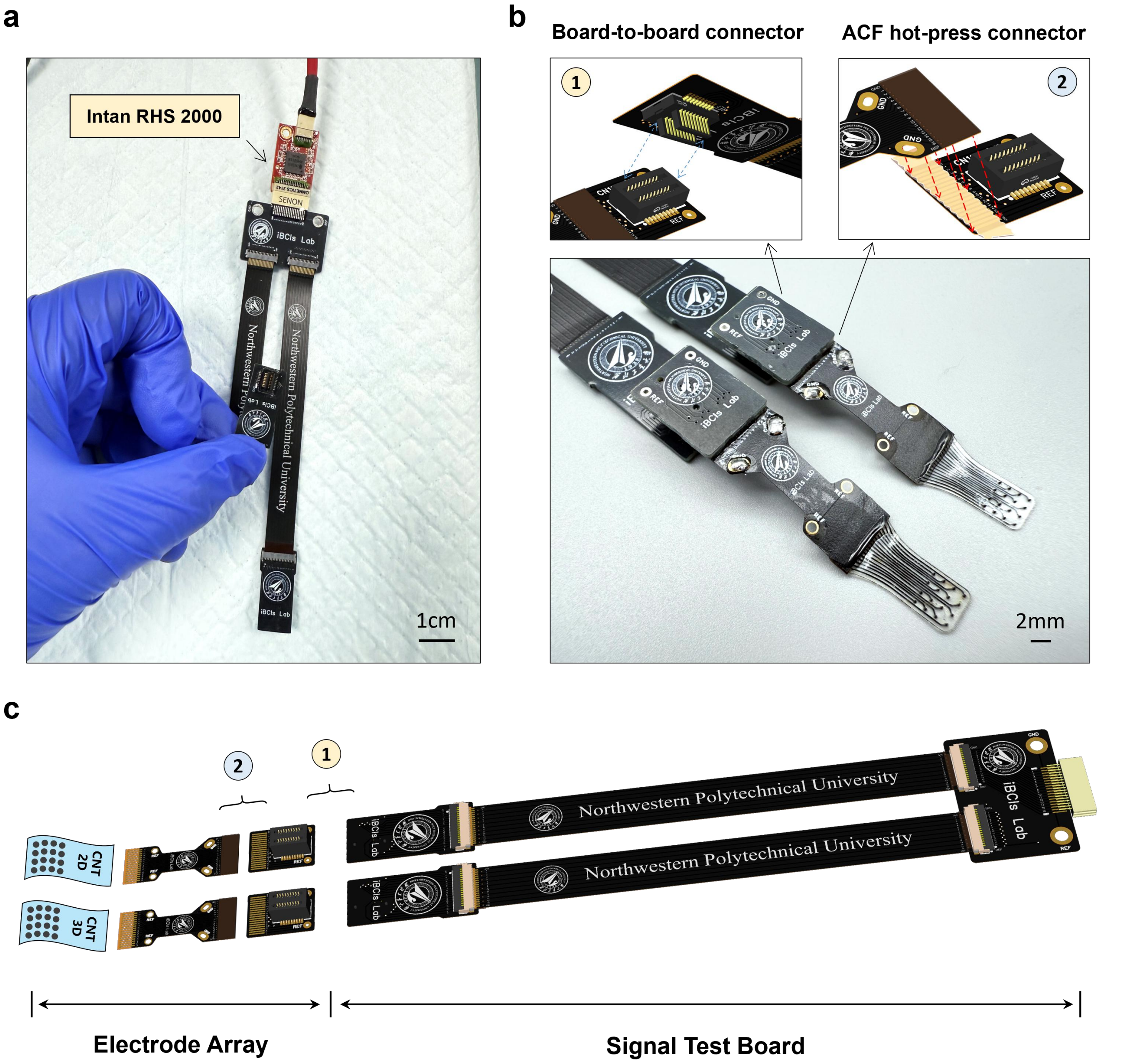


**Supplementary Fig. 22. The circuit design for simultaneous bilateral test with CNT-3D and CNT-2D arrays. a,** Photograph of the bilateral signal test board connected to the Intan RHS 2000 chip. **b,** Enlarged view of the two interconnection strategies integrated on the board, including a board-to-board connector and an anisotropic conductive film (ACF) hot-press connector. **c,** Three-dimensional model of the complete circuit board.

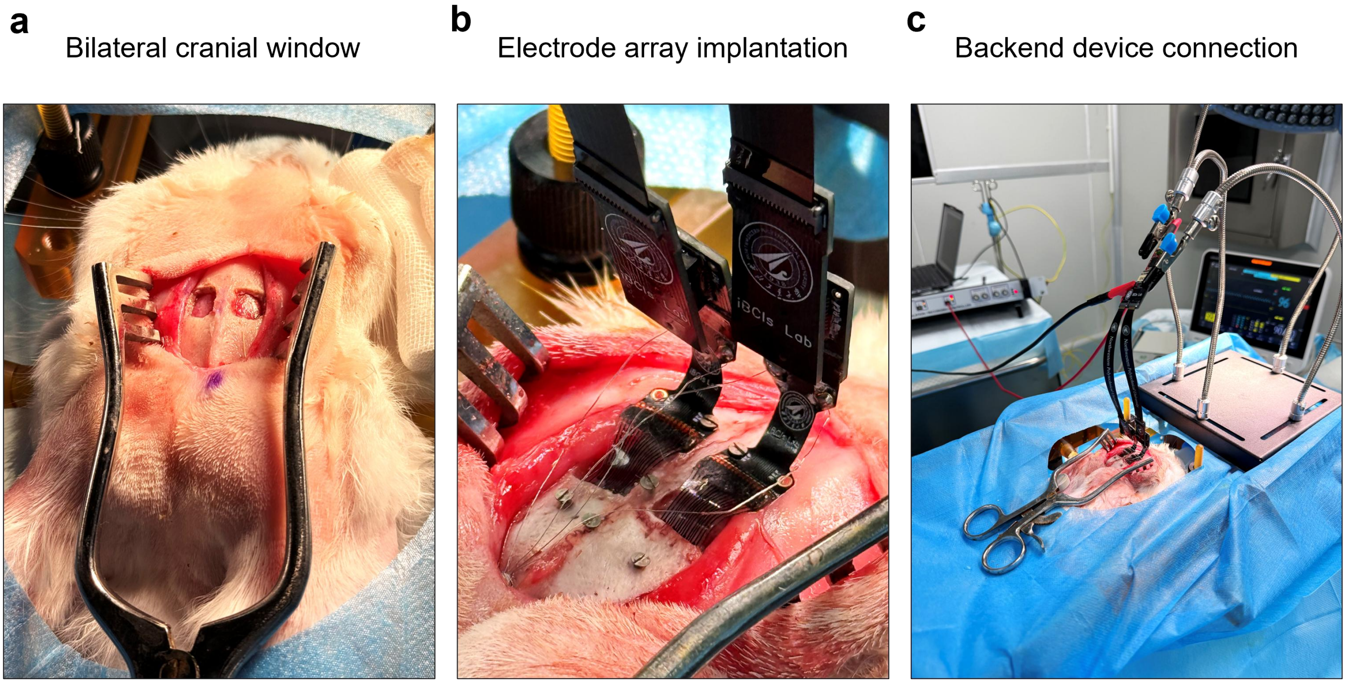


**Supplementary Fig. 23. Surgical implantation procedure for bilateral cortical recording in New Zealand white rabbit. a,** Top-view photograph of the bilateral craniotomy windows opened over the left and right primary motor cortex (M1), with the cortical surface exposed. **b,** Photograph showing the implantation of CNT-2D and CNT-3D arrays into the left and right cranial windows, respectively, together with platinum-iridium wire connections to the reference (REF) and ground (GND) Ti screws. **c,** Photograph of the implanted arrays connected to the back-end bilateral signal test board, which was further interfaced with the Intan system for in vivo impedance measurement.

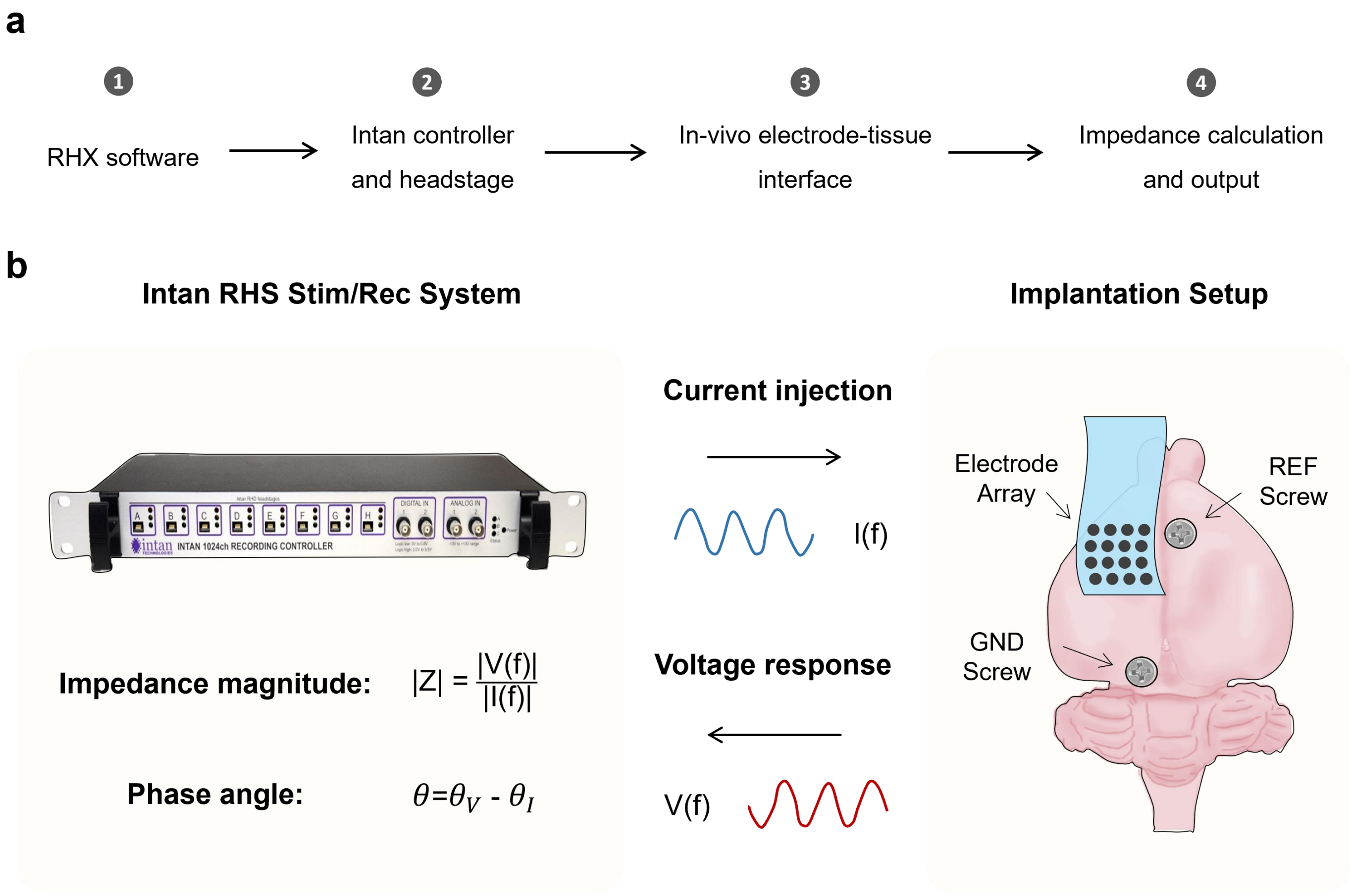


**Supplementary Fig. 24. The in vivo impedance measurement based on the Intan RHS 2000 system.** **a,** Schematic overview of the in vivo impedance-testing workflow. **b,** Schematic illustration of the measurement principle. The Intan RHS Stim/Rec system applies a sinusoidal current, *I(f)*, sequentially to each channel of the implanted electrode array and records the corresponding voltage response, *V(f)*, with respect to the reference (REF) and ground (GND) bone screws. The in vivo impedance magnitude and phase were then calculated as $|Z| = |V(f)|/ |I(f)|$, $\boldsymbol{\theta} = \boldsymbol{\theta_V} - \boldsymbol{\theta_I}$, respectively. The right panel illustrates the relative positions of the implanted electrode array, REF screw and GND screw on the rabbit skull.

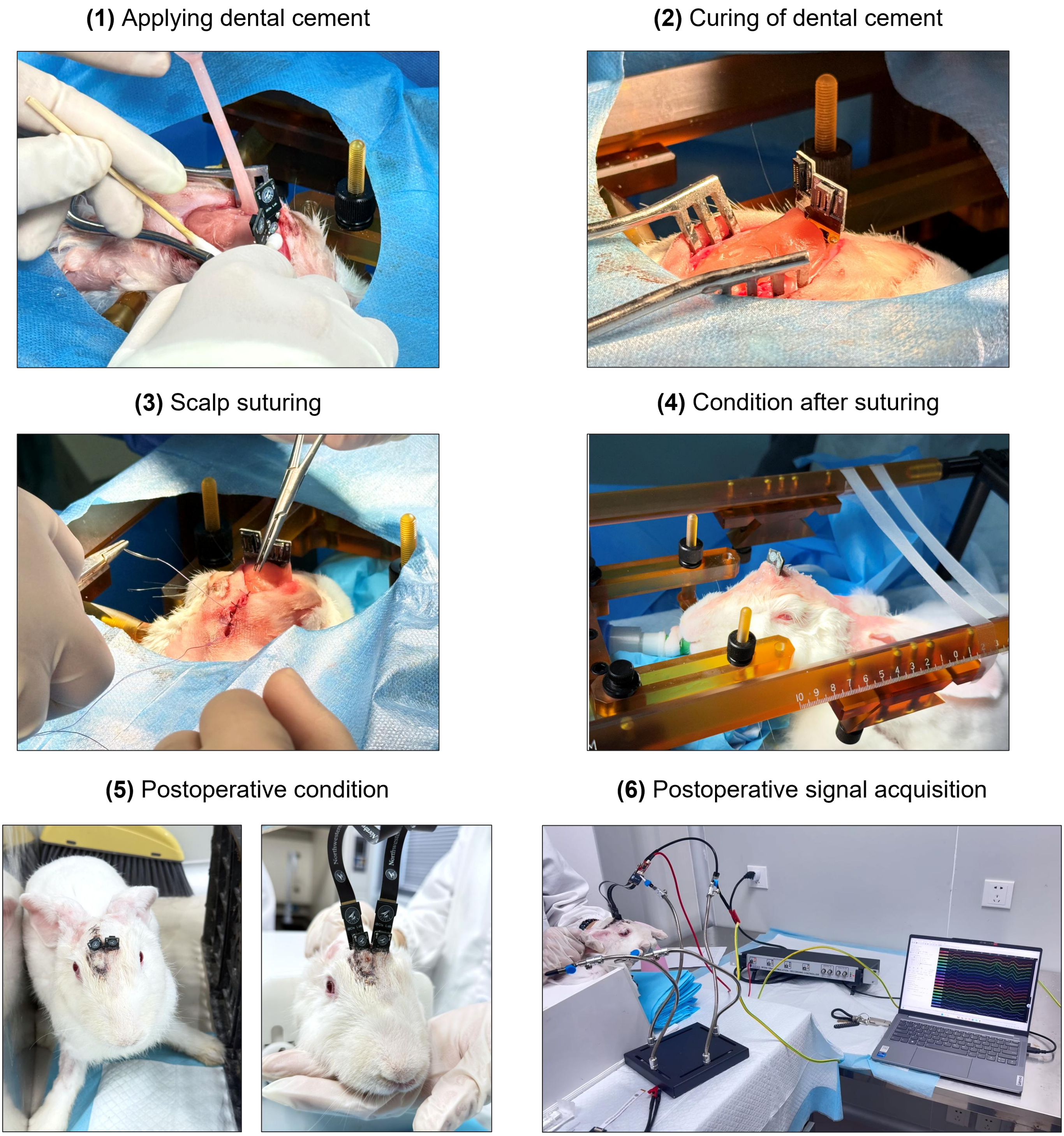


**Supplementary Fig. 25. Post-implantation fixation, wound closure and postoperative ECoG acquisition.**

**a**

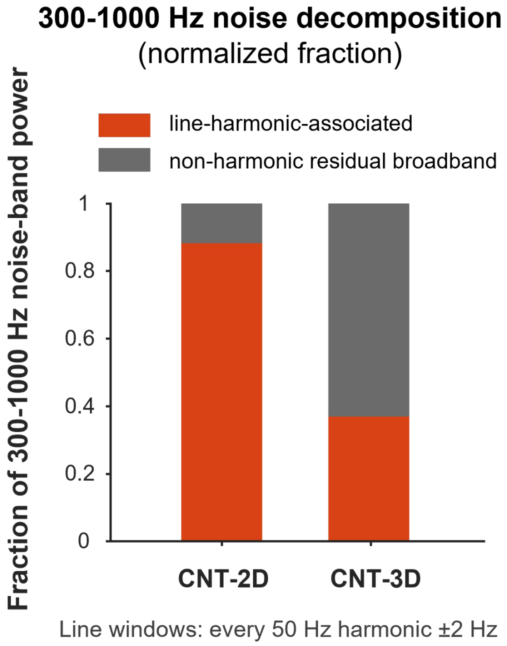


**b**

| Indicators | CNT-2D | CNT-3D |
|---|---|---|
| Total HF power (300-1000 Hz) | 24.55 dB | 15.04 dB |
| Line-harmonic-associated power | 20.53 dB | 4.98 dB |
| Residual broadband power | 22.36 dB | 14.58 dB |
| Line-harmonic fraction | 39.7% | 10.0% |

**Supplementary Fig. 26. Decomposition of high-frequency noise components in CNT-2D and CNT-3D cortical signals. a,** Noise decomposition plot in the 300-1000 Hz band shown as normalized fractions, separating line-harmonic-associated and residual broadband components. Line windows were defined as ±2 Hz around each 50 Hz harmonic. **b,** Quantitative comparison of high-frequency noise metrics in the 300-1000 Hz range for CNT-2D and CNT-3D arrays, including total high-frequency (HF) power, line-harmonic-associated power, residual broadband power and the line-harmonic fraction.

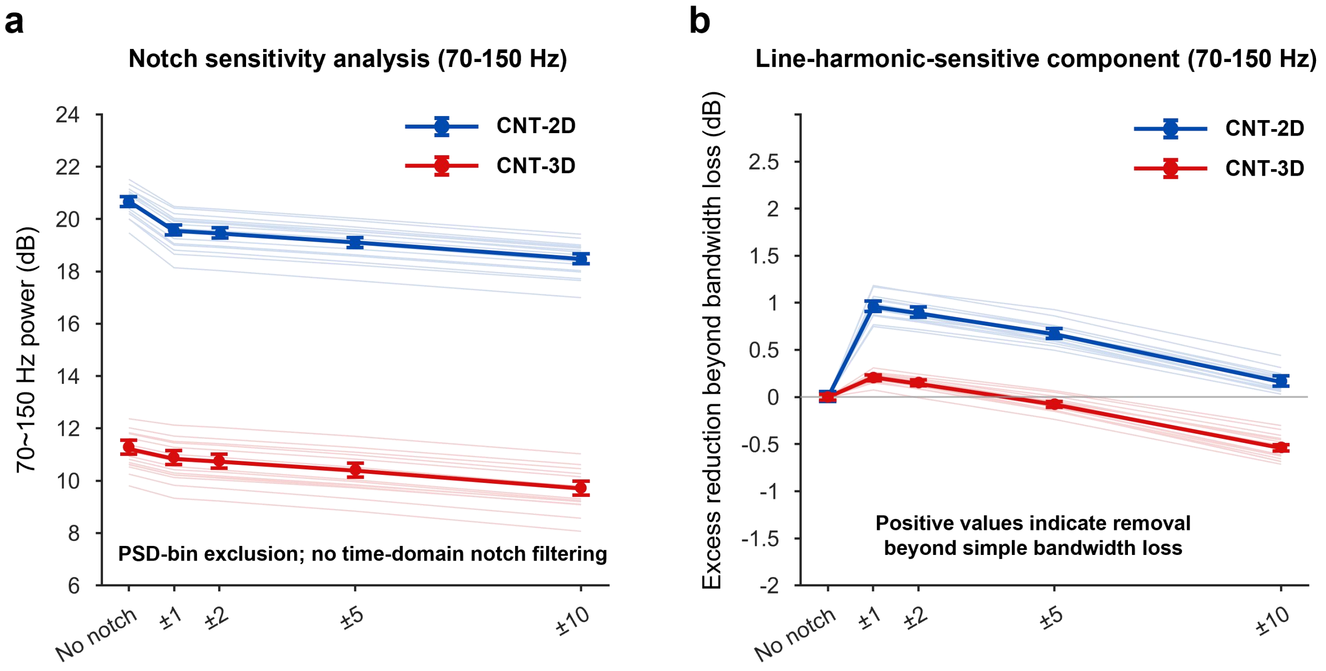


**c**

| Exclusion width | CNT-2D | CNT-3D | 2D-3D Difference |
|---|---|---|---|
| No exclusion | 20.68 dB | 11.21 dB | **9.47 dB** |
| ±1 Hz | 19.55 dB | 10.83 dB | **8.72 dB** |
| ±2 Hz | 19.45 dB | 10.73 dB | **8.72 dB** |
| ±5 Hz | 19.11 dB | 10.39 dB | **8.72 dB** |
| ±10 Hz | 18.48 dB | 9.71 dB | **8.77 dB** |

**Supplementary Fig. 27. Line-harmonic sensitivity analysis of the apparent 70-150 Hz power in CNT-2D and CNT-3D recordings. a,** Notch-sensitivity analysis of the 70-150 Hz band, showing how the measured band power changes as progressively wider exclusion windows are applied around power-line harmonics. **b,** Quantification of the line-harmonic-sensitive component within the 70-150 Hz range, highlighting the contribution of harmonic contamination to the apparent high-gamma-band power. **c,** Corresponding numerical summary of the notch-sensitivity analysis for different harmonic exclusion widths (no exclusion, ±1 Hz, ±2 Hz, ±5 Hz and ±10 Hz). As the harmonic exclusion width increases, the 70-150 Hz power of CNT-2D decreases more markedly than that of CNT-3D, demonstrating that the apparent elevation of the 70-150 Hz signal in CNT-2D contains a stronger line-harmonic-sensitive component.

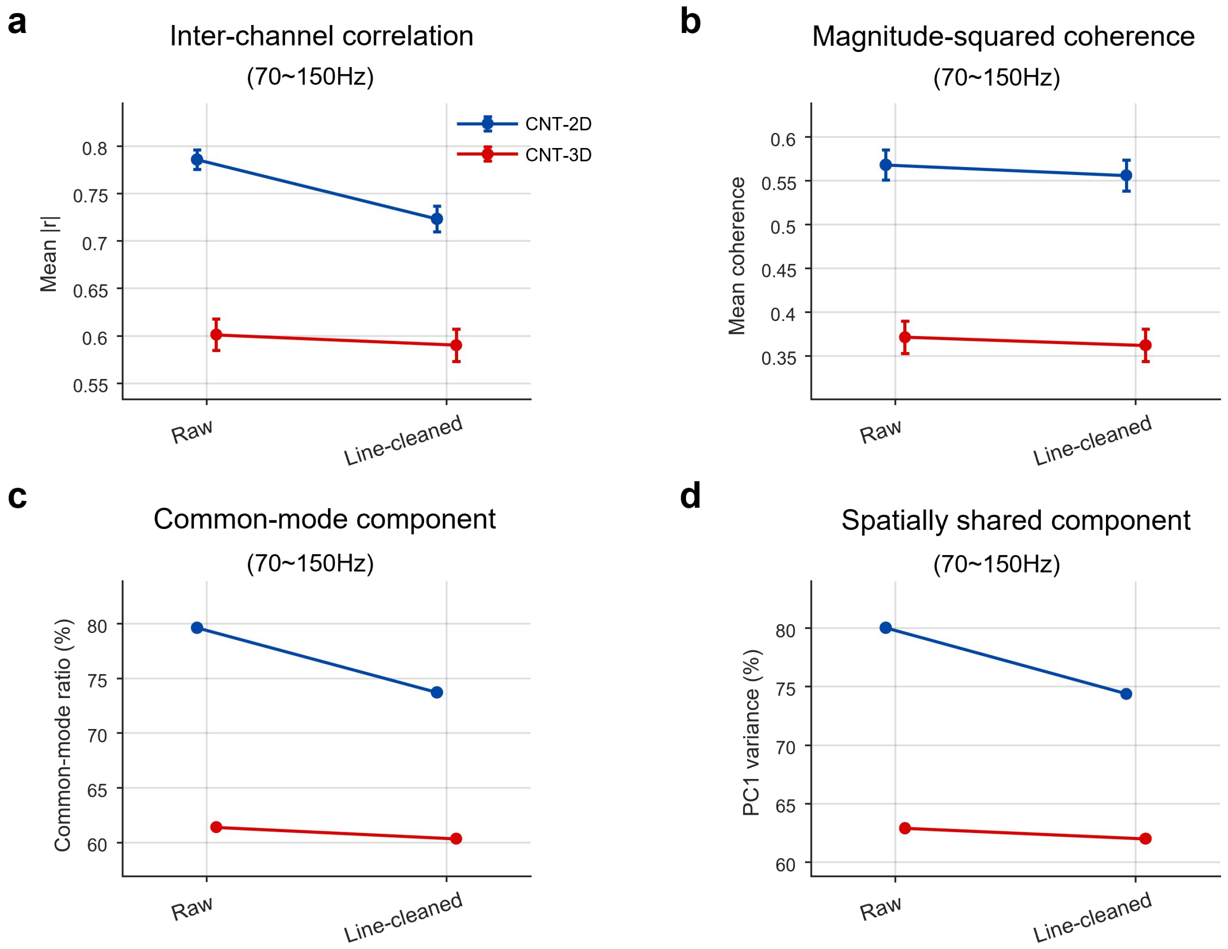


**e**

| Indicators | CNT-2D Raw | CNT-3D Raw | CNT-2D Line-cleaned | CNT-3D Line-cleaned |
|---|---|---|---|---|
| Mean abs correlation | 0.786 | 0.601 | 0.723 | 0.590 |
| Mean coherence | 0.568 | 0.371 | 0.556 | 0.362 |
| Common-mode ratio | 79.6% | 61.4% | 73.7% | 60.4% |
| PC1 variance fraction | 80.0% | 62.9% | 74.4% | 62.0% |

**Supplementary Fig. 28. Synchrony analysis of raw and line-cleaned 70-150 Hz activity. a,** Inter-channel correlation of raw and line-cleaned 70-150 Hz signals from CNT-2D and CNT-3D recordings. **b,** Magnitude-squared coherence analysis. **c,** Common-mode component analysis. **d,** Spatially shared component analysis based on the first principal component. **e,** Summary of synchrony metrics. Raw signals were analysed without notch filtering, whereas line-cleaned signals were processed to remove narrowband components around 100 and 150 Hz. CNT-2D exhibited stronger inter-channel correlation, coherence, common-mode contribution and spatially shared variance than CNT-3D in both raw and line-cleaned conditions, indicating that its apparent 70-150 Hz enhancement contains spatially diffuse, multi-channel-synchronized high-frequency contamination rather than localized physiological high-gamma activity.

**a**

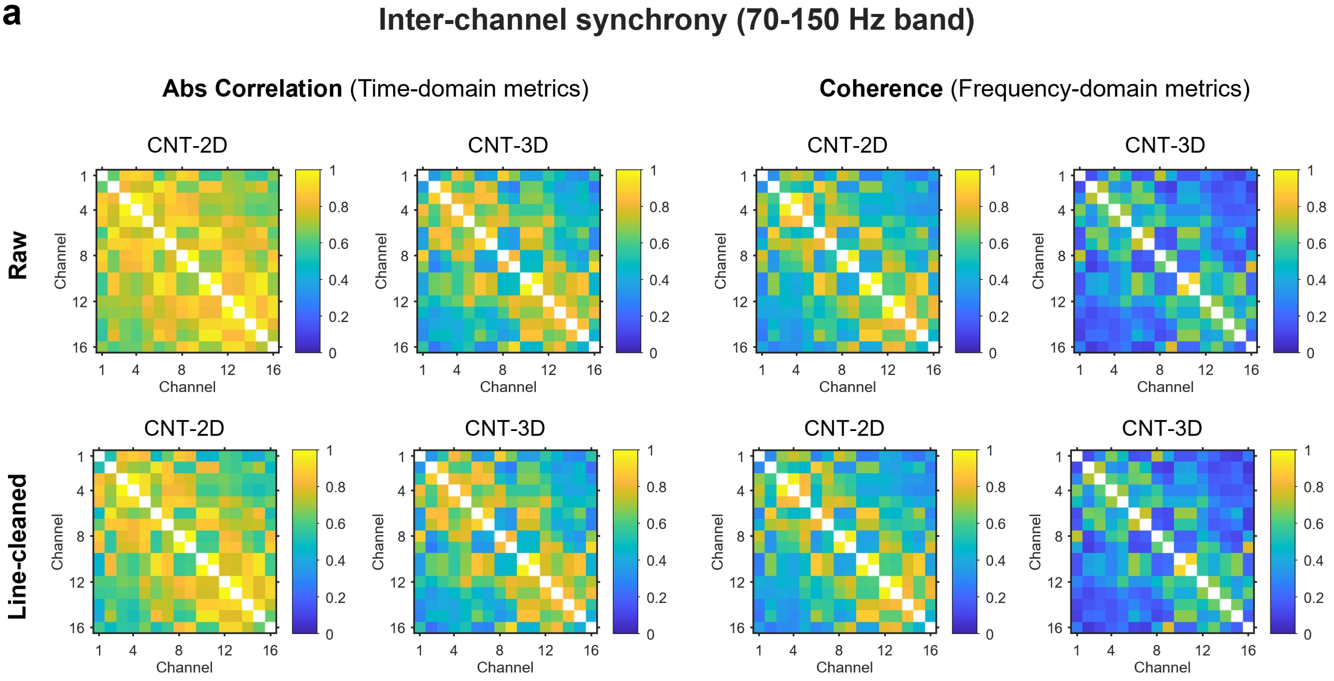


**b**

| | Threshold | Channel Pairs (CNT-2D) | Channel Pairs (CNT-3D) |
|---|---|---|---|
| Raw | abs correlation > 0.5 | 100% | 62.5% |
| | coherence > 0.5 | 55.0% | 30.8% |
| Line-cleaned | abs correlation > 0.5 | 95.0% | 62.5% |
| | coherence > 0.5 | 53.3% | 27.5% |

**Supplementary Fig. 29. Inter-channel synchrony analysis. a,** Heat maps of inter-channel synchrony in the 70-150 Hz band for CNT-2D and CNT-3D recordings. **b,** Summary table showing the proportion of channel pairs exceeding predefined synchrony thresholds for absolute correlation and coherence in the raw and line-cleaned conditions. CNT-2D consistently exhibited a higher fraction of strongly synchronized channel pairs than CNT-3D, both before and after line-harmonic removal.

**a**

**0.5-50 Hz band-pass trace**

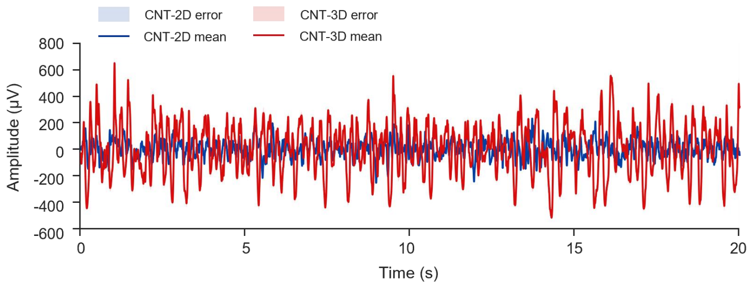


| Indicators | CNT-2D | CNT-3D |
|---|---|---|
| RMS | 65.7 µV | 203.0 µV |
| robust P2P | 208.9 µV | 664.5 µV |
| Spectrogram mean PSD | 18.88 dB | 29.23 dB |

**b**

**70-150 Hz band-pass trace**

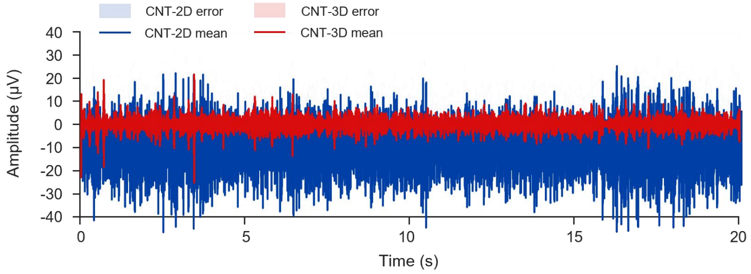


| Indicators | CNT-2D | CNT-3D |
|---|---|---|
| RMS | 9.06 µV | 2.68 µV |
| robust P2P | 29.7 µV | 7.91 µV |
| envelope mean | 12.60 µV | 3.56 µV |

**c**

**70-150 Hz amplitude envelope**

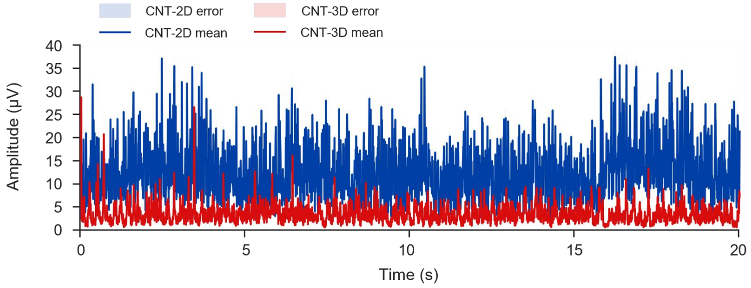


| Indicators | CNT-2D | CNT-3D |
|---|---|---|
| Envelope CV over time | 0.435 | 0.682 |
| Envelope P95/P50 | 1.95 | 2.42 |
| 10-s bin CV | 2.95% | 4.29% |

**Supplementary Fig. 30. Band-limited waveform analysis used to distinguish valid low-frequency ECoG signals from apparent high-frequency enhancement. a,** Average 0.5-50 Hz band-pass traces across 16 channels for CNT-2D and CNT-3D arrays, together with a summary table of trace RMS, robust peak-to-peak amplitude (robust P2P) and spectrogram mean PSD. **b,** Average 70-150 Hz band-pass traces across 16 channels. **c,** Average 70-150 Hz amplitude envelopes across 16 channels, together with summary metrics including envelope coefficient of variation over time (Envelope CV over time), envelope P95/P50 and 10-s bin CV.

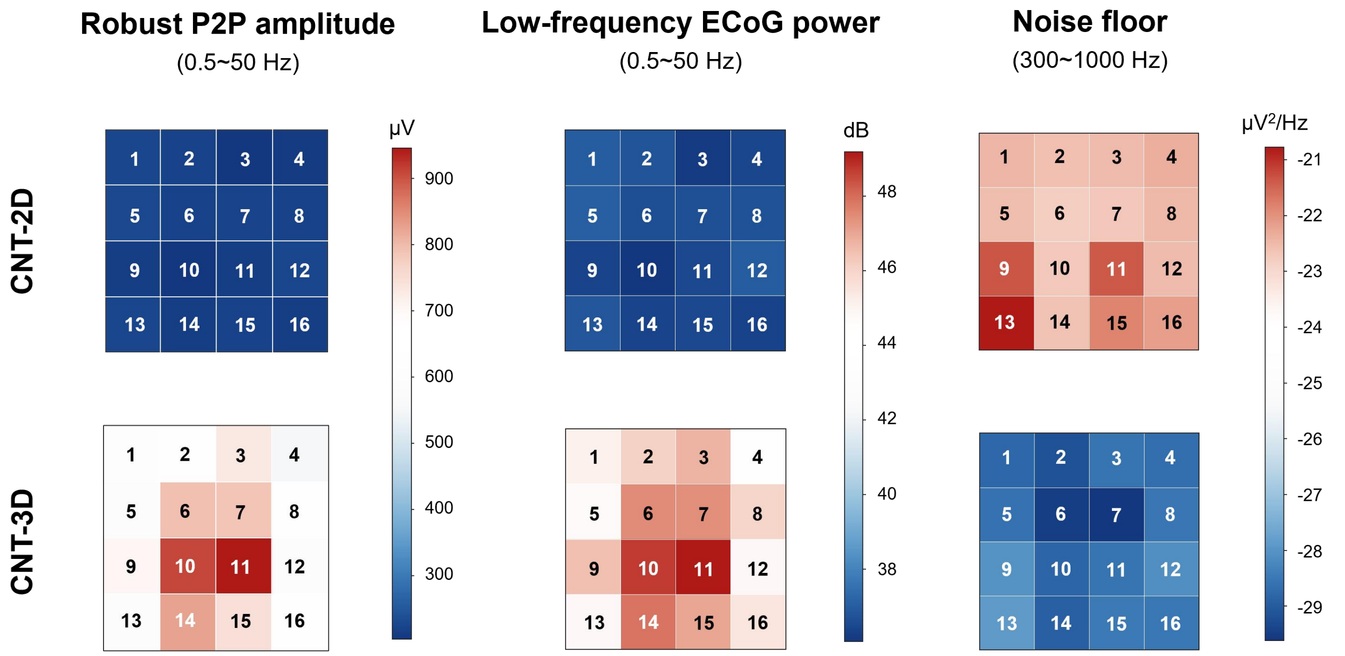


**Supplementary Fig. 31. Spatial comparison of signal amplitude, low-frequency power and noise floor between CNT-2D and CNT-3D arrays.** The heat maps comparing the 16-channel recording performance of CNT-2D and CNT-3D arrays, corresponding to the waveform comparison in Fig. 5b and the SNR analysis in Fig. 5f.

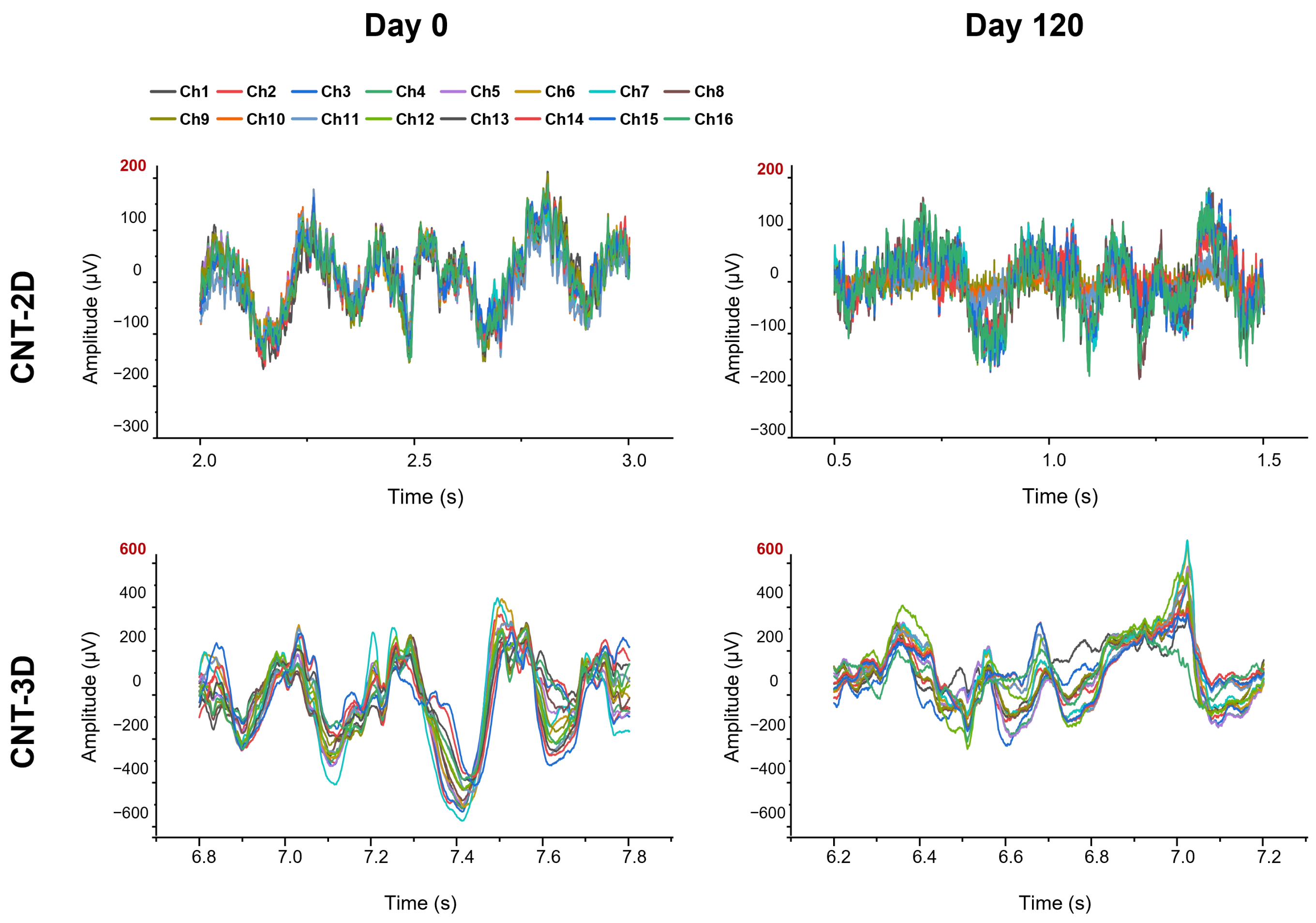


**Supplementary Fig. 32. Representative 16-channel ECoG recordings at Day 0 and 120.**

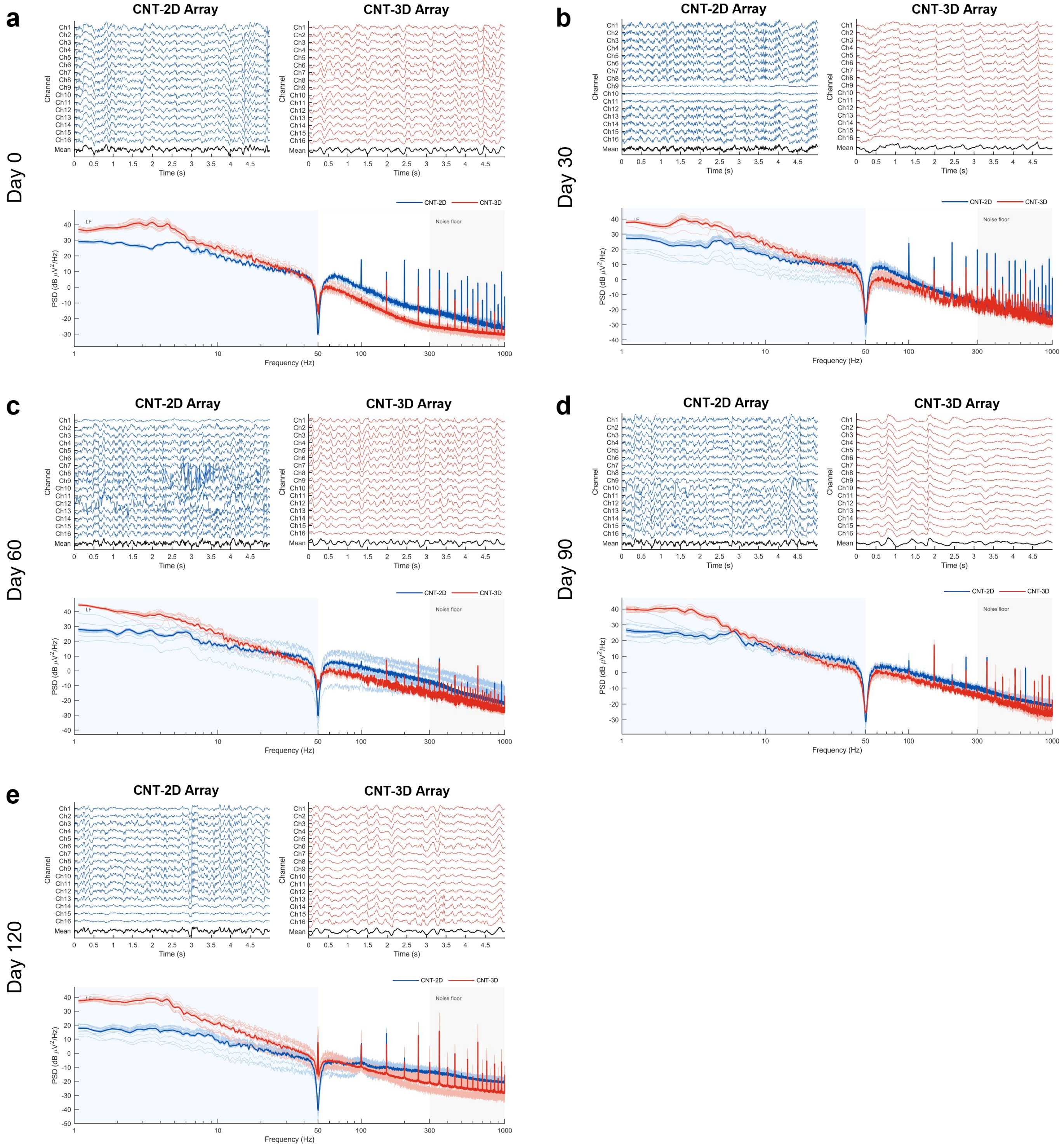


**Supplementary Fig. 33. Representative 16-channel ECoG traces and corresponding PSD curves recorded from CNT-2D and CNT-3D arrays.**

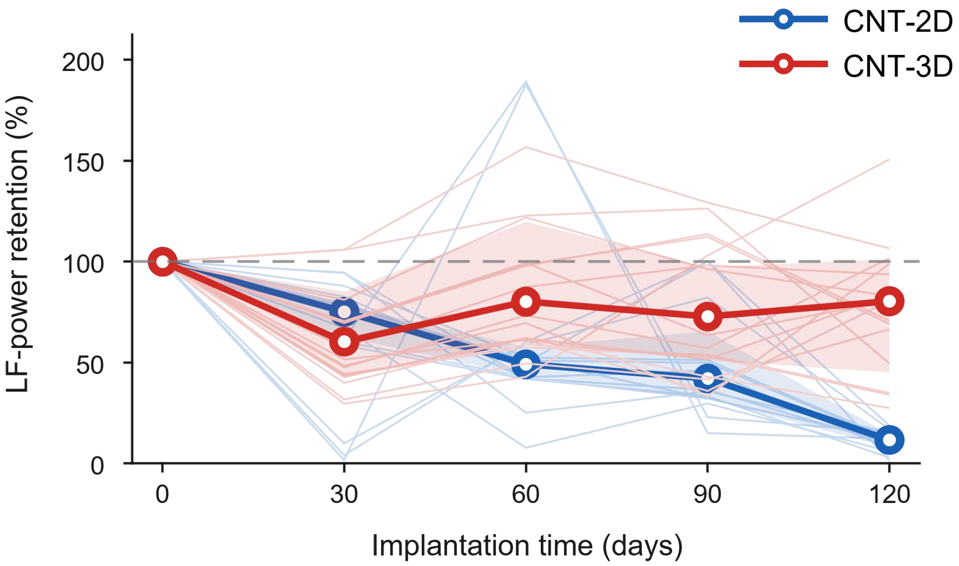


**Supplementary Fig. 34. Low-frequency (LF, 0.5-50 Hz) power retention relative to Day 0 over the 120-day implantation period for CNT-2D and CNT-3D arrays.** The LF power of CNT-2D progressively declined from 75.2% at Day 30 to 11.7% at Day 120, indicating substantial loss of the initial effective ECoG signal power over time. By contrast, CNT-3D showed a transient decrease to 60.4% at Day 30, followed by recovery, and retained approximately 80.4% of its Day 0 LF power at Day 120.

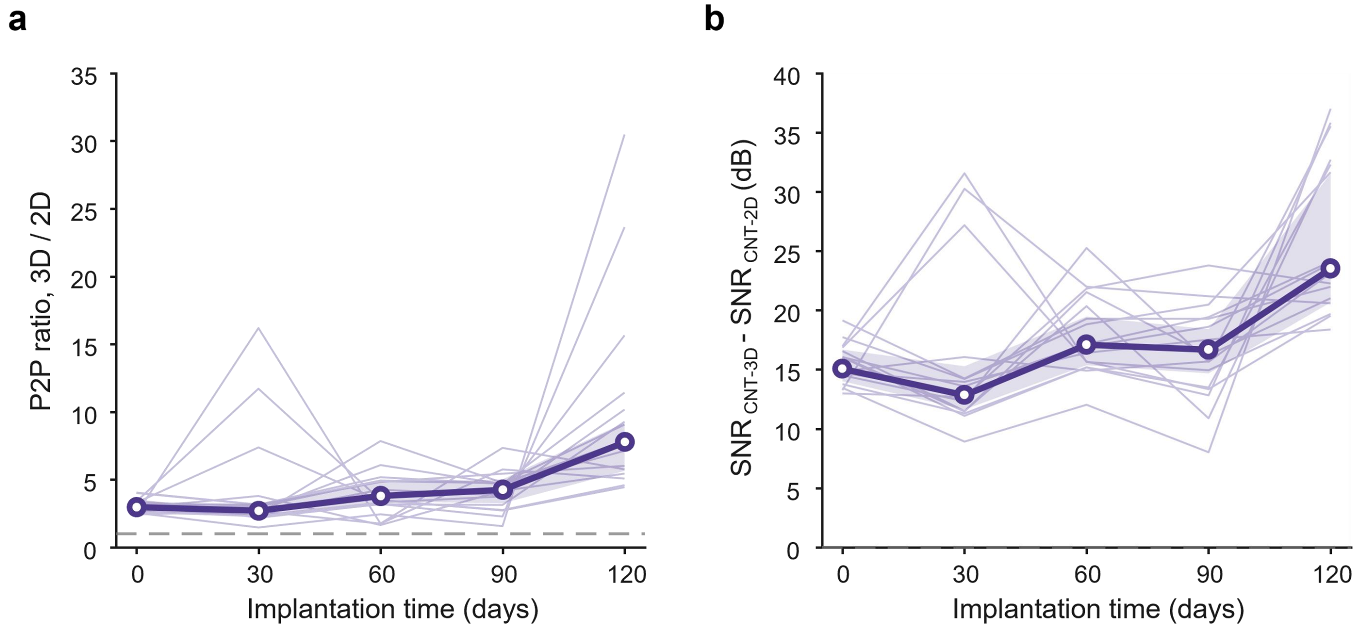


**Supplementary Fig. 35**. **Comparison of relative signal amplitude and SNR advantages of CNT-3D over CNT-2D arrays. a,** Time-course of the robust peak-to-peak amplitude ratio ($V_{pp,\ CNT\text{-}3D} / V_{pp,\ CNT\text{-}2D}$) over the 120-day implantation period. **b,** Time-course of the low-band $\Delta SNR$ ($SNR_{CNT\text{-}3D} - SNR_{CNT\text{-}2D}$).

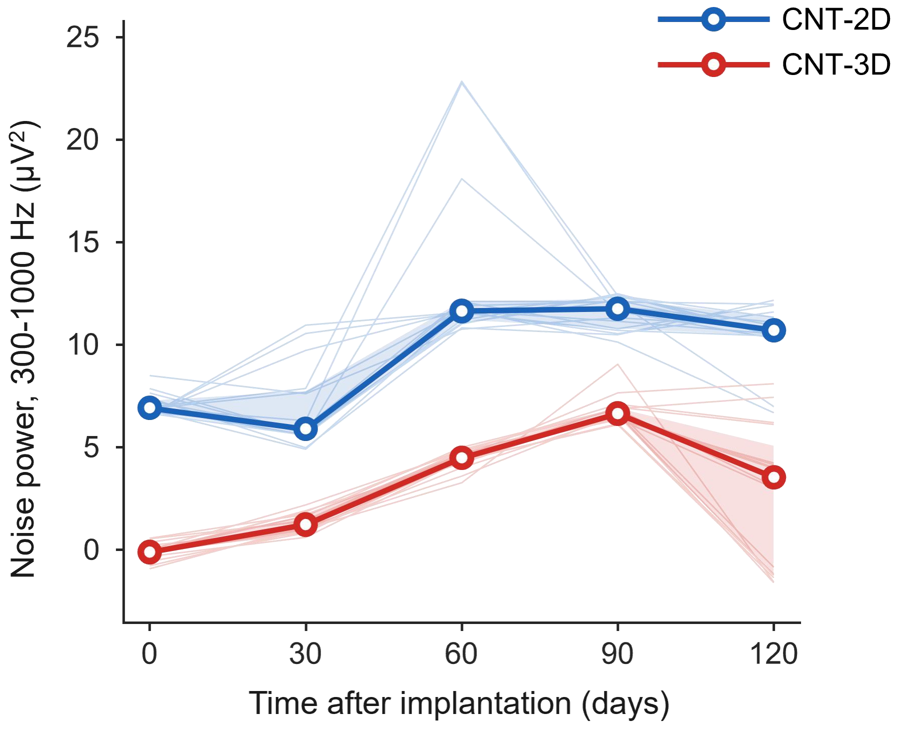


**Supplementary Fig. 36. Time-course of noise power in the 300-1000 Hz band for CNT-2D and CNT-3D arrays over the 120 days implantation period.**

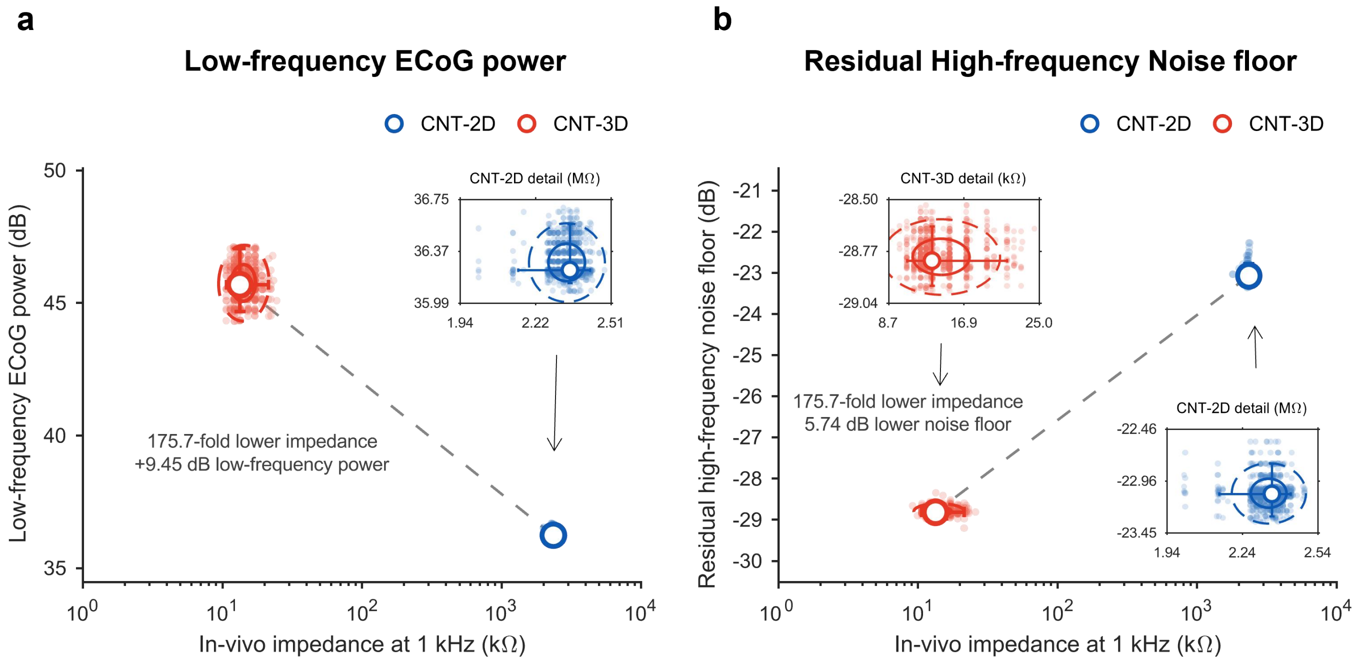


**Supplementary Fig. 37. Joint analysis of in vivo impedance and recording quality. a,** Relationship between low-frequency ECoG power and in vivo impedance at 1kHz for CNT-2D and CNT-3D arrays. **b,** Relationship between residual high-frequency noise floor and in vivo impedance at 1 kHz. This analysis complements Fig. 5j and shows that CNT-3D channels occupy a distinct operating regime characterized by lower impedance, higher low-frequency ECoG power and lower residual high-frequency noise.

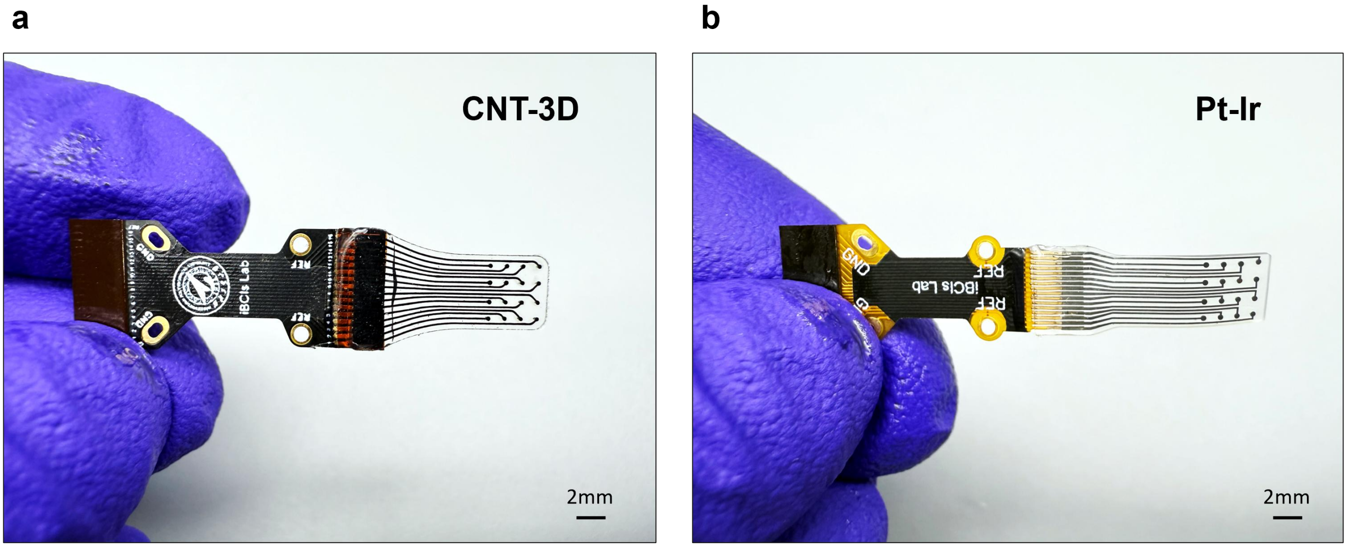


**Supplementary Fig. 38. Images of the CNT-3D and Pt-Ir array corresponding to the exploded-view schematics shown in Fig. 6a.** The two arrays were implanted over bilateral cortical regions in a New Zealand white rabbit to enable direct comparison of MRI images.

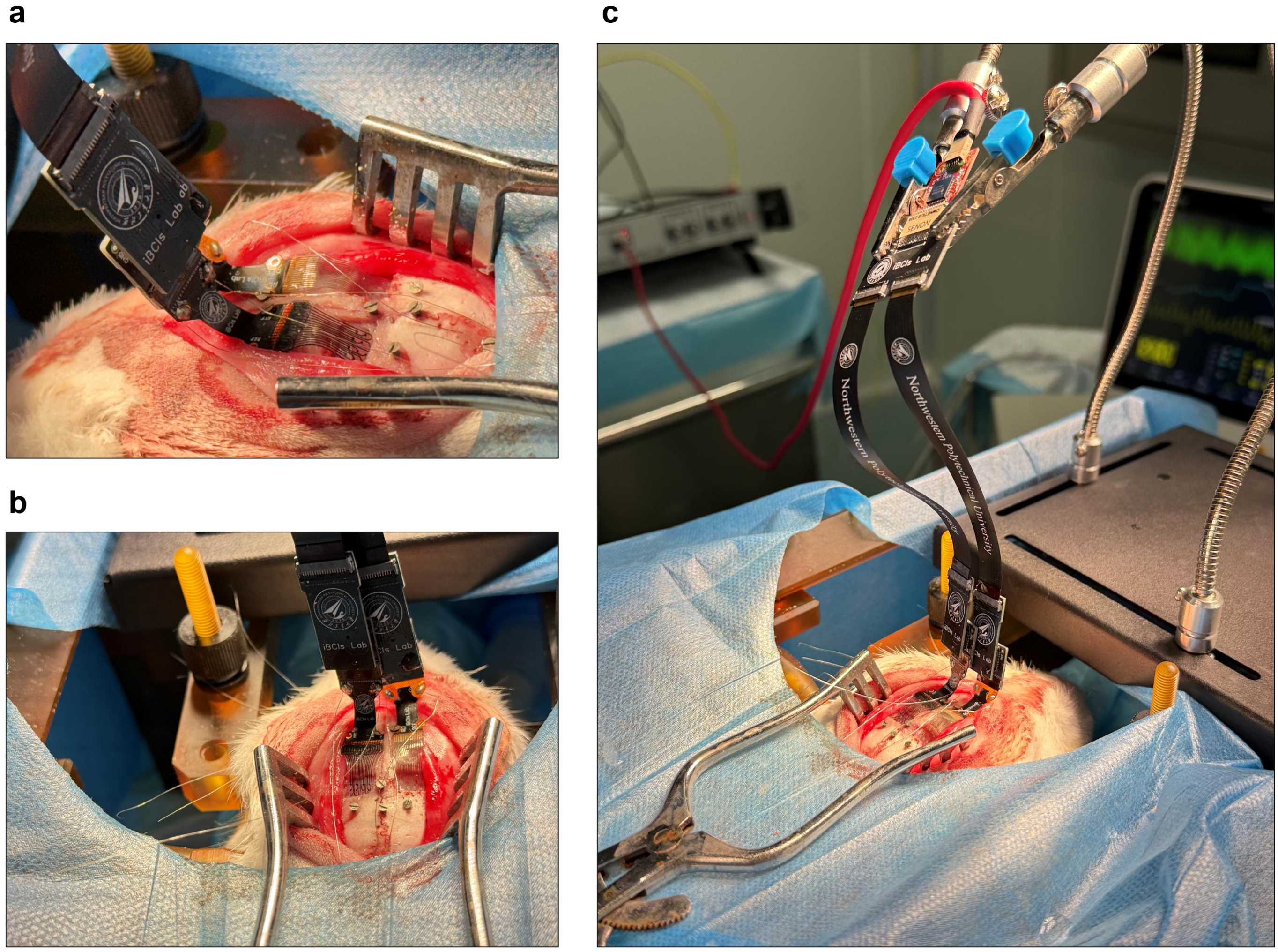


**Supplementary Fig. 39. Bilateral implantation of the CNT-3D and Pt-Ir arrays for MRI artefact evaluation. a,b,** Intraoperative side-view and top-view photographs showing bilateral implantation of the 16-channel CNT-3D array and the size-matched 16-channel Pt-Ir electrode. **c,** Photograph showing the implanted electrode arrays connected to the back-end Intan RHS 2000 system for electrical verification before MRI. This experimental configuration enabled direct in vivo comparison of MRI artefacts generated by the CNT-3D array and the conventional Pt-Ir electrode array under matched implantation conditions.

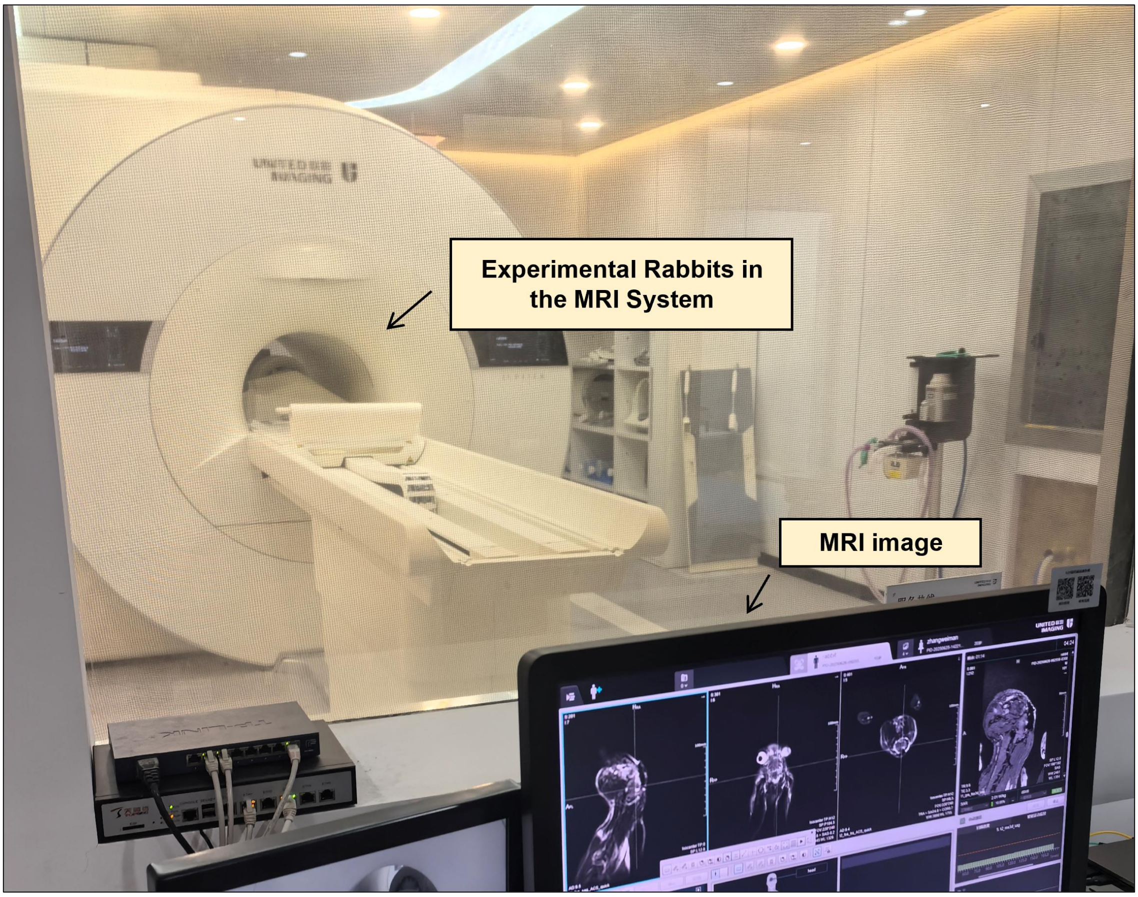


**Supplementary Fig. 40. In vivo MRI scanning setup for the implanted rabbit.** The MRI scan was performed after bilateral implantation of the CNT-3D and Pt-Ir arrays to compare implant-induced image artefacts under the same in vivo imaging conditions.

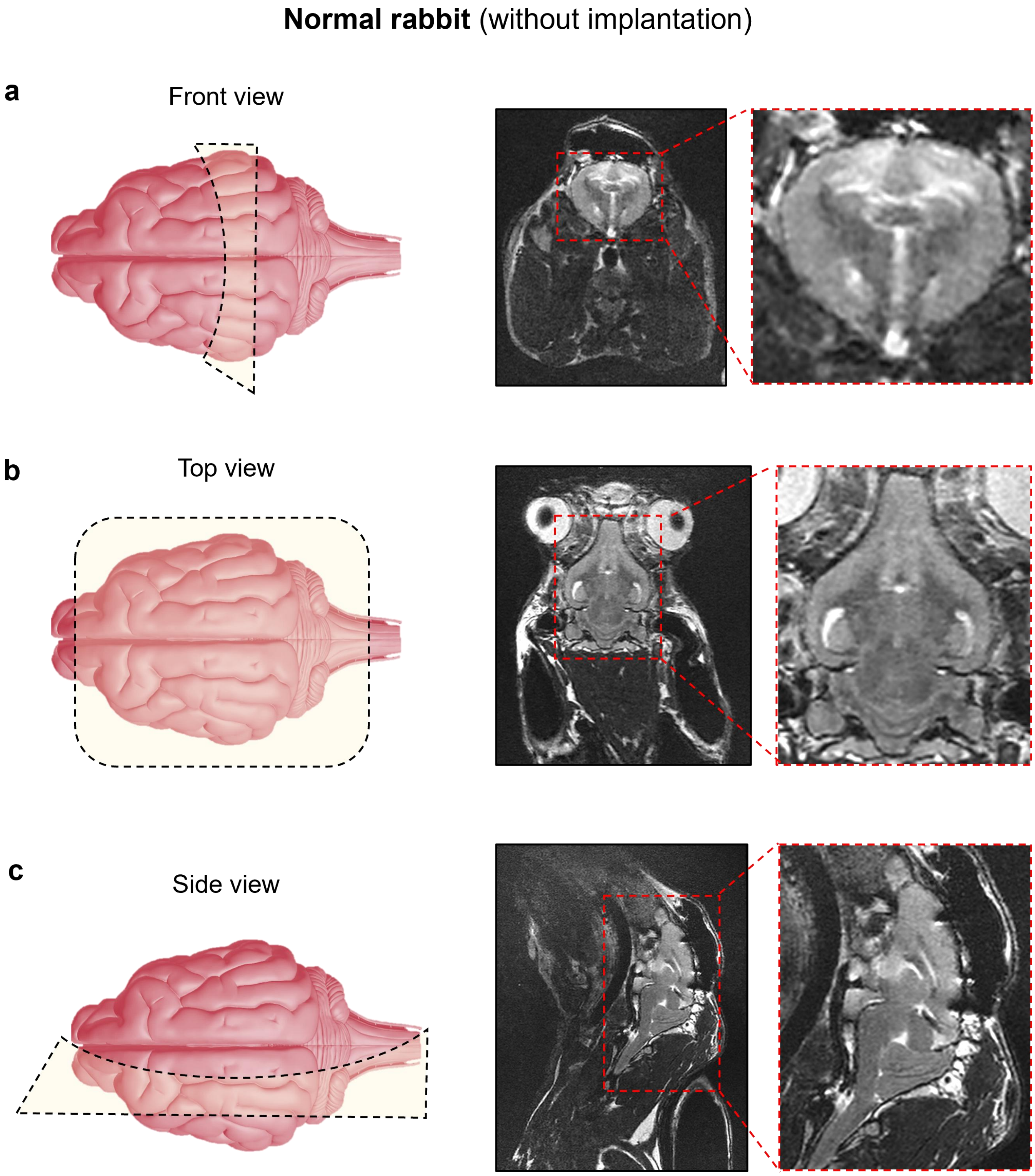


**Supplementary Fig. 41. The MRI images from a normal rabbit without implanted electrodes.** These brain MRI images serve as baseline anatomical references for evaluating implant-induced MRI artifacts.

a
b

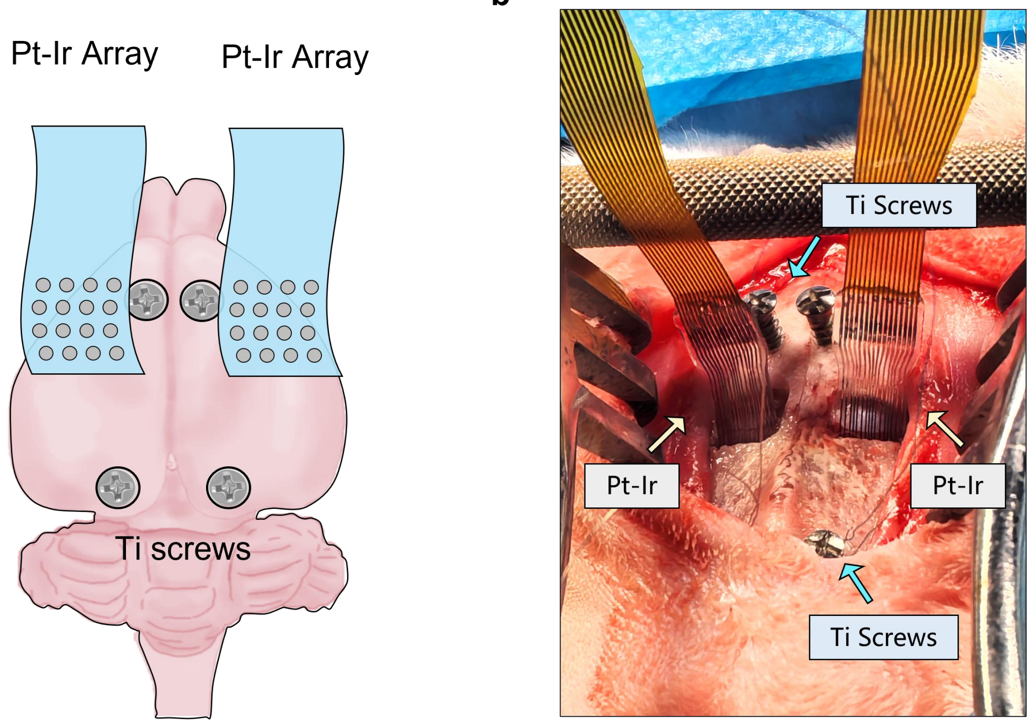


**Supplementary Fig. 42. Bilateral implantation of Pt-Ir arrays for MRI artifacts control experiments. a,** Schematic showing the implantation positions of two Pt-Ir arrays placed bilaterally over the rabbit cortex. **b,** Intraoperative photograph. This control configuration was used to evaluate MRI artifacts generated by conventional metallic electrodes.

#### **Control rabbit** (bilateral Pt-Ir array implantation)

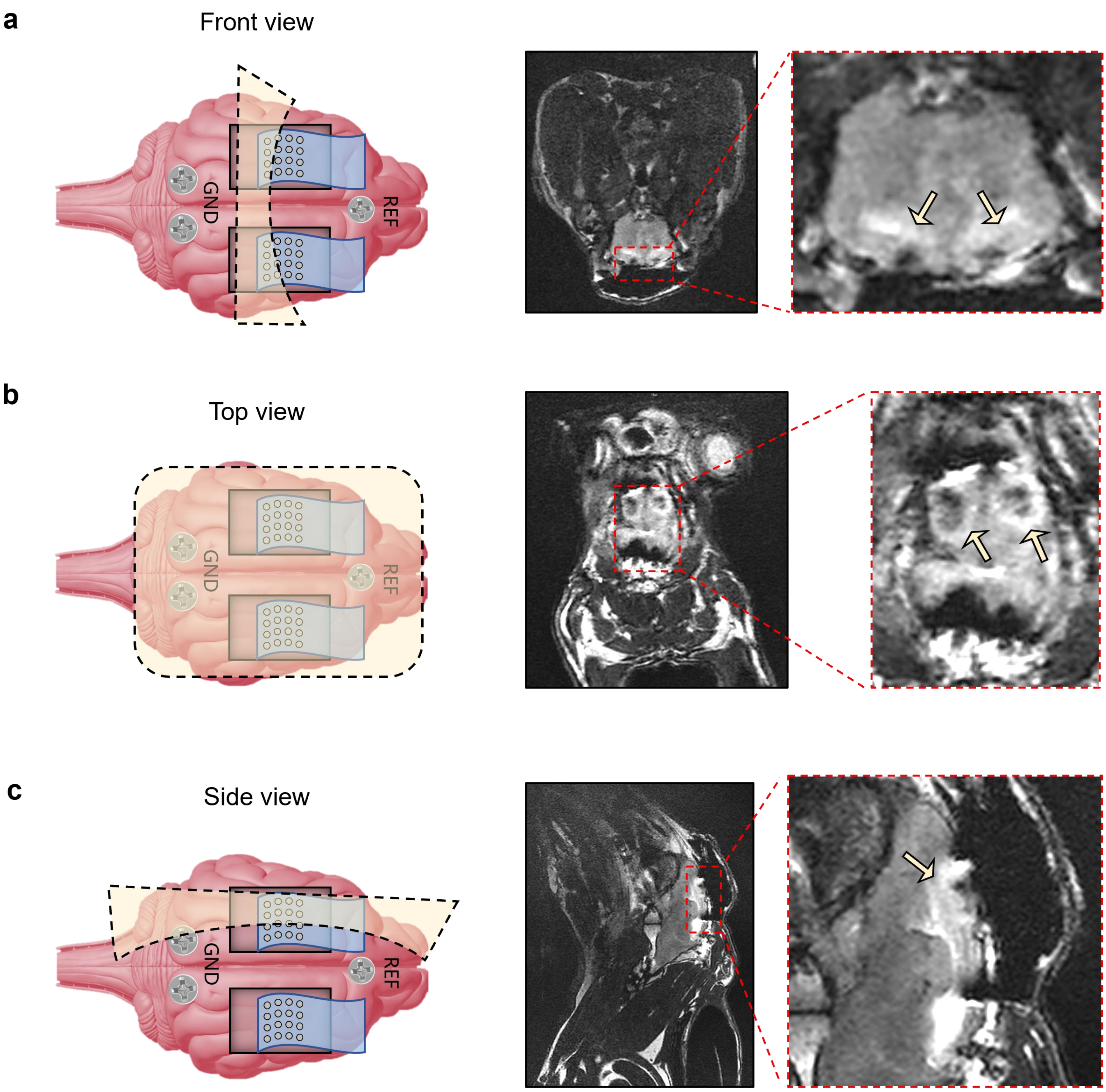


**Supplementary Fig. 43. Brain MRI images from control rabbits with bilateral Pt-Ir array implantation.** The bilateral Pt-Ir implants produced visible MRI artefacts around the implanted cortical regions, providing a metallic-electrode control for comparison with the CNT-3D array.

### Supplementary Tables

#### Supplementary Table 1. Qualitative benchmark of the ECoG electrode materials and architectures.

| Type | Material / architecture | $CSC_c$ / charge metric | \|Z\| at 1 kHz | Mechanical class / Young's modulus | MRI compatibility | Metal-free conductive path | Representative refs. |
|---|---|---|---|---|---|---|---|
| 2D Metal Electrodes | Au thin-film | Low to moderate, typically <1-3 mC $cm^{-2}$ | Moderate; strongly size-dependent, typically tens to hundreds of kΩ for micro sites | Usually stiff metal on flexible polymer | Generally usable but metal-dependent artifact risk | No | **[1, 2, 3]** |
| | Pt thin-film | Low to moderate; generally higher electrochemical stability than Au | Moderate | Stiff metal site on flexible substrate | Generally usable; artifact depends on geometry and scanner | No | **[1, 2, 4]** |
| | Pt-Ir / Pt composite | Low to moderate intrinsic CSC | Moderate to low when roughened | Stiff metal or metal-composite interface; substrate-dependent | Usually MRI-visible | No | **[3, 5, 23]** |
| 2D Coating-modified Electrodes | PEDOT:PSS on Au/Pt | High; often several to >10 fold higher than bare Au/Pt | Low to moderate; typically reduced by orders of magnitude vs bare small metal sites | Device mechanics usually dominated by polymer substrate and metal traces | Usually MRI-visible | Usually no, unless both sites and interconnects are PEDOT:PSS | **[1, 2, 9]** |
| | Pt black / nanoporous Pt | High apparent CSC | Low | Stiff metal nanostructured coating | Metal-dependent artifact risk | No | **[6, 24]** |
| | IrOx / SIROF | High CIC; widely used for stimulation | Low to moderate | Inorganic oxide coating on metal electrode | Metal-dependent artifact risk | No | **[5, 7]** |
| | TiN / conductive ceramic coating | High effective capacitance | Low to moderate | Stiff ceramic coating on flexible substrate | Metal-dependent artifact risk | No | **[5, 8]** |
| | Carbon black, CNT, graphene, rGO or MXene coating on metal | Moderate to high, depending porosity and coating thickness | Low to moderate | Full device usually metal/polymer hybrid | Carbon/MXene reduce optical/MRI issues, but metal traces remain | Usually no; carbon-coated metal is not fully metal-free | **[9–11]** |
| | Conductive hydrogel / PEDOT-carbon-fabric hydrogel | High interfacial capacitance | Lower than metal baseline in low frequency range | Soft, often kPa to MPa | Usually favorable if metal content is absent or minimal | Potentially yes, but interconnect must be verified | **[12, 13]** |
| 2D Carbon Electrodes | Graphene / multilayer graphene | Low to moderate | Moderate to high | Flexible thin-film; substrate typically PI, parylene, PET | Excellent optical compatibility; MRI compatibility often favorable | Usually no if Ti/Au contacts or metal traces are used | **[14–16]** |
| | rGO / nanoporous graphene / EGNITE | Moderate to high | Low to moderate | Flexible thin-film; substrate-dependent | Potentially favorable; must verify full device stack | Usually no unless full wiring is carbon-based | **[17, 18]** |
| | CNT or CNT-graphene film | Moderate to high | Low to moderate | Potentially soft if embedded in elastomer or free-standing film | Potentially favorable; depends on packaging and connectors | Possible, but only if sites, traces and pads are all carbon | **[19, 20]** |
| | Glassy carbon / laser-carbonized carbon | Moderate | Moderate | Stiff carbon film unless integrated with soft substrate | Generally favorable relative to metals | Usually not fully metal-free | **[21, 22]** |
| **3D Carbon Electrodes** | **CNT** | **6.48 mC $cm^{-2}$** | **8.6 kΩ** | **Soft elastomeric interface E ≈ 0.42 MPa** | **MRI-compatible (minimal artifact)** | **Yes, metal-free carbon sites, traces and pads** | **This work** |

**Footnote:** Values are summarized as representative ranges or qualitative envelopes. $CSC_c$, Z, CIC and areal capacitance are kept as reported by the original studies. "Metal-free" refers to the entire conductive pathway, including electrode sites, interconnects and contact pads.

#### Supplementary Table 2. Region-specific processing parameters for the laser drawing strategy.

| Patterned region | Functional purpose | Processing depth | Frequency | Pulse width | Scanning speed | Number of passes | Scan strategy | Hatch / line spacing |
|---|---|---|---|---|---|---|---|---|
| **1. Interconnect** | Define CNT-silicone interconnect trench | Through PVA only; underlying silicone retained | 50 kHz | 10 μs | 25 mm $s^{-1}$ | 2 | Cross-hatched raster filling | 0.02 mm |
| **2. Electrode site (3D)** | Form microcavity for protruding CNT-silicone electrode | Through PVA and silicone and into PMMA | 50 kHz | 2.5 μs | 25 mm $s^{-1}$ | 75 | Concentric circular filling | 0.015 mm |
| **3. Contact pad (3D)** | Form recessed/thickened conductive contact pad | Through PVA and silicone and shallowly into PMMA | 50 kHz | 2.5 μs | 100 mm $s^{-1}$ | 50 | Concentric circular filling | 0.03 mm |
| **4. Device outline** | Define final device boundary | Through the complete silicone device stack to PMMA | 50 kHz | 10 μs | 25 mm $s^{-1}$ | 10 | Contour scanning | N/A |

**Footnote:** All structures were patterned using a 355 nm UV nanosecond laser with a nominal maximum output power of 5 W. Region-specific penetration depth were controlled primarily by adjusting the pulse frequency, pulse width, scanning speed, number of passes and scan strategy.

**Supplementary Table 3. Benchmark comparison of fabrication strategies for minimally invasive ECoG electrodes.**

| No. | Benchmark metric | Photolithography-based MEMS / thin-film process | 3D printing / additive manufacturing | This method: depth-tunable laser drawing |
|---|---|---|---|---|
| 1 | **Soft silicone compatibility** | Moderate<br>Process-adapted silicone integration | Moderate-high<br>Ink/substrate dependent | High<br>Native /silicone-compatible process |
| 2 | **Photomask requirement** | Required | Not required | Not required |
| 3 | **Patterning resolution** | ~1–10 μm | ~10–100 μm typical | ~30 μm with ns UV laser<br>~10 μm possible with ps laser |
| 4 | **3D electrode geometry** | Limited (requires extra etching, plating, stacking or molding) | Strong (arbitrary 3D shanks, pillars or porous structures) | Cone diameter: 100-300 μm<br>Height: 0-300 μm |
| 5 | **3D structure type** | Mostly planar or quasi-planar | Usually penetrating 3D structures | Soft, non-penetrating carbon microcones |
| 6 | **Fabrication time** | Days to weeks | Few hours | Few hours |
| 7 | **Material cost per device** | High (typically dominated by cleanroom, metal and masks) | Variable (ink and printer dependent) | ~$1-3 per device, excluding connectors and labor |
| 8 | **Equipment burden** | High (cleanroom, aligner, deposition, etching) | Medium (aerosol jet / two-photon systems) | Low (UV laser, spin coater, oven) |
| 9 | **Process complexity** | Complex (typically 5-20+ microfabrication steps) | Moderate (printing plus curing, metallization, insulation) | Simple (laser patterning + carbon filling + encapsulation/release) |
| 10 | **Metal deposition / coating** | Usually required (Au, Pt, Pt-Ir, IrOx or PEDOT:PSS common) | Often required for conductivity or impedance reduction | Not required (CNT-silicone conductive network) |
| 11 | **Electrochemical tunability** | Mainly tuned by electrode area or coating | Tuned by printed geometry and material | Mainly tuned by electrode area and 3D micro geometry |
| 12 | **Reported stability** | High batch reproducibility | Variable (affected by ink, curing and post-processing) | High batch reproducibility |
| 13 | **Fabrication environment** | Cleanroom<br>Wet/dry microfabrication | Often non-cleanroom, but require sintering or post-processing | Common environment |
| 14 | **Main limitation** | Planar geometry, slow iteration, high fixed cost | Resolution-throughput trade-off, ink stability, packaging reliability | Lower resolution than MEMS |

**Footnote:** Values represent representative ranges rather than absolute process limits. Photolithography-based thin-film processes remain advantageous for high-resolution interconnects, multilayer alignment and reproducible soft ECoG manufacturing. Additive manufacturing provides stronger 3D geometric freedom but often requires conductive inks, metallization, insulation or post-processing. This depth-tunable laser drawing strategy combines maskless geometric programmability, soft silicone compatibility and metal-free carbon microcones, making it particularly suitable for non-penetrating close-contact neural interfaces.

**Supplementary Table 4. Specific dimensions of the representative 3D soft carbon electrode arrays shown in Fig. 2f.**

| Device / intended interface | Channels | Electrode layout | Electrode-site diameter | Nominal 3D height | Electrode pitch | Active-area dimensions | Overall device dimensions | Interconnect width |
|---|---|---|---|---|---|---|---|---|
| **Rabbit cortical array** | 16 | 4 × 4 | ~ 200 μm | ~ 180 μm | 1.2 mm | 5 × 5 $mm^2$ | 14.5 × 8.8 $mm^2$ | ~ 150 μm (Straight line) |
| **Primate cortical array** | 32 | 4 × 8 | ~ 600 μm | ~ 100 μm | 3.1 mm | 22.3 × 10 $mm^2$ | 32.3 × 14 $mm^2$ | ~ 150 μm (Straight line) |
| | 64 | 8 × 8 | ~ 600 μm | ~ 100 μm | 3.1 mm | 22.3 × 22.3 $mm^2$ | 32.3 × 25.6 $mm^2$ | ~ 150 μm (Straight line) |
| **Human-scale cortical array** | 32 | 4 × 8 | ~ 2 mm | 0 μm (Flat with the encapsulation layer) | 1 cm | 73.6 × 73.6 $mm^2$ | 81 × 87 $mm^2$ | ~ 100 μm (serpentine line) |
| | 64 | 8 × 8 | ~ 2 mm | 0 μm (Flat with the encapsulation layer) | 1 cm | 73.6 × 73.6 $mm^2$ | 81 × 87 $mm^2$ | ~ 100 μm (serpentine line) |
| **Spinal cord array** | 16 | Staggered arrangement (5 + 6 + 5) | ~ 4 × 1.5 $mm^2$ | 0 μm (Flat with the encapsulation layer) | 8 mm (Horizontal) 2.5 mm (Vertical) | 44.5 × 6.5 $mm^2$ | 55.9 × 10.7 $mm^2$ | ~ 150 μm (Straight line) |

### References


1. Ganji, M. et al. Scaling effects on the electrochemical performance of poly(3,4-ethylenedioxythiophene) (PEDOT), Au, and Pt for electrocorticography recording. *Adv. Funct. Mater.* **27**, 1703018 (2017).
2. Ganji, M., Tanaka, A., Gilja, V., Halgren, E. & Dayeh, S. A. Scaling effects on the electrochemical stimulation performance of Au, Pt, and PEDOT:PSS electrocorticography arrays. *Adv. Funct. Mater.* **27**, 1703019 (2017).
3. Fallegger, F. et al. MRI-compatible and conformal electrocorticography grids for translational research. *Adv. Sci.* **8**, 2003761 (2021).
4. Tchoe, Y. et al. Human brain mapping with multithousand-channel PtNRGrids resolves spatiotemporal dynamics. *Sci. Transl. Med.* **14**, eabj1441 (2022).
5. Cogan, S. F. Neural stimulation and recording electrodes. *Annu. Rev. Biomed. Eng.* **10**, 275-309 (2008).
6. Lee, J. Y. et al. Foldable three dimensional neural electrode arrays for simultaneous brain interfacing of cortical surface and intracortical multilayers. *npj Flex. Electron.* **6**, 86 (2022).
7. Cogan, S. F. et al. Sputtered iridium oxide films for neural stimulation electrodes. *J. Biomed. Mater. Res. B Appl. Biomater.* **89B**, 353-361 (2009).
8. Rodrigues, F. et al. Fabrication and characterization of polyimide-based 'smooth' titanium nitride microelectrode arrays for neural stimulation and recording. *J. Neural Eng.* **17**, 016010 (2020).
9. Castagnola, E. et al. PEDOT-CNT-coated low-impedance, ultra-flexible, and brain-conformable micro-ECoG arrays. *IEEE Trans. Neural Syst. Rehabil. Eng.* **23**, 342-350 (2015).
10. Garg, R. et al. Graphene and poly(3,4-ethylenedioxythiophene)–polystyrene sulfonate hybrid nanostructures for input/output bioelectronics. *ACS Appl. Nano Mater.* **6**, 8495-8505 (2023).
11. Driscoll, N. et al. MXene-infused bioelectronic interfaces for multiscale electrophysiology and stimulation. *Sci. Transl. Med.* **13**, eabf8629 (2021).
12. Oribe, S. et al. Hydrogel-based organic subdural electrode with high conformability to brain surface. *Sci. Rep.* **9**, 13379 (2019).
13. Tringides, C. M. et al. Viscoelastic surface electrode arrays to interface with viscoelastic tissues. *Nat. Nanotechnol.* **16**, 1019-1029 (2021).
14. Park, D.-W. et al. Graphene-based carbon-layered electrode array technology for neural imaging and optogenetic applications. *Nat. Commun.* **5**, 5258 (2014).
15. Bakhshaee Babaroud, N. et al. Multilayer CVD graphene electrodes using a transfer-free process for the next generation of optically transparent and MRI-compatible neural interfaces. *Microsyst. Nanoeng.* **8**, 107 (2022).
16. Garcia-Cortadella, R. et al. Graphene active sensor arrays for long-term and wireless mapping of wide frequency band epicortical brain activity. *Nat. Commun.* **12**, 211 (2021).

17. Lu, Y., Lyu, H., Richardson, A. G., Lucas, T. H. & Kuzum, D. Flexible neural electrode array based-on porous graphene for cortical microstimulation and sensing. *Sci. Rep.* **6**, 33526 (2016).
18. Viana, D. et al. Nanoporous graphene-based thin-film microelectrodes for in vivo high-resolution neural recording and stimulation. *Nat. Nanotechnol.* **19**, 514-523 (2024).
19. Chen, Y.-C. et al. An active, flexible carbon nanotube microelectrode array for recording electrocorticograms. *J. Neural Eng.* **8**, 034001 (2011).
20. Wei, S. et al. Shape-changing electrode array for minimally invasive large-scale intracranial brain activity mapping. *Nat. Commun.* **15**, 715 (2024).
21. Vomero, M. et al. Highly stable glassy carbon interfaces for long-term neural stimulation and low-noise recording of brain activity. *Sci. Rep.* **7**, 40332 (2017).
22. Vomero, M., Oliveira, A., Ashouri, D., Eickenscheidt, M. & Stieglitz, T. Graphitic carbon electrodes on flexible substrate for neural applications entirely fabricated using infrared nanosecond laser technology. *Sci. Rep.* **8**, 14749 (2018).
23. Lacour, S. P., Courtine, G. & Guck, J. Materials and technologies for soft implantable neuroprostheses. *Nat. Rev. Mater.* **1**, 16063 (2016).
24. Lim, J., Zoss, P. A., Powley, T. L., Lee, H. & Ward, M. P. A flexible, thin-film microchannel electrode array device for selective subdiaphragmatic vagus nerve recording. *Microsyst. Nanoeng.* **10**, 16 (2024).
